# Multispectral live-cell imaging with uncompromised spatiotemporal resolution

**DOI:** 10.1101/2024.06.12.597784

**Authors:** Akaash Kumar, Kerrie E. McNally, Yuexuan Zhang, Alex Haslett-Saunders, Xinru Wang, Jordi Guillem-Marti, David Lee, Buwei Huang, Sjoerd Stallinga, Robert R. Kay, David Baker, Emmanuel Derivery, James D. Manton

## Abstract

Multispectral imaging is an established method to extend the number of colours usable in fluorescence imaging beyond the typical limit of three or four, but standard approaches are poorly suited to live-cell imaging. We introduce an approach for multispectral imaging in live cells, comprising an iterative spectral unmixing algorithm and eight channel camera-based image acquisition hardware. This enables the accurate unmixing of low signal-to-noise ratio datasets, typical of live-cell imaging, captured at video rates. We use this approach on a commercial spinning disk confocal microscope and a home-built oblique plane light sheet microscope to image 1–7 spectrally distinct fluorophore species simultaneously, using both fluorescent protein fusions and small-molecule dyes. We further use de novo designed protein-binding proteins (minibinders), labelled with organic fluorophores, and use these in combination with our multispectral imaging approach to study the endosomal trafficking of cell-surface receptors at endogenous levels.

## Introduction

Living cells contain thousands of different components, each with their own specific interaction partners and complex dynamics. Fluorescence microscopy has proven to be a powerful tool for studying the dynamics of life as it provides background-free images of cells and tissues with molecular specificity. However, the broad spectra of typical fluorophores compatible with biological imaging limit the number of colours that can be employed without spectral overlap and cross-talk between colour channels. Hence, typically only two or three cellular components can be studied in the same sample.

Multispectral imaging with spectral unmixing provides a way around this limitation, by computationally assigning light captured in discrete wavebands to different fluorophores [1]. However, conventional implementations of spectral unmixing [2–9] suffer from three key problems that have limited their use, particularly for live-cell imaging:

1. light is chromatically separated into a large number of wavebands (*e*.*g*. 32), leading to low signal levels in any one band;
2. illumination is provided as a spot or a line which must be scanned across the field of view, leading to slow acquisition rates and high irradiances;
3. linear spectral unmixing algorithms deal poorly with noise, leading to inaccurately reconstructed data.

Recently, Valm et al. introduced a light-sheet-based multispectral imaging system, which used sequential excitations of differing wavelengths in combination with camera-based readout [10]. This system obviates the first two problems as all the emission light is collected on the same detector and an entire plane is illuminated at once. However, the need to sequentially excite the sample with different wavelengths reduces imaging speed by a factor at least as large as the number of channels. In addition, the spectral unmixing was carried out using the standard linear matrix inversion approach [11], requiring that high signal-to-noise ratio (SNR) data must be acquired for accurate results and hence further increasing the illumination dose. Overall, the system was capable of imaging a full cell in six channels every 9.2 s for 100 time-points.

Here, we present an optimised approach for simultaneous multispectral acquisition and subsequent offline unmixing of fluorescence microscopy data, ideally suited for imaging live samples with minimal phototoxicity. First, we establish an iterative spectral unmixing algorithm and show how its improved performance over the state-of-the-art allows the successful unmixing of low SNR live-cell data. We then describe a cost-effective hardware module that provides any camera-based fluorescence microscope with 8-channel multispectral imaging capabilities with no penalty in spatiotemporal resolution and with 100 % photon efficiency. We then show how these two developments can be combined to achieve multispectral imaging of live cells in 2D+*t &* 3D+*t* on both spinning disk and light sheet microscopes, providing full cell volumes in as little as 0.3 s or full cell projections in as little as 0.1 s. Finally, we capitalise on the combination of our imaging technology with de-novo designed protein binders to directly observe endosomal sorting of endogenous trans-membrane receptors in living cells.

## Results

### Spectral unmixing of low SNR data

First, we investigated whether traditional spectral unmixing approaches were suitable for application to live-cell imaging datasets. We simulated multispectral datasets by computationally mixing fluorescence from eight fluorophores into eight spectral channels (Figure 1a) and adding Poisson (*i*.*e*. shot) noise (Figure 1b). Unmixing these data using conventional linear unmixing (*i*.*e*. applying the inverse of the mixing matrix to the mixed signals) showed that this produced poor results (Figure 1c), with more errors associated with unmixing lower SNR data (see Supplementary Figure 1a-d). In particular, linear unmixing predicted negative quantities of targets in many pixels (shown in red), which is physically impossible. This occurs because linear unmixing is incapable of dealing with Poisson (shot) noise, the predominant form of noise present in low SNR datasets, such as those generated by live-cell imaging.

**Figure 1:**
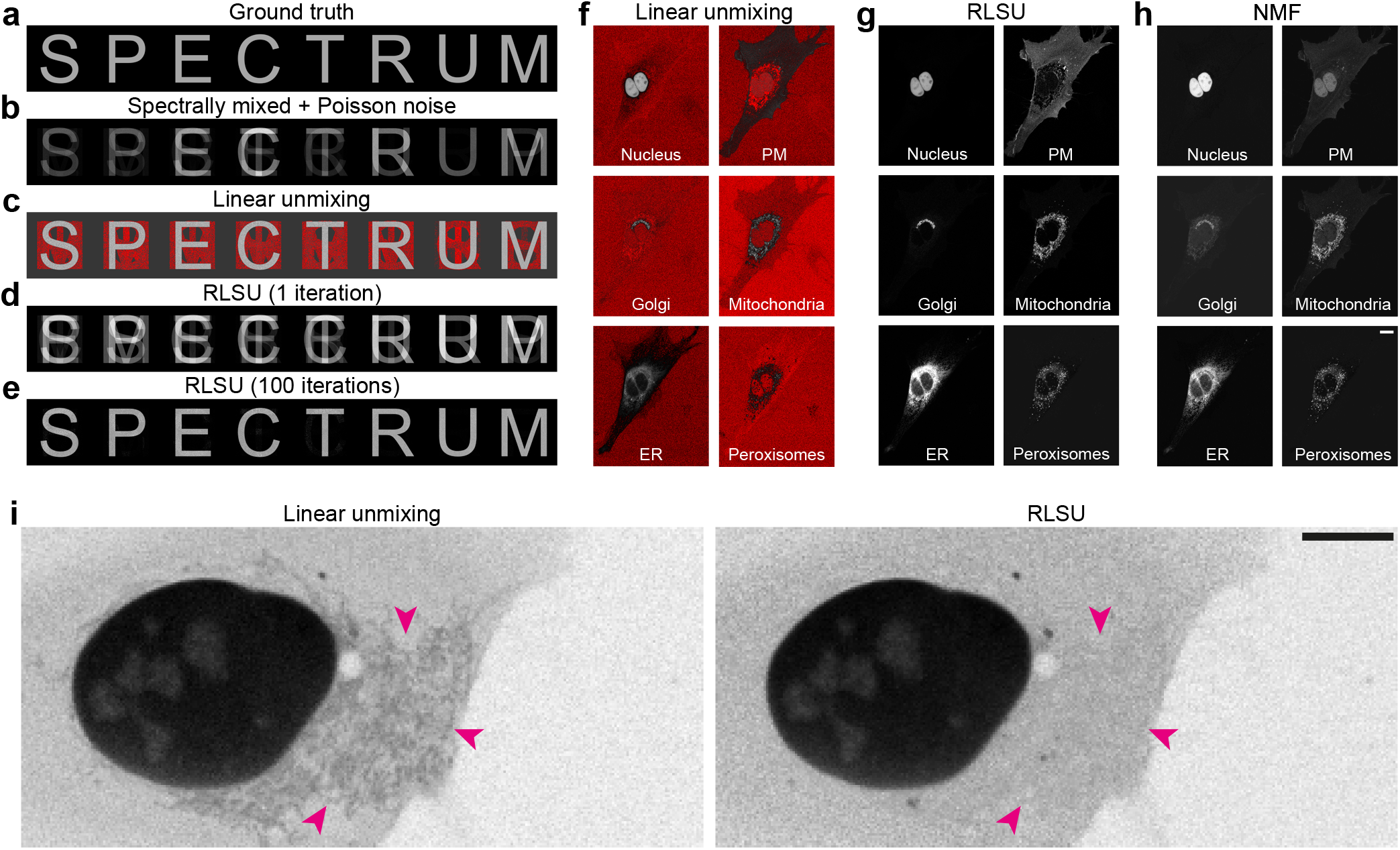
Richardson–Lucy spectral unmixing (RLSU) outperforms linear unmixing and non-negative matrix factorisation. **a** Simulated ground truth data of eight objects (letters of the word SPECTRUM) labelled with different fluorophores. **b** Simulated acquired data, spectrally mixed and including Poisson (shot) noise. **c** Reconstructed objects using data from (b) and linear unmixing. Negative values are shown in red. **d** Reconstructed objects using data from (b) and one iteration of RLSU. At this stage the algorithm has not converged on a solution. **e** Reconstructed objects after convergence using data from (b) and 100 iterations of RLSU. **f** Linearly unmixed objects from an experimentally-acquired dataset of a U2OS cell co-expressing six different fluorescent protein species. Negative values are shown in red. **g** RLSU unmixed objects from the same dataset in (f). **h** Non-negative matrix factorisation (NMF) unmixed objects from the same dataset in (f). Note that signals have not been correctly assigned to objects (*e*.*g*. nuclear signal in the plasma membrane channel). **i** Comparison of linear unmixing and RLSU for another real dataset in which mitochondrial signal bleeds through to the nuclear channel after linear unmixing, but is correctly absent after RLSU. Scale bars = 10 μm.

We reasoned that using an iterative approach, tailored to incorporate the effects of Poisson (shot) noise, would provide superior results. In particular, we elected to repurpose the Richardson–Lucy algorithm (RL) [12, 13] as this is fast, straightforward to use, and has been well-studied over the five decades since its development. While RL has previously been used for deconvolving images using a known point spread function (PSF), the algorithm is, in principle, applicable to any system where the measurement process can be described by a linear operator and the noise is described by Poisson statistics. Various researchers have used Richardson–Lucy deconvolution to enhance the linewidths of spectral profiles, but these algorithms do not separate the different components as required for spectral unmixing [14–16]. Instead, here we replace the action of convolving with the PSF with the action of mixing with the mixing matrix, which describes the contribution of each underlying object to each channel (see Supplementary Note A for more details). Hence, this new spectral unmixing algorithm does not deconvolve the data and so does not suffer from the high-spatial-frequency artefacts often seen after deconvolution. However, our approach can be extended to both unmix and deconvolve the data simultaneously (see Supplementary Note B and Supplementary Figure 2) but, for simplicity, we do not use this in this report.

Applying this Richardson–Lucy spectral unmixing algorithm (RLSU) to the same simulated dataset shows that while one iteration is insufficient to accurately unmix the different components (Figure 1d), 100 iterations produces a much higher quality result than linear unmixing (Figure 1e). In our hands, RLSU also outperformed a recent implementation of phasor-based unmixing, HyU [9], for both eight-channel (Supplementary Figure 3) and 32-channel data (Supplementary Figure 4). Noticeably, no negative quantities are generated by RLSU — indeed, this is impossible if the initial estimate of the components is everywhere positive, as the iterative update is multiplicative in nature. We discuss the appropriate Cramér–Rao lower bound in Supplementary Note C and show that we reach it in Supplementary Figure 5. Repeating this process for different SNR levels, and incoporating both Poisson (shot) noise and read noise, we found our iterative algorithm always outperformed linear unmixing (see Supplementary Figure 1, Supplementary Figure 6, Supplementary Figure 7 and Supplementary Video 1). Importantly, read noise only distorts the quality of unmixing results when the variance of the read noise is comparable to the variance of the Poisson (shot) noise. For modern sensors for which the read noise is small, such as the 2.32 e^*−*^ read noise of the cameras used in this work, this means that the effects of read noise are only significant for signal levels below 5 counts. We also found that RLSU outperforms linear unmixing for highly overlapping spectral signals, such as EGFP and EYFP (20 nm peak-to-peak separation), and can unmix signals with a peak-to-peak spectral separation as low as 4 nm using only two channels (see Supplementary Note D and Supplementary Figure 8).

Encouraged by these results, we applied our algorithm to data acquired on a commercially available multispectral confocal microscope (Zeiss LSM 710 with 32-channel QUASAR detector). U2OS cells transfected with a polycistronic plasmid, ColorfulCell [17], encoding six fluorescent protein species targetted to the nucleus (TagBFP), plasma membrane (PM) (Cerulean), mitochondria (mAzamiGreen), Golgi apparatus (Citrine), endoplasmic reticulum (ER) (mCherry) or peroxisomes (iRFP670) were imaged live, with data exhibiting SNRs up to 13 (Supplementary Figure 9). We then attempted to unmix the live-cell data using both RLSU and linear unmixing.

Linear unmixing did not always accurately reassign signals to the correct labelled structures, such as the Golgi signal incorrectly present in the mitochondria channel in Figure 1f, and once again produced numerous pixels with unphysical negative values. In contrast, RLSU produced unmixed objects that accurately resembled the single-labelled controls (Figure 1g), with no observable bleedthrough or misassignment of signals. A comparison with an implementation of non-negative matrix factorisation (NMF), a blind spectral unmixing approach also designed to handle Poisson (shot) noise but without requiring a priori knowledge of the mixing matrix [18], showed that while this did not produce negative quantities, none of the signals were correctly assigned (Figure 1h). Most strikingly, significant nuclear signal was present in the plasma membrane channel.

Applying both linear unmixing and RLSU on various cell images of differing signal levels, we found that in all cases RLSU was more robust against channel misassignment. Figure 1i shows an example of this, where a weak mitochondrial signal has been assigned to the nuclear channel by linear unmixing, but is accurately absent in the RLSU results. In both cases, the weak background signal outside the nucleus is due to cytosolic fluorescent proteins that have not yet been redirected to the nuclear compartment by the fused targetting signal. Furthermore, we tested the sensitivity of RLSU to small errors in the mixing matrix by unmixing simulated ColorfulCell data with a mixing matrix formed by an incorrect combination of fluorophores (see Supplementary Figure 10). Through this, we found we could, for example, replace mCherry’s mixing matrix values with those for Alexa Fluor 594 with no penalty in the quality of the unmixing results, despite their slightly different spectral profiles. However, mScarlet’s more shifted spectrum led to noticeable reconstruction errors when the mixing matrix was updated to use its values.

### Acquiring video-rate spectral information

Given the performance of RLSU on low-SNR data, we sought to develop imaging hardware that would allow us to capture the necessary raw signals at rates much faster than those afforded by the point-scanning QUASAR detector. In particular, we decided that the hardware should have five key characteristics and should:

1. be compatible with any camera-based microscope, by replacing the native camera;
2. enable multispectral data capture at the original rate of the parent instrument, thereby maintaining temporal resolution;
3. be diffraction-limited, thereby maintaining the spatial resolution of the parent instrument;
4. be maximally photon-efficient, thereby maximising SNR;
5. be easily reconfigurable to best match the spectral characters of the fluorophores used in an experiment.

These considerations led us to design a system using dichroic mirrors, in a tree-like arrangement, to redirect light to multiple cameras (see optical path illustration in Figure 2a and hardware photo in b). In contrast to conventional multispectral approaches using diffraction gratings or interferometers, this enables the capture of the full spatial and spectral information in a single shot. Furthermore, as light is merely redirected by the dichroic mirrors, not absorbed as by a filter, there is no loss of photons through the tree-like arrangement — a photon that does not end up in a specified channel must end up in one of the others. While this does not obviate the absorbative losses in the extra lenses used, these are small (∼3 %). We settled on using seven dichroics (spectra plotted in Figure 2c) to provide eight channels of data (spectra plotted in Figure 2d), as this was the most effective choice for imaging over the typical 450 nm to 700 nm spectral range used in live-cell fluorescence microscopy experiments (see Supplementary Note E). In the interests of portability, rejection of excitation light is achieved using the primary dichroic, and if necessary a notch filter, in the parent instrument.

**Figure 2:**
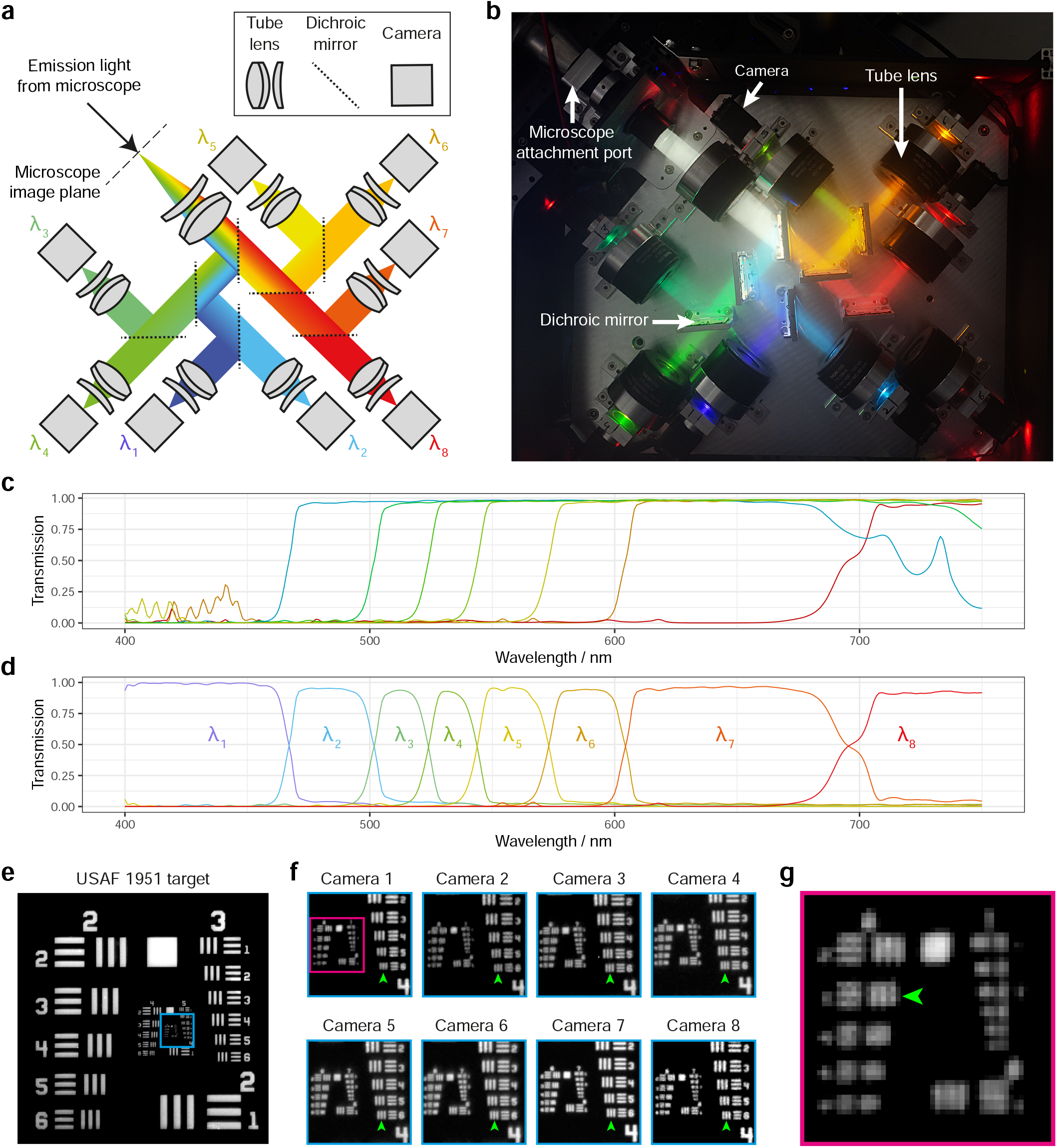
Multispectral acquisition module compatible with camera-based fluorescence microscopes. **a** Optical path illustration for the multispectral imaging module. Light enters from the top left and is redirected, based on wavelength, onto eight cameras by seven dichroic mirrors. **b** Photograph of assembled multispectral imaging module. **c** Transmission spectra of the seven dichroic mirrors used. **d** Spectra of the eight channels arising from the spectra in (c). **e** USAF 1951 target imaged in the most blue channel. **f** Enlarged view of target region indicated by cyan box in (e), shown for all eight channels. Green arrows indicate finest resolvable group across all channels. **g** Enlarged view of target region indicated by magenta box in (f), shown for channel 1 (green arrow indicates finest resolvable group in channel 1).

Raytracing showed that our design would maintain diffraction-limited resolution over the full field of view of the tube lenses used (20 mm, 0), even on the path where the light traverses three tilted dichroic mirrors (Supplementary Figure 11). To enable future extension to 16 channels, extending the spectral range into the near infrared, we also validated that diffraction-limited resolution was maintained in the case that four tilted dichroics were used. Testing the resolution of the assembled system using a USAF 1951 target (Figure 2e) showed that the resolving power of the system was limited by the Nyquist sampling rate (6.21 μm/px, see Supplementary Note F). This sampling rate was chosen to match that of a typical sCMOS camera, as commonly used with high-resolution fluorescence microscopes (6.5 μm/px).

By design, the sampling rate of the system can be changed by altering the focal length ratio of the input and output lenses. As constructed, we used an input lens with 180 mm focal length, and output lenses with focal lengths of 100 mm. Alternative tube lenses with focal lengths from *f* = 165 mm to 600 mm are also available, giving effective pixel sizes spanning 3.45 μm to 20.7 μm. Further adjustments to the effective pixel size would either require replacing the output lenses or selecting different camera sensors.

In order to ensure maximum mechanical stability, we elected to not mount the dichroic mirrors kinematically. As a result, each image is slightly displaced off-axis due to small angular deviations of the dichroic mirror orientation from the design specification. However, these shifts were sufficiently small that they could be corrected by a simple image registration routine and left *>* 80 % of the field of view usable (Supplementary Figure 12).

### Multispectral spinning disk confocal microscopy

First, we coupled our multispectral module to a spinning disk confocal microscope (SDCM). Point spread function measurements showed resolutions equivalent to those obtained without the multispectral module, demonstrating that the module did not compromise spatial resolution (Supplementary Figure 13). Imaging U2OS cells transfected with the six-colour ColorfulCell plasmid, and stained with LysoTracker Yellow, produced eight camera images spanning the visible spectrum (Figure 3a). To facilitate the choice of appropriate fluorophores and generation of the corresponding mixing matrix, we developed a web application, hosted at beryl.mrc-lmb.cam.ac.uk/calculators/ spectral_unmixing/. This uses manufacturer-supplied spectra for the dichroics used in the module, as well as the dichroic and notch filter in the spinning disk microscope, in addition to reference fluorescence emission spectra from fpbase.org, to enable the calculation of appropriate mixing matrices and provide feedback on whether the experiment is spectrally feasible (Supplementary Figure 14). Using a mixing matrix generated by the web application from an appropriate selection of fluorophores (Figure 3b), we unmixed the camera images to form seven object channels (Figure 3c).

**Figure 3:**
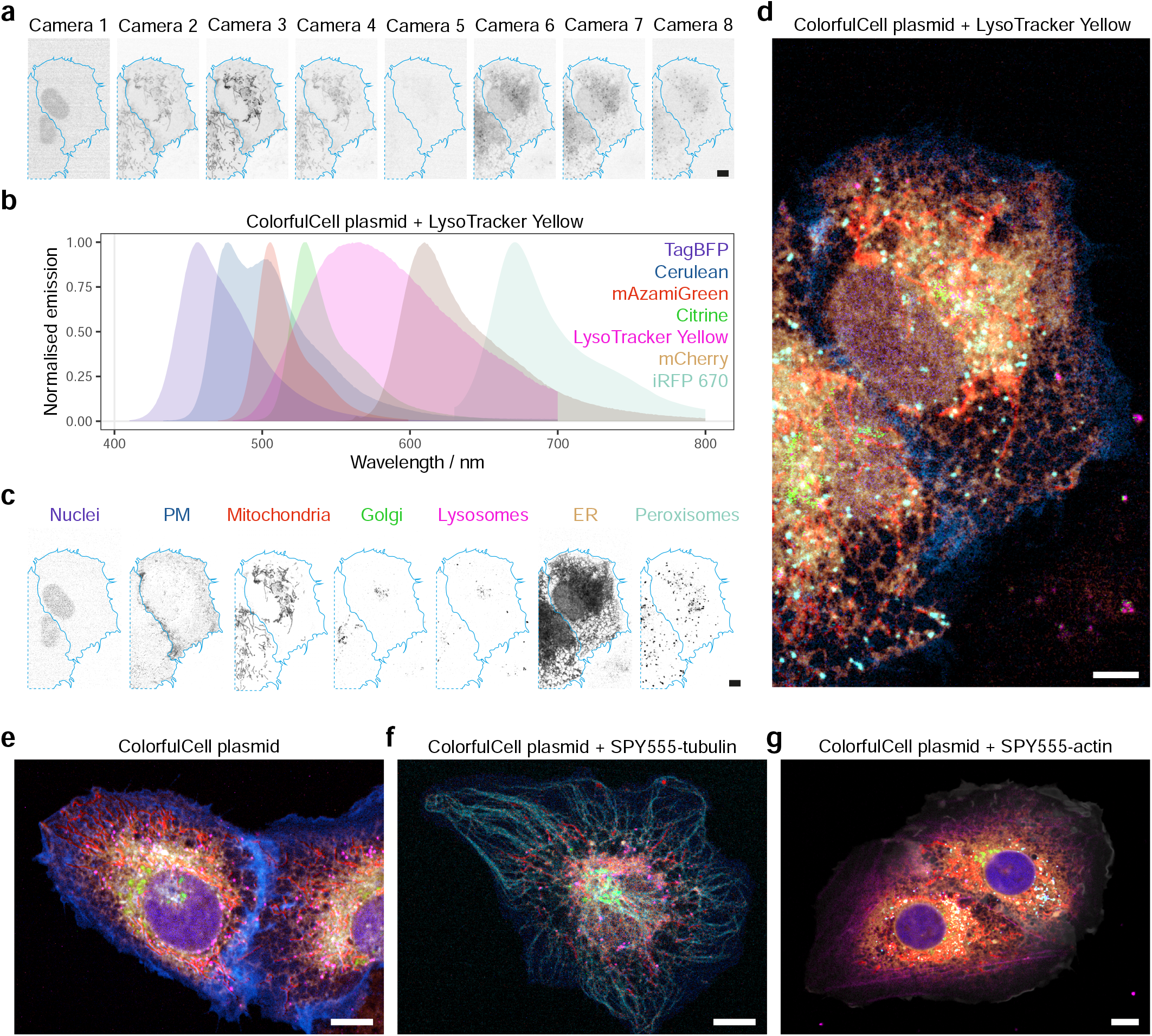
Multispectral live-cell imaging on a spinning disk confocal microscope. **a-d** U2OS cells transfected with the ColorfulCell plasmid were incubated with LysoTracker Yellow and imaged live on a spinning disk confocal instrument equipped with the multispectral camera system. The eight individual camera views (**a**) were acquired simultaneously, then processed by RLSU umixing using a mixing matrix generated from the emission spectra of seven fluorophores (**b**) to generate unmixed images for the seven fluorophores (**c**). **d** Colour merge of channels in (c). **e-g** Other examples of RLSU unmixing results for multispectral spinning disk confocal imaging of U2OS cells transfected with ColorfulCell plasmid (**e**), or additionally incubated with SPY555-tubulin (**f**) or SPY555-actin (**g**) dyes. Scale bars = 10 μm.

Despite the four-fold smaller number of spectral channels compared to the QUASAR detector, we obtained unmixed images that accurately reflected the expected distributions of fluorescent proteins in their respective organelles (*cf*. Supplementary Figure 15). As the full spatial and spectral information were acquired in one exposure time, we found that imaging speeds could be increased more than 100-fold over our previous confocal QUASAR point-scanning (Supplementary Video 2). Furthermore, the flexibility of the web application interface enabled us to swap LysoTracker Yellow for other live-cell imaging dyes and recompute mixing matrices. These datasets also unmixed accurately, without the need for control measurements of singly-labelled cells to form the mixing matrix (Figure 3e,f,g, Supplementary Video 3, Supplementary Video 4 and Supplementary Video 5).

Attempts to extend our imaging from 2D+*t* to 3D+*t* were challenged by a noticeable level of phototoxicity after only a few timepoints (Supplementary Figure 16 and Supplementary Video 6), as expected for fast live-cell spinning disk confocal microscopy. While this could be partially mitigated by adding a recovery interval between timepoints, this significantly compromised imaging speed. In practice, we found that the maximum achievable volumetric imaging rate without significant photodamage was typically of the order of one volume per minute.

### Multispectral volumetric and projection light sheet microscopy

To reduce photodamage and increase volumetric acquisition speed, we next coupled our multispectral acquisition module to an oblique plane light sheet microscope (OPM) [19, 20]. Point spread function measurements showed that the module again did not compromise the resolution of the instrument, which was comparable to the resolution achieved with the spinning disk confocal microscope (Supplementary Figure 17). As before, the raw signals are correctly unmixed by RLSU into the expected six cellular compartments labelled by the ColorfulCell plasmid (see Figure 4a, Supplementary Video 7 and Supplementary Video 8). As expected, the gentler illumination strategy used by the multispectral OPM allowed us to routinely perform simultaneous six-colour volumetric imaging of cells for at least 200 timepoints, representing a more than ten-fold improvement compared to SDCM imaging (Supplementary Video 9).

**Figure 4:**
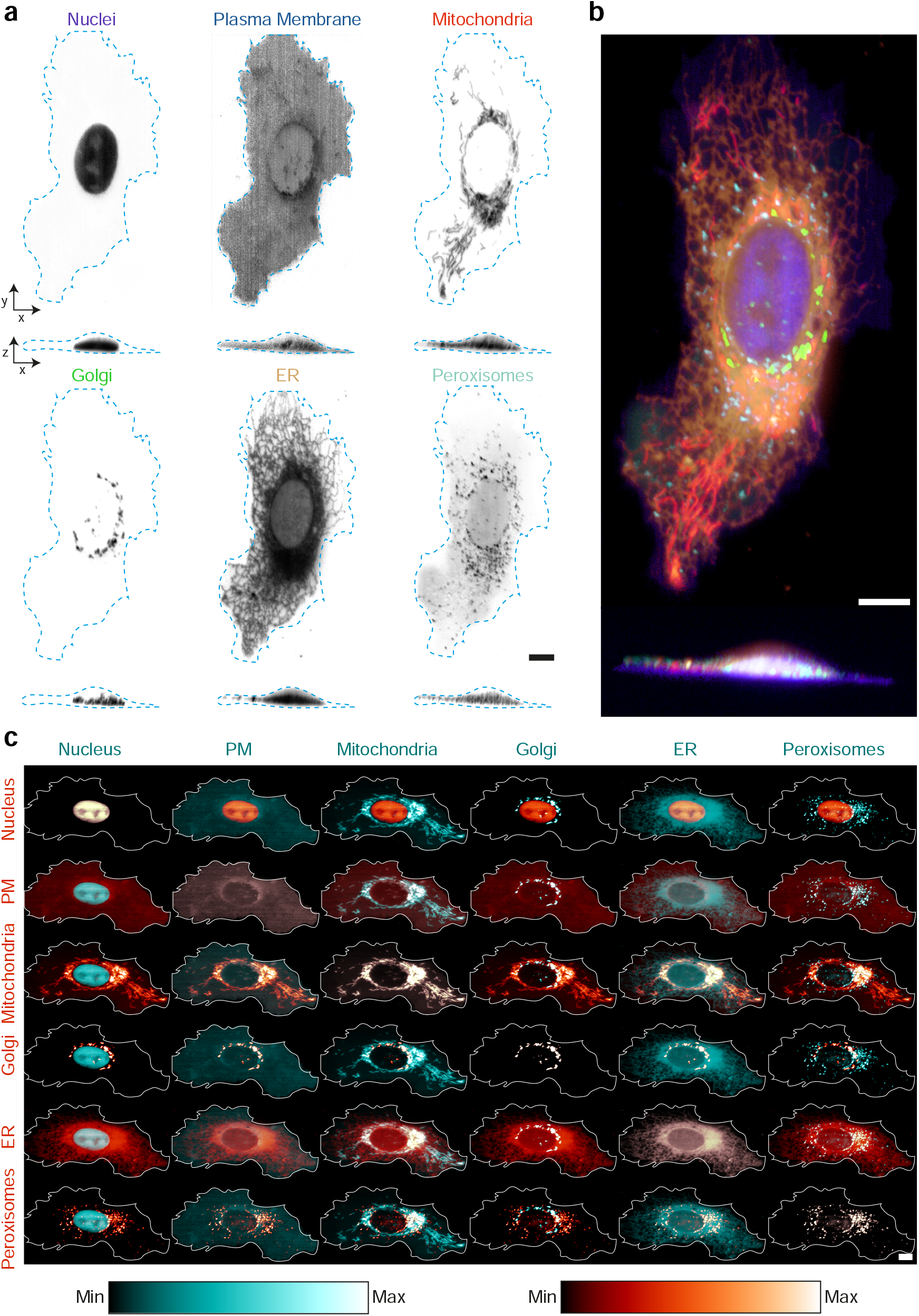
Multispectral live-cell volumetric imaging on an oblique plane light sheet microscope. **a** U2OS cells transfected with the ColorfulCell plasmid were imaged live by multispectral oblique plane light sheet microscopy followed by RLSU unmixing. Top-down and orthogonal view maximum intensity projections shown (cell boundaries in cyan). **b** Colour merge of individual channels shown in (a). **c** Pairwise-montage of channels shown in (a) (cell boundaries in white). Scale bars = 10 μm.

While the colour merge, shown in Figure 4b, provides a good overview of the cell, we found that it was often hard to discern the interactions between components with such high-dimensional datasets. To combat this, we used a ‘pair-wise montage’ visualisation (Figure 4c), which shows two components at a time. Each *n* of the *N* objects is assigned a first colour and repeated *N* times down the *n*th column, then assigned a second colour and repeated *N* times across the *n*th row. In this way, the leading diagonal of the visualisation shows the overlap of one object with itself (*i*.*e*. the object itself is displayed in the colour obtained by adding the first colour to the second), while other elements show the overlap of one object with the others. As the object combinations are symmetric about the diagonal, part of the visualisation could be excluded as it conveys the same information as is available elsewhere. However, in practice we found that it was helpful to see both orderings of the colours.

To achieve even faster multispectral imaging speeds, we utilised the recently described shear-warp angled projection technique [21]. By adding a galvo-based image shifting unit in front of the multispectral module and synchronising the sweep of the light sheet, focal plane and image shift, we could acquire projection images of an entire cell from arbitrary viewing angles in one camera exposure.

This combination achieves whole-cell multispectral projections at 10 Hz (Figure 5a,b and Supplementary Video 10). Laser powers were increased from those used for volumetric imaging in order to maintain signal levels, but we could not achieve sufficient irradiances for the 405 nm laser to provide acceptable signals from the TagBFP and Cerulean labels. As such, we elected to deactivate this laser to reduce any potential photodamage but continued to unmix data from cells transfected with the ColorfulCell plasmid with a six-component mixing matrix. After unmixing, both the TagBFP and Cerulean channels correctly contained no signal.

**Figure 5:**
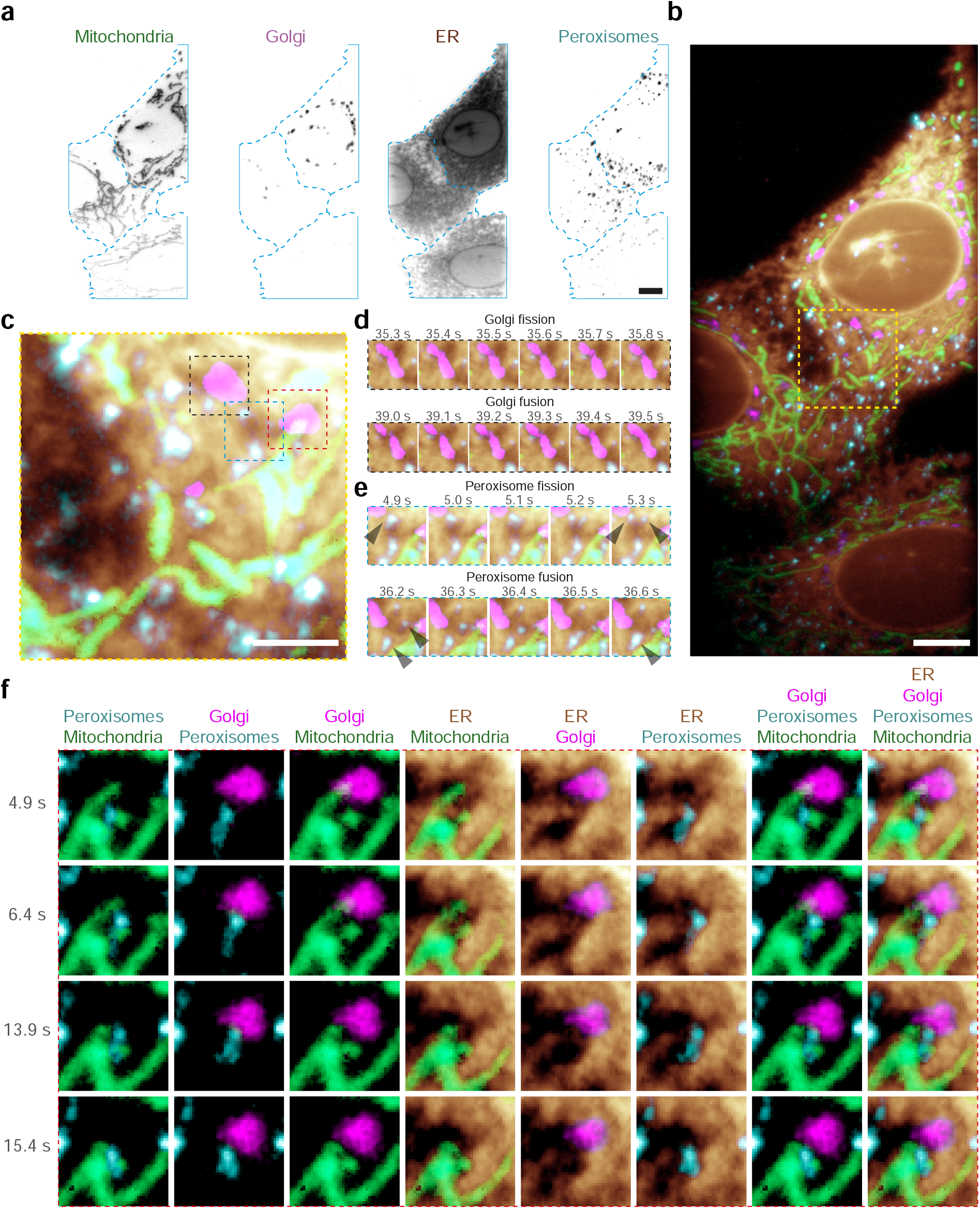
Multispectral live-cell projection imaging on a light sheet microscope. **a** U2OS cells transfected with the ColorfulCell plasmid were imaged by multispectral light sheet projection imaging followed by RLSU umixing (cell boundaries in cyan). Four channels of a single frame from a movie shown (Supplementary Video 10). **b** Colour merge of channels shown in (a). **c** Enlarged image of yellow box in (b). **d** Later frames showing Golgi fission, then fusion, in the black box in (c). **e** Frames from the cyan box in (c) showing peroxisome fission/fusion. Black arrows indicate peroxisomes that undergo fission or fusion within the frames shown. **f** Frames displaying two-way, three-way and four-way dynamic organelle interactions within the region delineated by the red box in (c). Scale bars = 10 μm.

The increased acquisition rate provided by the multispectral projection imaging allowed us to see both fast, sub-second organelle dynamics for hundreds of timepoints (*e*.*g*. ER remodelling), but also slower dynamics within the same movie. For instance, as can be seen in Figure 5d, part of the Golgi apparatus fissions in 0.5 s and then, 4 s later, refuses. Similarly, Figure 5e shows fission and fusion events for a number of peroxisomes on a similar timescale, and Figure 5f shows examples of two-, three- and four-way organelle interactions extracted from the same dataset.

It is worth emphasising that while in the above we have focussed on complex samples with a high number (six or seven) of fluorescent probes, multispectral imaging with our approach does not compromise the imaging quality or speed for simpler samples with just, say, two fluorophores. In fact, multispectral imaging is more photon efficient than conventional bandpass filter imaging systems, as light is merely redirected rather than rejected.

To illustrate this point, we performed two-color light-sheet imaging of macropinocytic cup closure in a *Dictyostelium* strain expressing LifeAct-mCherry and EGFP fused to a phosphatidylinositol (3,4,5)-trisphosphate (PIP3) reporter (Supplementary Figure 18). This is a particularly challenging sample, as *Dictyostelium* are known to be particularly light-sensitive and macropinocytic cup closure occurs rapidly in three dimensions, requiring fast volumetric imaging. After careful optimisation of exposure times and laser powers, we could reliably acquire movies containing hundreds of time-points without cell death, in line with our previous experience in imaging this strain on both our non-multispectral original OPM system and a field synthesis light sheet microscope [22]. In particular, we achieved full cell volumes at 2 Hz, which was enough to follow the process of macropinocytic cup closure and, by relying on simultaneous multispectral aquisition rather than sequential acquisition, the images were devoid of motion blur or colour misregistration (Supplementary Video 11).

### Imaging the dynamics of receptor sorting using de novo designed receptor minibinders

While fluorescent protein fusions provide fluorescence images free of non-specific binding, tagging at endogenous levels requires time-consuming genome editing, a process that must be repeated for each colour/target used. As a proof-of-concept experiment, we instead used de novo designed protein-binding proteins (minibinders), labelled with small molecule dyes, to visualise the endosomal sorting of endogenous cell surface receptors.

Endosomal sorting is the process by which cells sort different transmembrane receptors towards three major routes following their endocytosis: degradation in lysosomes, recycling back to the plasma membrane, or retrograde transport to the Golgi apparatus [23]. Imaging endosomal sorting is difficult, first because endosomes move rapidly in 3D and second because overexpression of fluorescently-labelled receptors can be detrimental. Recently, we developed pipelines to computationally design specific binders for proteins of interest [24, 25]. We therefore thought to fluorescently label minibinders, expressed and purified from *E. coli*, that recognise the extracellular region of endogenous transmembrane receptors, and use them to reveal the dynamics of receptor trafficking and sorting in cells (Supplementary Figure 19a). These de-novo-designed binders offer five advantages for multiplexed labelling of endogenous transmembrane receptors when compared to antibodies or fluorescent protein fusions introduced through genome editing:

1. they are small (∼80–100 amino acid residues) and so are less likely to perturb trafficking;
2. they are designed to express in, and purify from, bacteria well, facilitating access to the reagent;
3. they can be specifically labelled with bright organic dyes in a way that does not affect their target-binding ability, as opposed to non-specific labelling for antibodies;
4. they are designed to be monovalent and so do not induce receptor clustering;
5. they can be designed to be pH independent, so as not to detach from their target in the acidic environment of late endosomes.

As a proof-of-concept experiment, we fluorescently labelled four minibinders targetting the transferrin receptor (TfRB), the insulin-like growth factor 2 receptor (IGF2RB), the bone morphogenetic protein receptor type 2 (BMPR2B) and integrin *α*5*β*1 receptors (I*α*5*β*1B, see methods and Supplementary Figure 19b for minibinder design and Supplementary Figure 19c for biochemical characterisation). These four minibinders were labelled with a combination of fluorophores that we had previously validated could be unmixed when imaged using the multispectral OPM (see Supplementary Table 2 and Supplementary Figure 19). When incubated individually with cells, fluorescent minibinders internalise into punctate dynamic structures (Supplementary Figure 20a, see also Supplementary Figure 20b for control that minibinders are unlikely to be internalised by fluid-phase endocytosis). These structures were reminiscent of endosomes, which was confirmed by their colocalisation with WASH, an early/sorting endosome marker [26] (Supplementary Figure 20c,d and see also Supplementary Video 12). We thus incubated U2OS cells expressing nuclear TagBFP with all four binders simultaneously alongside labelled epidermal growth factor (EGF) and imaged them with the multispectral OPM (Figure 6b,c).

**Figure 6:**
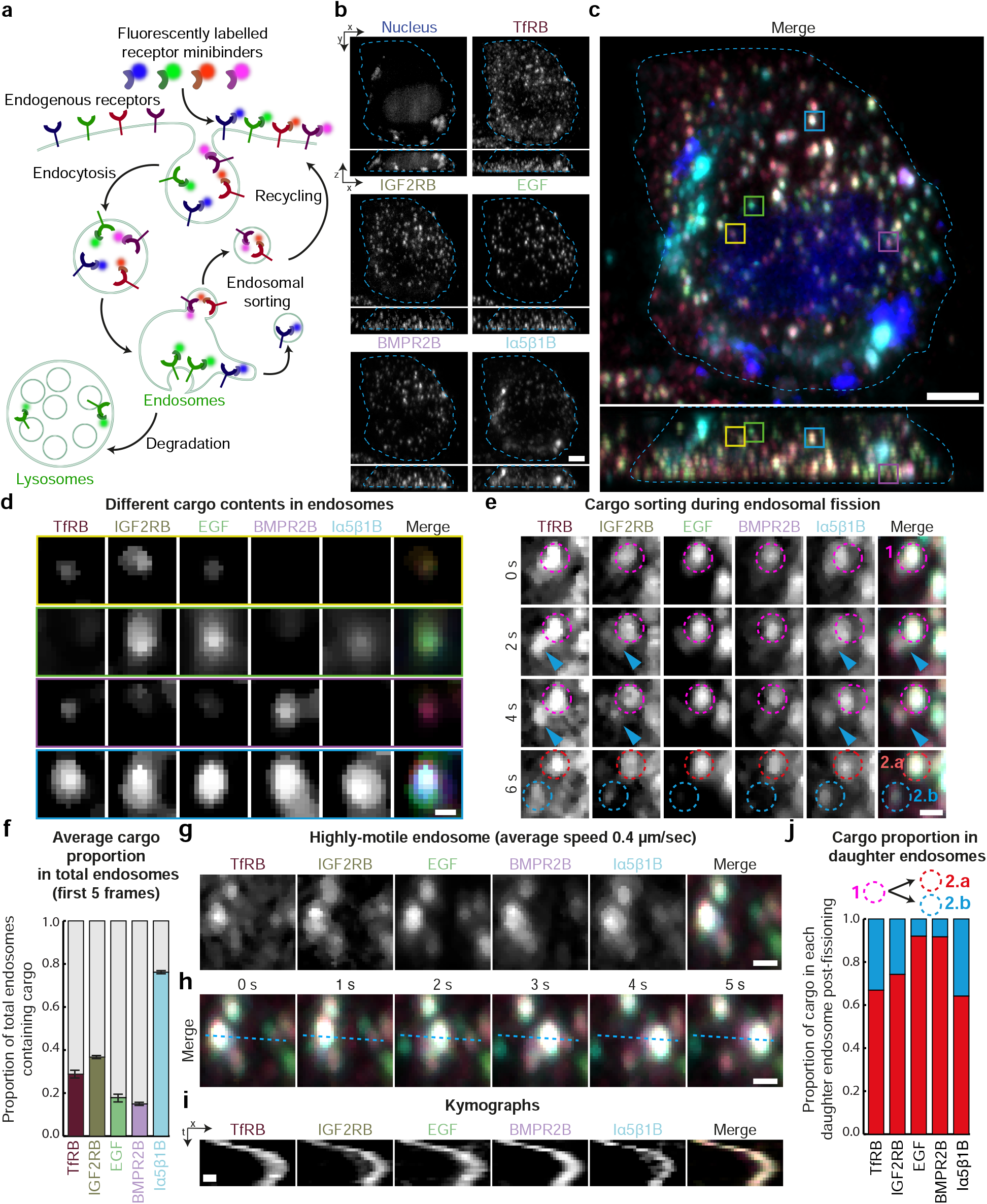
Multispectral light sheet imaging of intracellular trafficking using computationally designed receptor binders. **a** Schematic depicting the use of fluorescently-labelled, computationally designed receptor binders for imaging receptor trafficking. **b** Single frame of RLSU unmixed results (Supplementary Video 13) of a TagBFP-NLS-expressing HeLa Kyoto cell loaded with four fluorescently-labelled binders (TfRB, IGF2RB, BMPR2B & I*α*5*β*1B) and one fluorescently-labelled receptor ligand (EGF). Top-down and orthogonal view maximum intensity projections shown (cell boundaries in cyan). **c** Colour merge of channels in (b). **d** Individual channels and colour merge of endosomes indicated by yellow, green, magenta and cyan boxes in (c). **e** Individal channels and colour merge of a single endosome during cargo sorting. Frames from Supplementary Video 13 displaying fissioning of this compartment are shown. Magenta dashed circle indicates the parent endosome (1). Cyan arrow indicates the elongation of an intermediate tubule. Red and cyan circles indicate the two daughter endosomes post-fissioning (2.a and 2.b, respectively). **f** Graph depicting the average proportion of total endosomes containing each cargo *±* standard deviations across the first five frames of Supplementary Video 13. **g** Individual channels and colour merge of a highly-motile endosome at *t* = 0. **h** Colour merge of endosome depicted in (g) undergoing directional transport. **i** Kymographs of the highly-motile endosome shown in (g–h). **j** Graph depicting the cargo proportion transferred from the the parent endosome (magenta dashed circle. endosome 1) to each daughter endosome post fissioning (red and cyan dash circles, endosomes 2.a and 2.b respectively) in (e). Scale bars: b–c = 5 μm, d–e, g–h = 1 μm, i = 0.5 μm and 2.5 s.

Over time, binders were internalised and appeared in highly motile, diffraction-limited objects, as expected for receptors trafficking within the endocytic pathway (Supplementary Video 13). Importantly, the combination of the multispectral imaging module and the rapid imaging afforded by the OPM allowed us to achieve a volumetric imaging rate of 3 Hz in all channels (Supplementary Video 14). This not only allowed us to track all endosomes in the cell in 3D without motion blur but, also, with the absence of colour misregistration caused by delays in acquiring colour channels in typical sequential schemes (Figure 6g-i).

This allowed us to determine the content of each endosome, which revealed that individual compartments showed markedly different distributions of the different minibinders, and thus of their cognate receptors (Figure 6d quantified in f). For example, the cyan-boxed compartment in Figure 6c,d contains all labelled receptors and EGF, while the yellow-boxed compartment lacks integrin *α*5*β*1 and BMP receptor. Such a selective enrichment of specific receptors in specific endosomes is expected as we selected receptors known to traffic via different routes, and they would thus tend to be sorted away from each other (Supplementary Figure 19a).

The high spatiotemporal resolution of the multispectral OPM coupled to the increased brightness conferered by the fluorescent minibinders allowed us to directly image flow of the different receptors within the endomembrane system. For instance, Figure 6e shows an endosome containing all labelled binders, plus EGF. Within a few seconds this compartment elongates a tubule and fissions into two daughter endosomes, with one daughter endosome lacking EGF and BMPR2 (see also Figure 6j for quantification). Hence, during this fission event, the EGF and the BMP receptor binders have been sorted away from the integrin, IGF2 and transferrin receptor binders. Conversely, we could image fusion events during which the content of organelles exchange receptors (Supplementary Figure 21).

## Discussion

In summary, we have developed an iterative spectral unmixing algorithm capable of handling low SNR datasets along with camera-based multichannel acquisition hardware. Together, these developments enable fast multispectral microscopy on any camera-based microscope, such as the spinning disk confocal and oblique plane microscopes used here. The diffraction-limited hardware design ensures spatiotemporal resolution is uncompromised and, in the future, could be trivially extended to an increased number of channels. Currently, the post-acquisition spectral unmixing code takes longer to process a frame than the acquisition hardware takes to acquire it (∼1 s versus 10 ms to 250 ms). In the future, this speed could be improved using algorithmic acceleration, such as the approach described by Biggs and Andrews [27].

While the cameras we chose are inherently less sensitive than typical cameras used for scientific imaging (with quantum yields of ∼70 % vs. ∼95 %), the fact that light is merely redistributed to a different detector, rather than filtered, offsets this. If further sensitivity is required, an alternative arrangement using two scientific cameras and commerically available four-way image splitters could be used. We did not investigate this possibility here due to reasons of cost and the associated reduction in the usable field of view, but are currently developing such a system for single molecule imaging. While we saw no evidence that co-illumination with multiple simultaneous wavelengths led to an increased rate of photobleaching or phototoxicity, multispectral imaging in itself does nothing to reduce these effects. While one-colour imaging allows for the use of the most photostable fluorophore available, multispectral imaging necessarily requires the use of additional, less photostable, alternatives. As such, continued fluorophore improvement, such as the creation of StayGold [28], and the development of new technologies to decrease photodamage, such as optical triplet-state depletion [29] and engineered imaging media [30, 31], will be important in the future development of multispectral imaging.

The use of de novo designed minibinders, as demonstrated here, offers an attractive alternative to genetically encoded fluorophores as they can be used to label targets at endogenous levels with significantly more photostable small-molecule dyes. Such binders allowed us to study the sorting of endogenous cell-surface receptors without genome editing, an approach which could potentially be applied to patient-derived cells. However, while we foresee fluorescent binders as an upcoming reagent of choice for targets with an extracellular domain, further work is required to enable efficient cytosolic delivery for intracellular targets.

Altogether, the developments described herein enable the study of multiple, simultaneous sub-cellular interactions at sub-second timescales over minutes or hours. Crucially, this comes with no penalty in spatiotemporal resolution or detection sensitivity. In the future, we envisage extending the spectral range into the near infra-red to enable better deep tissue imaging, and combining the multispectral imaging unit with single molecule localisation microscopy to significantly enhance resolution.

## Acknowledgements

JDM thanks Andrew York and Rainer Heintzmann for interesting discussions regarding Richardson–Lucy deconvolution and acknowledges support from the Royal Society through a University Research Fellowship (URF\R1\221086 & RF\ERE\221078). KM acknowledges support from the Wellcome Trust through a Sir Henry Wellcome Postdoctoral Fellowship (220480/Z/20/Z). ED is funded by the MRC (MC_UP_1201/13) and Human Frontier Science Program (Career Development Award CDA00034/2017). JDM and ED thank Jake Grimmett, Toby Darling and Ivan Clayson for scientific computing infrastructure and support. AK and JDM thank the Light Microscopy Facility of the MRC LMB for access to the Zeiss LSM 710 instrument used in this work, and thank the Mechanical and Electronic Workshops of the MRC LMB for their assistance.

This work was supported by the Medical Research Council, as part of United Kingdom Research and Innovation (also known as UK Research and Innovation) [MC_UP_1201/13]. For the purpose of open access, the MRC Laboratory of Molecular Biology has applied a CC BY public copyright licence to any Author Accepted Manuscript version arising.

## Conflicts of interest

AK and JDM are inventors on a patent application describing the multispectral imaging approach (WO2024003253A1). XW, JG-M, DL, BH and DB are inventors on various patent applications describing minibinders used in this work. The other authors declare no competing interests.

## Methods

### Multispectral imaging unit design and construction

After deciding on an eight-channel system (see Supplementary Note E), we consulted catalogues of commercially available dichroic mirrors to select seven that would span the visible spectrum in roughly equal steps. We selected zt532rdc, zt594rdc, zt670rdc-xxrxt, zt491rdc, zt458rdc, zt514rdc and zt561rdc, all from Chroma Technology. Constructing a system that splits light onto eight separate cameras, rather than eight regions of one camera, both seemed simpler and would give a larger field of view. As such, we investigated machine vision systems that could support eight or more cameras and selected the xiX platform from Ximea GmbH. As we desired an effective pixel pitch similar to that of sCMOS cameras (6.5 μm/px) and wanted to use off-the-shelf tube lenses for the image relays (in order to maintain diffraction limited performance), we selected a combination of the IMX250 imaging chip (3.45 μm/px) with an *f* = 180 mm input lens and *f* = 100 mm output lenses (Ximea MX050MG-SY, Thorlabs TTL180-A, Thorlabs TTL100-A) to give an effective pixel pitch of 6.21 μm/px.

With these components selected, we designed a CAD model using Solidworks of an eight-way image splitter comprising a central aluminium alloy base plate to which all dichroics, cameras and output lenses were attached via holders and a ‘peninsula’ plate to hold the input lens and C-mount thread. With this design, the input lens focal length can be freely changed, with the peninsula swapped for one of the appropriate length to keep the input image plane in the correct location. The spacings between components had previously been checked using Zemax OpticStudio to ensure that diffraction-limited imaging was possible over the entire field-of-view (Supplementary Figure 11).

Components were milled in-house using a Haas TM 1E and assembled. To fix dichroics to the holders, we used DOWSIL 730 FS (Dow Corning) as this does not contract while curing (and hence does not warp the dichroic) and can easily be removed with a razor blade if required. Cameras were roughly placed near the focal plane of each output lens using measurements from the CAD model and then finely focussed by imaging the bricks of a distant building (∼700 m) through a window, having previously calculated that this distance would produce the same image location as imaging at infinity.

Cameras were controlled through Micro-Manager, using a manufacturer-supplied device adaptor. As the ‘Multi Camera’ device adaptor, which allows the use of multiple cameras, only supports four cameras by default, we edited and recompiled this component to increase the limit to eight. Camera synchronisation was handled via TTL signals and the manufacturer’s synchronisation box (xSWITCH, Ximea GmbH).

### Multispectral point-scanning confocal microscopy

For multispectral point-scanning confocal microscopy, U2OS cells expressing the ColorfulCell plasmid were imaged live on a commercial laser scanning confocal microscope (Zeiss LSM 710) equipped with a QUASAR detector, a 63×, 1.4 NA oil-immersion objective (Zeiss Plan-Apochromat, 63×/1.4 oil DIC 420782-9900) and a heated enclosure for imaging at 37 °C and 5 % CO2. The 405 nm (Diode 405-30), 488 nm (Argon laser, LASOS Lasertechnik GMBH, RMC 781 Z1), 561 nm (DPSS 561-10) and 633 nm (HeNe 633) excitation wavelengths were used for simultaneous excitation of the sample. For the UV light path, the MBS 405 dichroic mirror was selected to direct the 405 nm excitation light to the sample. For the visible light path, the MBS 488/561/633 dichroic mirror was selected to direct the excitation light from the 488 nm, 561 nm and 633 nm lasers to the sample. The resulting emission was imaged on the QUASAR detector (410.5 nm to 694.9 nm in 32 channels, 9.2 nm spectral resolution) by selecting the ‘Lambda’ mode and directing the light to ‘ChS Detector’ (Spectral detector) in the software. Live cells were imaged in 2D or 3D Z-stacks (1912 × 1912 pixels) with 4× line averaging, in line sequential mode, with a 2.196 μs pixel dwell time resulting in ∼18.8 s per plane imaged and a 70.6 nm effective pixel size. To calibrate the gain and hence SNR in Supplementary Figure 9, we used the single-image gain & offset calibration software developed by Heintzmann et al. [32].

### Multispectral spinning disk confocal microscopy

Imaging was performed on a spinning disk confocal instrument composed of a Zeiss AxioObserver microscope stand equipped with a fast-piezo stage (ASI), a 63×, 1.4 NA oil-immersion objective (Zeiss Plan-Apochromat, 63x/1.4oil DIC 420782-9900) and a spinning disk confocal unit (CrestOptics X-Light V3). The spinning disk confocal unit had two camera ports: a Photometrics Prime 95B back-illuminated sCMOS camera was attached to the perpendicular camera port whilst the eight-channel multispectral camera unit was attached to the straight-on camera port via a C-mount thread. A motorised mirror determined which detector emitted fluorescence was sent to. The multispectral camera unit was mounted on pedestal posts (Thorlabs) for stability and to ensure that the unit was at the correct height in relation to the optical path height of the spinning disk confocal unit. Excitation was provided via a multiline, solid-state laser illuminator (89-North LDI-7) with the following wavelengths available: 405, 445, 470, 520, 528, 555 and 637 nm. The entire system was operated by Micro-Manager. The eight cameras of the multispectral camera acquisition module were controlled by the xSWITCH (Ximea) where cameras 1-7 were set to general purpose input (GPI) whilst camera 8 was set to general purpose output (GPO). In the control software, the exposure-start trigger signals for cameras 1-7 were set to be synchronised to the rising edge exposure signal of camera 8 during an acquisition. In this way the eight cameras acquired data simultaneously. The other hardware components were controlled via a National Instruments DAQ card (NI PCIe-6321), where a 5 Volt general purpose output (GPO_5V) trigger from the xSWITCH box controlling the cameras was used to trigger the laser lines and piezo stage during acquisitions. In this way the lasers and the piezo stage motion were triggered based on the rising edge of the camera exposure signals. A quad-band dichroic mirror (ZET405/470/555/640rpc-UF1, Chroma), emission filter (ZET405/470/555/640m-OD8) and excitation cleanup filter (ZET405/470/555/640x, Chroma) were employed to enable simultaneous four colour excitation using the 405, 470, 555 and 637 nm lasers. The resulting emission from the sample was recorded on the eight cameras of the multispectral acquisition unit described above. The temperature of the sample was kept at 37 °C using a temperature control chamber (microscopeheaters.com) for live-cell imaging.

### Multispectral oblique plane light sheet microscopy

Oblique plane microscopy was performed on a custombuilt setup (see Supplementary Figure 22), which we previously described in a manuscript by Watson & Krüger et al. [33]. In brief, fluorescence was collected by a 100× 1.35 NA silicon-immersion objective (Nikon MRD73950) and directed through an *f* = 200 mm tube lens (Thorlabs TTL200-A) and *f* = 70 mm scan lens (Thorlabs CLS-SL) onto a galvanometric mirror (Thorlabs GVS001). This mirror acted to scan the image plane, and light sheet illumination, through the sample in a telecentric manner. Light reflected off the mirror was then directed through an *f* = 39 mm scan lens (Thorlabs LSM03-VIS), another *f* = 200 mm tube lens (Thorlabs TTL200-A) and a 40× 0.95 NA air-immersion objective (Nikon MRD70470) into the remote focussing volume [34]. A plane tilted at 30° to the optical axis was then imaged by a 40× 1.0 NA solid-immersion objective (AMS-AGY v1, Applied Scientific Instrumentation) and another *f* = 200 mm tube lens (Thorlabs TTL200-A) onto the input plane of the multispectral camera system.

Excitation light from 405 nm, 488 nm, 561 nm and 638 nm lasers (LBX-405-100-CSB-PPA, LBX-488-100-CSB-PPA and LBX-638-100-CSB-PPA, Oxxius & OBIS 561 nm LS 100 mW, Coherent) was made co-linear and coupled into a polarisation-maintaining single mode fibre (PM-S405-XP, Thorlabs). A reflective parabolic collimator (RC12APC-P01, Thorlabs) attached to the other end of the fibre produced a 12 mm beam (1/*e*^2^ diameter) which was focussed by an *f* = 50 mm achromatic cylindrical lens (LJ1695RM-A, Thorlabs) to form a light sheet. The NA of the light sheet could be varied via an adjustable slit (Thorlabs VA100CP) placed before the cylindrical lens. Another adjustable slit (Thorlabs VA100CP) was placed in the focal plane of the cylindrical lens to limit the lateral extent of the light sheet illumination. After cropping, the light sheet was directed by an *f* = 75 mm achromatic doublet (Thorlabs AC254-75-A-ML) and two kinematically-mounted mirrors onto a kinematically-mounted mirror located in a relayed pupil plane. The first two mirrors were used to position the light sheet illumination correctly in the pupil, with appropriate tilt, while the third mirror could be used to scan the light sheet from side to side in sample space, facilitating alignment. The illumination light was then coupled into the main beam path, between the second *f* = 200 mm tube lens and 40× 0.95 NA air-immersion objective with a quad-band dichroic beamsplitter (Semrock Di03-R405/488/561/635-t3-25×36).

As with the Crest V3 spinning disk confocal setup, the microscope was controlled via Micro-manager [35] and hardware synchronised with a National Instruments DAQ card (NI PCIe-6321), installed in a custom PC. Of note, this PC featured a 4× NVMe SSD PCIe mounting card (M.2 Xpander-Aero, MSI) to ensure data from all eight cameras could be written to disk without bandwidth limitations, and a 10G fibre network card to allow acquired data to be transferred to long-term storage promptly. Oblique plane light sheet microscopy stacks were computationally deskewed using our home-made software suite lsfm_tools, implementing a linear interpolation, which is available from https://github.com/jdmanton/lsfm_tools.

### Single channel spinning disk confocal microscopy

For sequential, single channel imaging of cells (Supplementary Figure 20b,c), imaging was performed using a spinning disk confocal instrument composed of a Nikon Ti stand equipped with a perfect focus system, a fast piezo Z-stage (ASI) and a 60×, 1.4 NA oil-immersion objective (Plan Apochromat Lambda), and a spinning disk head (Yokogawa CSU-X1). Images were recorded with a Photometrics Prime 95B back-illuminated CMOS camera run in pseudo global shutter mode and synchronised with the spinning disk wheel. Excitation was provided by 488 and 630 nm lasers (Coherent OBIS mounted in a Cairn laser launch) and imaged using dedicated single-bandpass filters for each channel mounted on a Cairn Optospin wheel (Chroma 525/50 for EGFP and Semrock 647lp for Alexa Fluor 647). The temperature was kept at 37 °C using a temperature control chamber (microscopeheaters.com). The system was operated with Metamorph.

### Image registration software

Image registration software was implemented in Python 3 using the `tifffile, numpy` and `imreg_dft` modules. Reference images, either of back-illuminated lens tissue or of spinning disk confocal pinholes, were collected and used to generate affine transformations to register each channel to channel 1 via cross-correlation (see examples in Supplementary Figure 12). Affine transform parameters were saved to disk and later used to register experimental data, before unmixing.

### Spectral unmixing software

Richardson–Lucy spectral unmixing software was developed in Python 3 using the `numpy, cupy, tifffile` and pandas modules. Briefly, registered TIFF stacks are loaded, along with a CSV file representing the mixing matrix for the experiment in question. For each z plane of the stack, image data are reshaped into a 2D matrix and the initial RLSU estimate is initialised with a matrix of all ones. Matrix multiplication is carried out on the GPU. After a user-defined number of iterations (default: 1000), RLSU is terminated and the RLSU result is reshaped into a hyperstack, which is then saved to disk as a TIFF. When processing oblique plane light sheet microscopy data, multispectral images were registered and unmixed first, then deskewed.

### Cell culture

U2OS FlipIn Trex cells and HeLa Kyoto cells (RRID:CVCL_1922) were cultured in DMEM-Glutamax (Gibco) supplemented with 10 % fetal bovine serum (FBS, Gibco) and 1 % penicillin-streptomycin (Gibco) at 37 °C with 5 % CO2. 3T3 FlipIn cells stably expressing EGFP WASH [26] were cultured in DMEM-Glutamax (Gibco) supplemented with 10 % Donor Bovine Serum (Gibco) and 1 % penicillin-streptomycin (Gibco) at 37 °C with 5 % CO2. Cells were transfected with Lipofectamine 3000 (Invitrogen) according to the manufacturer’s instructions, and imaged after one day of expression. Cells were regularly screened for mycoplasma using MycoAlert Mycoplasma Detection Kit (Lonza). On the day of the experiment, transfected (or stable) cells were always plated on glass-bottom dishes (World Precision Instruments, FD35) coated with fibronectin (Sigma, F1141, 50 μg/ml in PBS), for 2 h at 37 °C in DMEM-10 % serum. For imaging, the media was then changed to pre-warmed, filtered Leibovitz’s L15 medium (Gibco), enriched with 4.5 g/l glucose, 10 % FBS and 20 mM HEPES (Gibco). In some instances live cell dyes, such as SPY 555 tubulin, SPY 555 actin (Spirochrome) or LysoTracker Yellow HCK 123 (ThermoFisher Scientific) were used. Lyophilised live-cell dyes were reconstituted in 50 μl DMSO to prepare a 1000x stock solution as per manufacturer’s instruction. This stock solution was stored at *−*20 °C. On the day of the experiment the probe was diluted to a 1x staining solution in full growth media and vortexed briefly. Cells in imaging dishes were incubated at 37 °C with the staining solution for one hour. The staining solution was then removed, the cells were washed in PBS and warm imaging medium was added for imaging.

### *Dictyostelium* culture and imaging

*Dictyostelium discoideum* strain HM3478 used in this work is derived from Ax2(Kay) [36] by transformation [37] with the plasmid pPI304 (available from Addgene under the reference 113232) expressing reporters for PIP3 and actin (PH-PkbE-eGFP, LifeAct-mCherry, respectively). Cells were grown at at 22 °C in HL5 medium (Formedium, Hunstanton, UK) containing 10 μg/ml G418. They were harvested in log phase and imaged in SUM medium, which maintains macropinocytosis but reduces auto-fluorescence, as described [38, 39].

### Minibinder design

Full details of the design of the transferrin receptor minibinder are given by Sahtoe et al. [24], and details of the IGF2R minibinder design are given in Huang et al. [40]. These were designed using an established Rosetta design pipeline [25]. Manuscripts describing the design of the BMPR2 and integrin *α*5*β*1 minibinders are currently in preparation.

### Constructs

The ColorfulCell plasmid used throughout this manuscript, which encodes six fluorescent protein targetted to different organelles was described previously by Sladitschek et al. [17] and obtained from Addgene. For NLS-TagBFP expression (Figure 6), a construct encoding a nuclear localisation sequence (NLS) followed by TagBFP was synthethised by Twist Bioscience and cloned into a plasmid containing a CMV promoter.

For integrin *α*5*β*1, IGFR2 and BMPR2 minibinder production, genes encoding the minibinders were optimised for bacterial expression using codon optimisation and RNA ddG minimisation as described previously [41]. Minibinder sequences were then flanked in 5^*′*^ with a FseI restriction side followed by DNA sequence encoding a unique reactive cysteine KKCKK to enable downstream maleimide labelling (see below) and in 3^*′*^ by a 10×histidine tag for protein purification followed by a AscI restriction site. The resulting synthetic construct was synthesised by Twist Bioscience, and sub-cloned into a modified pGEX vector to express a protein of interest downstream of the gluthathione S transferase (GST) purification tag followed by TEV and 3C cleavage sequences [33]. Ultimately, this strategy resulted in minibinders flanked by GST-TEV-3C-KKCKK in N-terminus and 10×His in C-terminus (that is, GST-TEV-3C-KKCKK-Minibinder-10×His).

### Minibinder purification and labelling

Biochemical characterisation of the minibinders used in this study is presented in Supplementary Figure 19c. Unless stated otherwise, all purification steps were performed at 4 °C in the cold room. SDS-PAGE was performed using Nu-PAGE 4–12% Bis-Tris gels (ThermoFisher) according to the manufacturer’s instructions. InstantBlue (Sigma) was used for total protein staining of gels. Protein concentrations were determined using a BCA protein assay kit (Pierce) or by absorbance at 280 nm in a Nano Drop spectrophotometer (ThermoFisher) used in cuvette mode. In the latter case, molar absorption coefficients calculated from the amino acid sequences were used, and when characterising minibinders labelled with organic dyes, absorbance at 280 nm was corrected for dye absorbance using the correction factors provided by the manufacturer. In coomassie-stained gel imaging were performed on a ChemiDoc Touch instrument (Bio-Rad).

For integrin *α*5*β*1, IGF2R and BMPR2 minibinders, pGEX vectors encoding GST-TEV-3C-KKCKK-Minibinder-10×His (see above) were transformed into E. coli BL21(DE3) (Novagen, 69450-3), and bacteria were grown in 2×TY enriched with 100 μg/ml carbenicillin (Melford, C46000) at 37 °C, with 220 RPM shaking. Once culture reached an OD600 of 0.8, bacteria were then cooled down before induction at at 20 °C overnight (∼18 hours) with 0.5 mM isopropylthio-*β*-galactoside (IPTG, Generon, 02122). Cells were harvested through centrifugation at 4000 *g* for 20 minutes at 4 °C. The bacterial pellet was then stored at *−*20 °C until protein purification.

For GST-3C-integrin *α*5*β*1 minibinder-KKCKK-10×His purification, bacterial pellets were resuspended in 30 ml LYSIS buffer [25 mM HEPES, 150 mM NaCl, 1 mM DTT, 1 mM EDTA, 10 μg/ml DNAse, 1 mM MgCl_2_, 70 μg/ml lysozyme and cOmplete mini EDTA-free protease inhibitor cocktail (Roche, 11836170001)] per liter of cultured E. coli pellet. Resuspended bacteria were incubated at 4 °C for 15 mins with rolling prior to sonication on ice for 2 minutes 30 seconds (10 seconds on, 30 seconds off, 40 % amplitude). Lysate was then centrifuged at 40000 *g* for 40 minutes at 4 °C. The soluble supernatant was incubated with 1 ml glutathione sepharose 4B resin (GE Healthcare, 17-0756-01) for 2.5 hours at 4 °C with rolling. After incubation, resin and supernatant was poured into a gravity flow column and resin was washed extensively with LYSIS buffer without protease inhibitors/DNAseI/Lysosyme and eluted with ELUTION buffer [25 mM HEPES, 150 mM NaCl, 1 mM EDTA, 1 mM DTT, 10 mM reduced glutathione, pH 7.2]. To remove the reduced glutathione, eluted protein was dialysed overnight at 4 °C into STORAGE buffer [25 mM HEPES, 150 mM NaCl, 1 mM EDTA, 1 mM DTT, pH 7.2]. After dialysis, protein was incubated with 40 U of GST-tagged HRV 3C Protease (Pierce, 88946) to remove the GST tag and incubated overnight at 4 °C. Free GST and 3C protease were removed by incubating the dialysed protein with 1 ml of fresh glutathione sepharose 4B resin for 1.5 hours at 4 °C with rolling. Upon running through a disposable collumn (Bio-Rad), flow through containing the integrin *α*5*β*1 minibinder-KKCKK 10×His was collected and flash frozen in liquid nitrogen for storage.

IGF2R minibinder-KKCKK-10×His was purified in a similar fashion, with few modifications: LYSIS buffer was modified into [25 mM HEPES, 150 mM KCl, 1 mM DTT, 1 mM EDTA, 10 μg/ml DNAse, 1 mM MgCl_2_, 70 μg/ml lysozyme, 10 % glycerol + cOmplete mini EDTA-free protease inhibitor cocktail, pH 7.4]. GST-bound binder GST-minibinders bound to glutathione sepharose 4B were washed first in WASH1 [25 mM HEPES, 150 mM KCl, 1 mM DTT, 1 mM EDTA, 10 % glycerol, pH 7.4], secondly in WASH2 [WASH1 with 500 mM KCL final] and finally washed in WASH1 again. ELUTION buffer was [25 mM HEPES, 150 mM KCl, 1 mM DTT, 1 mM EDTA with 20 mM reduced glutathione, pH 8]. Eluted protein was dialysed overnight in [25 mM HEPES, 150 mM KCl, 1 mM DTT, 1 mM EDTA, pH 7.4] followed by further dialysis into STORAGE buffer [25 mM HEPES, 150 mM KCl, 1 mM EDTA, pH 7.4], before GST and 3C removal as above.

Similarly, for BMPR2 minibinder-KKCKK-10×His, LYSIS buffer was [25 mM HEPES pH 8, 150 mM NaCl, 1 mM DTT, 1 mM EDTA, 10 μg/ml DNAse, 1 mM MgCl_2_, 70 μg/ml lysozyme + cOmplete mini EDTA-free protease inhibitor cocktail]. GST-minibinders bound to glutathione sepharose 4B were washed first in WASH1 [25 mM HEPES pH 8, 150 mM NaCl, 1 mM EDTA, 1 mM DTT] and then washed in WASH2 [25 mM HEPES pH 8, 500 mM NaCl, 5 mM betamercaptoethanol, 20 % glycerol, 0.003 % DDM] and then washed a final time with WASH1. ELUTION buffer was [25 mM HEPES pH 8, 150 mM NaCl, 1 mM DTT, 1 mM EDTA and 10 mM reduced glutathione]. Eluted protein was incubated with 40 U of HRV 3C Protease to remove the GST tag and incubated overnight at 4 °C with simultaneous dialysis into [25 mM HEPES, 150 mM NaCl, 1 mM DTT, 1 mM EDTA, pH 7.4] followed by further dialysis the next day into STORAGE buffer [25 mM HEPES, 150 mM NaCl, 1 mM EDTA, pH 7.4]. GST and 3C protease were removed as above.

Single cysteine versions of a transferrin receptor minibinder (2DS25), as well as an interface knockout (KO) variant abolishing receptor binding were expressed and purified as previously reported [24].

### Minibinder fluorescent labelling

Biochemical characterisation, in particular in gel fluorescence of the minibinders used in this study is presented in Supplementary Figure 19c, and validation of their use in cells is presented in Supplementary Figure 20. For labelling of transferrin receptor minibinder, and respective interface knockout control, a 10-fold molar excess of Alexa Fluor 488 C5 maleimide (ThermoFisher) or maleimide Alexa Fluor 647 C2 maleimide (ThermoFisher) was added dropwise whilst vortexing to a 200 μL solution of minibinder at 1 mg/ml in [20 mM Tris pH 7.4, 100 mM NaCl, 1 mM TCEP]. After incubation overnight in the dark at 4 °C with rocking, free dye was removed through 6 sequential rounds of dilution into 500 μL PBS followed by concentration down to 50 μL using an Amicon Ultra 0.5 ml centrifugal filter unit with a 3 kDa MWCO - (Pierce).

For IGF2R minibinder labelling, a 10-fold molar excess of ATTO 514 maleimide (ATTO-TEC) was added dropwise to a 200 μL solution of minibinder in [SI25mM HEPES, SI150mM mM NaCl, SI1mM EDTA, pH 8] whilst vortexing, followed by a 3-hour incubation at room temperature with rocking in the dark. Excess free dye was removed through 9 sequential rounds of dilution into 500 μL PBS followed by concentration down to 50 μL using an Amicon Ultra 0.5 ml centrifugal filter unit with a 10 kDa MWCO.

For BMPR2 minibinder labelling, 10 μL of Alexa Fluor 647 C2 maleimide (ThermoFisher) was added dropwise to 1 mL of purified binder (10 μM) in STORAGE buffer [25 mM HEPES, 150 mM NaCl, 1 mM EDTA, pH 7.4], followed by a 2-hour incubation at room temperature with rocking in the dark. This ratio provides a 10-fold molar excess of dye over protein. Despite multiple attempts with different techniques, the excess free Alexa Fluor 647 dye could not be removed, but control experiments using the same amount of quenched dye alone showed that free dye at the low concentrations we use for binder experiments does not lead to appreciable endosomal signal (data not shown).

For integrin *α*5*β*1 minibinder labelling, purified binder was dialised against PBS to remove the DTT present in the STORAGE buffer. Then 25 μL of 0.2 M sodium bicarbonate, pH 9 was added to 250 μL protein in PBS to increase pH to 8.3 followed by dropwise addition of 10 μL of a 10 mM solution of CF680R NHS in DMSO while vortexing. After a 2-hour incubation at room temperature with rocking in the dark, free dye was removed through seven sequential rounds of dilution into 500 μL PBS followed by concentration down to 50 μL using an Amicon Ultra 0.5 ml centrifugal filter unit with a 3 kDa MWCO.

### Fluorescent minibinder uptake

To monitor receptor trafficking at endogenous levels (Figure 6), we used a pulse-chase experimental design with the purified fluorescent minibinders described above. HeLa Kyoto cells transiently expressing NLS-TagBFP for 24h were plated onto fibronectin-coated imaging dishes with a small 10 mm glass coverslip (World Precision Instruments, FD35-10) and allowed to adhere for one hour in 50 μl of DMEM media enriched with 10 % FBS and 1 % P/S. After the cells had adhered, the medium was removed and replaced with 50 μl of ice-cold DMEM containing 5 μl of TfR binder Alexa Fluor 488 (stock concentration 90 μM, so 6 μM final), 2 μl of IGF2R binder ATTO 514 (stock concentration 21 μM, so 0.6 μM final), 10 μl of BMPR2 binder Alexa Fluor 647 (stock concentration 10 μM, so 1.3 μM final), 5 μl of I*α*5*β*1 binder CF680R (stock concentration 200 μM, so 14 μM final) and 2 μg/ml of EGF Alexa Fluor 555 (catalogue number: E35350, ThermoFisher), all in PBS. Cells were then kept on ice for 30 minutes to prevent endocytosis and allow for fluorescently-labelled binder proteins to bind to their respective receptors. Note that the medium did not contain FBS in order to serum-starve the cells, which causes receptors to cluster at the plasma membrane. After 30 minutes on ice, the medium was removed, the cells were washed five times with PBS to remove excess unbound minibinder and the medium was replaced with 50 μl of pre-warmed L15 medium supplemented with 10 % FBS, 4.5 g/l of glucose and 20 mM of HEPES to trigger receptor endocytosis. After 10 minutes in the pre-warmed media the cells were imaged on the multispectral OPM. The 405, 488, 561 and 638 nm lasers were used to excite the six fluorophores in the sample simultaneously and the resulting emission was recorded for every plane on the eight cameras of the multispectral acquisition module. Eight-channel full cell volumes were imaged between 1-3 Hz, as per the imaging conditions described in Supplementary Table 3.

To show that each minibinder could be uptaken by cells individually (Supplementary Figure 20a), HeLa Kyoto cells were incubated with indicated minibinders at the same concentration as we used in the multiplexed uptake experiment described above, replacing the other binders by a corresponding volume of PBS, and cells were imaged on the multispectral OPM as above. To demonstrate that endocytosed minibinders indeed reached early-sorting endosomes, NIH-3T3 stably expressing GFP-WASH, a sorting endosome marker, were plated onto fibronectin-coated imaging dishes as above and incubated with 50 μl of L15 media supplemented with 10 % DCS, 20 mM of HEPES, 4.5 g/l glucose and with either 5 μl of TfR binder Alexa Fluor 647 (stock concentration 8.5 μM, so 0.7 μM final) or 10 μl of BMPR2 binder Alexa Fluor 647 (stock concentration 10 μM, so 1.7 μM final). The cells were incubated with the binders on the microscope at 37 °C for 10 minutes to allow receptor internalisation prior to imaging using the single channel spinning disk confocal microscope described above.

### Quantification of cargo abundances

To determine the proportion of the total endosomes containing each cargo in Supplementary Video 13, commercial image analysis software was used (Imaris, Oxford Instruments).

Briefly the five cargo channels were projected in ImageJ to create a pseudochannel corresponding to the total trafficking compartments present in the movie. In Imaris each channel (including the ‘total compartments’ pseudochannel) was segmented in 3D using the ‘Spots’ function, accounting for the anisotropy of the PSF. As the signals in each of the channels were near the diffraction-limit, the xy and z diameters used for segmentation were 0.5 μm and 1 μm, respectively. An object-based colocalisation method was carried out for each cargo channel with the ‘total comparmtents’ pseudo-channel in the software with the distance for colocalisation classified as being below 0.5 μm. This allows for the proportion of each cargo co-localising with the total segmented signals to be determined at each timepoint. For normalisation, the number of co-localising signals for each channel was determined as a ratio compared with the total segmented signals. This data was averaged across the first five timeframes of the movie, (Figure 6f).

## Notes

### Summary of Updates

Minor edits to the main text throughout, eight supplementary figures added, along with a supplementary note describing the appropriate Cramer-Rao lower bound.

