## Supplementary Material for "Multispectral live-cell imaging with uncompromised spatiotemporal resolution"

### A Richardson–Lucy spectral unmixing

In the Richardson–Lucy algorithm, an estimate of the underlying object, $u_t$, is updated by the iterative step:

$$u_{t+1} = u_t \cdot \mathrm{H}^{\mathrm{T}}\left(\frac{d}{\mathrm{H}u_t}\right), \tag{S1}$$

where $d$ is the measured (noisy) data, H is the measurement operator and $\mathrm{H}^{\mathrm{T}}$ is the transpose (dual) of the measurement operator [12, 13]. Both the multiplication and division operate elementwise. For deconvolution applications, H represents a convolution with the point spread function, with $\mathrm{H}^{\mathrm{T}}$ being a convolution with the flipped (in $x$, $y$ and $z$) point spread function.

For Richardson–Lucy spectral unmixing (RLSU), we set H to the mixing matrix that describes the contribution of each underlying object to each channel. For example, consider one object, A, that contributes 5 counts to channel 1 for every 15 counts to channel 2, along with another object, B, that contributes 0 counts to channel 1 (*i.e.* all counts go into channel 2). Then,

$$\mathrm{H} = \begin{bmatrix} 0.25 & 0.00 \\ 0.75 & 1.00 \end{bmatrix}, \tag{S2}$$

and

$$\mathrm{H}^{\mathrm{T}} = \begin{bmatrix} 0.25 & 0.75 \\ 0.00 & 1.00 \end{bmatrix}. \tag{S3}$$

Let us now say that, in an experiment, A emits with a rate of 100 photons/s and B emits with a rate of 200 photons/s. For perfect detection efficiency over an integration time of 1 s, we would then expect:

$$\hat{d} = \begin{bmatrix} 0.25 & 0.00 \\ 0.75 & 1.00 \end{bmatrix} \begin{bmatrix} 100 \\ 200 \end{bmatrix} = \begin{bmatrix} 25 \\ 275 \end{bmatrix}. \tag{S4}$$

However, as we are dealing with counts, Poisson statistics apply and so the elements of $d$ are drawn from Poisson distributions with the elements of $\hat{d}$ as rate parameters. As such, we may in fact measure:

$$d = \begin{bmatrix} 33 \\ 284 \end{bmatrix}. \tag{S5}$$

Traditional spectral unmixing (linear unmixing, LU) would then apply the (pseudo)inverse of H to $d$, to obtain:

$$\begin{bmatrix} 4 & 0 \\ -3 & 1 \end{bmatrix} \begin{bmatrix} 33 \\ 284 \end{bmatrix} = \begin{bmatrix} 132 \\ 185 \end{bmatrix}. \tag{S6}$$

Hence, after standard linear unmixing, we infer that A emits with a rate of 132 photons/s and B emits with a rate of 185 photons/s.

For RLSU, we begin with an initial estimate:

$$u_0 = \begin{bmatrix} 1 \\ 1 \end{bmatrix}, \tag{S7}$$

which produces predicted data

$$\mathrm{H}u_0 = \begin{bmatrix} 0.25 & 0.00 \\ 0.75 & 1.00 \end{bmatrix} \begin{bmatrix} 1 \\ 1 \end{bmatrix} = \begin{bmatrix} 0.25 \\ 1.75 \end{bmatrix}. \tag{S8}$$

This choice of $u_0$ is arbitrary, but all elements must be positive definite in order to ensure a positive result. Note that the predicted data are not necessarily integer, even though the real measured data are. The first updated estimate is then given by:

$$u_1 = \begin{bmatrix} 1 \\ 1 \end{bmatrix} \cdot \begin{bmatrix} 0.25 & 0.75 \\ 0.00 & 1.00 \end{bmatrix} \begin{bmatrix} 33/0.25 \\ 284/1.75 \end{bmatrix} = \begin{bmatrix} 154.7143\ldots \\ 162.2857\ldots \end{bmatrix} \tag{S9}$$

Further updates then give:

$$u_2 = \begin{bmatrix} 151.4032 \\ 165.5968 \end{bmatrix}, \quad u_3 = \begin{bmatrix} 148.5256 \\ 168.4744 \end{bmatrix}, \quad u_4 = \begin{bmatrix} 146.0386 \\ 170.9614 \end{bmatrix}, \quad u_5 = \begin{bmatrix} 143.8994 \\ 173.1006 \end{bmatrix}, \quad \ldots \tag{S10}$$

After the 24th iteration, we have

$$u_{24} = \begin{bmatrix} 132.4331 \\ 184.5669 \end{bmatrix}, \tag{S11}$$

which rounds to

$$\hat{u} = \begin{bmatrix} 132 \\ 185 \end{bmatrix}. \tag{S12}$$

Further iterations produce floating point numbers ever closer to 132 and 185, *i.e.* RLSU converges to the same solution as obtained by applying the inverse of the mixing matrix to the measured data (Supplementary Equation S6).

At first glance, this process may seem pointless — we have used many matrix multiplications to obtain the same result as we could easily obtain with just one. However, now consider a modified, more realistic, experiment, for which we have:

$$H = \begin{bmatrix} 0.0882 & 0.0009 & 0.0166 & 0.0081 \\ 0.5002 & 0.0365 & 0.0206 & 0.2662 \\ 0.2697 & 0.2896 & 0.0171 & 0.5238 \\ 0.1419 & 0.6731 & 0.9457 & 0.2019 \end{bmatrix}, \quad \hat{d} = \begin{bmatrix} 11.3798\ldots \\ 82.3518\ldots \\ 110.0174\ldots \\ 196.2509\ldots \end{bmatrix}, \quad d = \begin{bmatrix} 13 \\ 96 \\ 104 \\ 186 \end{bmatrix}, \tag{S13}$$

with each underlying object producing an expected count of 100. Now, RLSU produces an estimate of

$$u_{\text{RLSU}} = \begin{bmatrix} 111.3737 \\ 0.0000 \\ 150.6401 \\ 136.9862 \end{bmatrix}, \tag{S14}$$

while the standard LU approach produces

$$u_{\text{inv}} = \begin{bmatrix} 98.5806 \\ -46.9786 \\ 179.4886 \\ 167.9094 \end{bmatrix}. \tag{S15}$$

As well as producing an unphysical negative number, the LU estimates for the third and fourth component are pushed further away from the ground truth value of 100. Calculating root-mean-square-errors gives 90.2 for LU and 59.3 for RLSU. Setting the second component of $u_{\text{inv}}$ to zero and recomputing the RMSE gives 72.3, *i.e.* still higher than that of RLSU. Arguably, the Kullback–Leibler divergence is a more appropriate measure of accuracy, but this cannot be computed for the LU approach due to the negative number present in the estimate.

Together, these (somewhat contrived) demonstrations illustrate the general observation that RLSU produces estimates equivalent to LU when LU predicts all-positive estimates, but produces more accurate estimates when LU predicts estimates containing some negative numbers. Importantly, the RLSU estimates in this case are more accurate than a modified LU estimate where all negative numbers have been zeroed.

#### B Simultaneous spectral unmixing and deconvolution

As both diffractive blurring and spectral mixing are linear operations ($H_{blur}$ and $H_{mix}$, respectively), their combined effect is also a linear operator ($H_{full}$) and hence should be amenable to inversion using the Richardson–Lucy algorithm. To test this, we simulated datasets in which eight ground truth objects corresponding to the letters in the word SPECTRUM (Supplementary Figure 2a) were each blurred with eight different point spread functions (PSFs, with size increasing from bluest channel to reddest channel). These blurred images were then mixed according to a mixing matrix (thereby completing the action of the composite operation) and Poisson (shot) noise added (Supplementary Figure 2b).

To calculate a Richardson–Lucy iteration, we need to know the form of the transpose of the operation, $H_{full}^T$. Following linear algebra, we know that

$$H_{full}^T = (H_{mix}H_{blur})^T = H_{blur}^T H_{mix}^T, \tag{S16}$$

and so we need to mix with the transpose of the mixing matrix first, and then blur with flipped PSFs. Compared with normal Richardson–Lucy spectral unmixing (RLSU, Supplementary Figure 2c), incorporating deconvolution into the unmixing produces sharper images without compromising the quality of the unmixing (Supplementary Figure 2). We validate that the combined action of unmixing and deconvolving does not lead to errors in signal quantification by comparing the mean value of each ground truth object to those produced by standard linear unmixing, RLSU and the combined deconvolution-unmixing (Supplementary Figure 2). Furthermore, Supplementary Figure 2 shows that the mean squared error between ground truth and unmixed objects is minimised by the deconvolution-unmixing approach. However, a visual inspection shows that the deconvolution-unmixing results suffer from the same kind of high-frequency ‘squiggle’ artefacts that plague normal Richardson–Lucy deconvolution and so we expect that, in the future, better results will be obtained by incorporating some form of explicit or implicit regularisation. It must be emphasised that these artefacts arise from the deconvolution part of the iteration and are not present in the Richardson–Lucy spectral unmixing as used in the rest of the manuscript.

#### C Cramér–Rao lower bound analysis of spectral unmixing

To calculate the Cramér–Rao lower bound for estimating objects given Poisson-noisy mixed signals, we begin with the model for the expected photon counts:

$$\mu_k = \sum_{l=1}^{k} H_{kl} x_l, \quad k = 1, 2, \ldots, K \tag{S17}$$

for $L$ fluorophore species and $K$ channels. For the sake of simplicity, we assume that $K = L$ and that the mixing matrix, $H$, is invertible. Furthermore:

$$H_{kl} = \int d\lambda F_k(\lambda) G_l(\lambda), \tag{S18}$$

with $F_k(\lambda)$ the detection spectrum and $G_l(\lambda)$ the emission spectrum, is assumed to be normalised such that

$$\sum_k H_{kl} = 1 \quad \text{for all } l. \tag{S19}$$

Then, the total number of expected detected photons satisfies:

$$\sum_k \mu_k = \sum_{k,l} H_{kl} x_l = \sum_l x_l, \tag{S20}$$

so that in the unmixing the total number of photons is conserved.

The log-likelihood is:

$$\log \mathscr{L} = \sum_k \left[ m_k \log \mu_k - \mu_k - \log(m_k!) \right], \tag{S21}$$

with $m_k$ the observed photon counts. These satisfy $\langle m_k \rangle = \mu_k$ and $\langle m_k m_l \rangle = \mu_k \mu_l + \mu_k \delta_{kl}$, according to Poisson statistics. The derivative of $\log \mathscr{L}$ with respect to the estimated $x_l$ is:

$$\frac{\partial \log \mathscr{L}}{\partial x_l} = \sum_k \frac{\partial \log \mathscr{L}}{\partial \mu_k} \frac{\partial \mu_k}{\partial x_l} = \sum_k \left[ \frac{m_k}{\mu_k} - 1 \right] H_{kl} = \sum_k \frac{m_k}{\mu_k} H_{kl} - 1. \tag{S22}$$

For the Cramér–Rao lower bound we need the Fisher information matrix:

$$F_{ij} = \langle \frac{\partial \log \mathscr{L}}{\partial x_i} \frac{\partial \log \mathscr{L}}{\partial x_j} \rangle = 1 - \sum_k \frac{\langle m_k \rangle}{\mu_k} H_{ki} - \sum_k \frac{\langle m_k \rangle}{\mu_k} H_{kj} + \sum_{k,l} \frac{\langle m_k m_l \rangle}{\mu_k \mu_l} H_{ki} H_{lj} = \sum_k \frac{1}{\mu_k} H_{ki} H_{kj}. \tag{S23}$$

We can write this conveniently in matrix form as:

$$F = H^T M^{-1} H, \tag{S24}$$

with $M_{kl} = \mu_k \delta_{kl}$ and $\mathrm{M}^{-1}_{kl} = \frac{1}{\mu_k}\delta_{kl}$. Now, the inverse of F follows as:

$$\mathrm{F}^{-1} = \mathrm{H}^{-1}\mathrm{M}(\mathrm{H}^{-1})^{\mathrm{T}}, \tag{S25}$$

and so the Cramér–Rao lower bound is:

$$\langle \Delta x_l^2 \rangle \geq \left[\mathrm{F}^{-1}\right]_{ll} = \sum_k \mu_k \left[(\mathrm{H}^{-1})_{lk}\right]^2. \tag{S26}$$

This is also what one would expect if the estimator is:

$$\hat{x}_l = \sum_k (\mathrm{H}^{-1})_{lk} m_k, \tag{S27}$$

and then, using error propagation:

$$\langle \Delta \hat{x}_l^2 \rangle = \sum_{k,p} (\mathrm{H}^{-1})_{lk}(\mathrm{H}^{-1})_{lp} \langle \Delta m_k \Delta m_l \rangle = \sum_k \mu_k \left[(\mathrm{H}^{-1})_{lk}\right]^2. \tag{S28}$$

To demonstrate that we approach this Cramér–Rao lower bound, we ran a simulation of eight ground truth objects, each with value 100, mixed using the following mixing matrix (obtained from our web app using spectra for TagBFP, Citrine, mAzamiGreen, Qdot 565, Qdot 585, Qdot 605, Qdot 655 and Qdot 705):

$$\begin{bmatrix} 0.0093 & 0.2376 & 0.0005 & 0.0003 & 0.0000 & 0.0000 & 0.0001 & 0.8259 \\ 0.1681 & 0.4641 & 0.0190 & 0.0030 & 0.0007 & 0.0003 & 0.0010 & 0.0973 \\ 0.3825 & 0.1817 & 0.0682 & 0.0038 & 0.0006 & 0.0003 & 0.0010 & 0.0431 \\ 0.2671 & 0.0908 & 0.6250 & 0.1201 & 0.0322 & 0.0193 & 0.0182 & 0.0213 \\ 0.1180 & 0.0253 & 0.2742 & 0.6957 & 0.4525 & 0.0077 & 0.0016 & 0.0096 \\ 0.0390 & 0.0001 & 0.0090 & 0.1597 & 0.4916 & 0.2084 & 0.0264 & 0.0024 \\ 0.0147 & 0.0003 & 0.0040 & 0.0169 & 0.0217 & 0.7313 & 0.3742 & 0.0003 \\ 0.0013 & 0.0000 & 0.0001 & 0.0004 & 0.0007 & 0.0326 & 0.5774 & 0.0000 \end{bmatrix} \tag{S29}$$

1,000 independent realisations of Poisson (shot) noise were generated and the noisy signals unmixed using RLSU. Supplementary Figure 5 shows the distribution of the reconstructed values, with the means shown as black triangles. These are correctly located around 100, with the experimental standard deviation (shown as a solid errorbar) matching well with the square root of the Cramér–Rao lower bound (shown as a dotted errorbar).

#### D Effects of decreased spectral separation on unmixing results

As RLSU was developed to reconstruct ground truth objects from spectrally overlapping datasets, we wanted to assess its performance when decreasing the degree of spectral separation between fluorophores. Two channel spectrally-overlapping Poisson (shot) noisy datasets were simulated using ground truth objects ‘S’ and ‘P’, the emission spectra of EGFP and EYFP, and an idealised dichroic mirror with an infinitely steep cut-on wavelength equidistant from the spectral peaks of EGFP (510 nm) and EYFP (530 nm). EGFP and EYFP were chosen as they have significant spectral overlap, near identical spectral shape and only 20 nm peak-to-peak spectral separation. As shown in Supplementary Figure 8a, after 100 iterations, RLSU was able to faithfully reconstruct the ground truth objects ($\mathrm{RMSE}_S = 0.289, \mathrm{RMSE}_P = 0.247$) whilst the results from linear unmixing contained many negative pixel values (indicated in red) and erroneous pixel reassignment ($\mathrm{RMSE}_\mathrm{S} = 0.604, \mathrm{RMSE}_\mathrm{P} = 0.636$).

To further test the degree of spectral separation required for accurate reconstruction via RLSU, the EGFP and EYFP emission spectra were shifted by 5 nm towards the cut-on wavelength of the dichroic mirror resulting in a 10 nm peak-to-peak spectral separation (Supplementary Figure 8b). In this condition, RLSU could still accurately reconstruct the ground truth objects ($\mathrm{RMSE}_\mathrm{S} = 0.823, \mathrm{RMSE}_\mathrm{P} = 0.851$, compared with $\mathrm{RMSE}_\mathrm{S} = 0.998, \mathrm{RMSE}_\mathrm{P} = 1.039$ for linear unmixing). This was also true with a 4 nm peak-to-peak spectral separation (Supplementary Figure 8c, $\mathrm{RMSE}_\mathrm{S} = 1.829, \mathrm{RMSE}_\mathrm{P} = 1.897$ for RLSU and $\mathrm{RMSE}_\mathrm{S} = 2.11, \mathrm{RMSE}_\mathrm{P} = 2.182$ for linear unmixing). However, with a 0 nm spectral separation the marginally different spectral shapes of EGFP and EYFP provided insufficient information for accurate spectral unmixing (Supplementary Figure 8d).

#### E Selecting the optimum number of channels

A dichroic mirror splits multicolour light into two paths, one transmitted and one reflected. Hence, by arranging dichroic mirrors in a full-branched tree-like format, $2^N$ colour channels are produced using $2^N - 1$ dichroic mirrors, where $N$ is the number of levels of the tree. One, two and four channels are each an insufficient number, while 32 channels or more would require an infeasible number of dichroic mirrors. The question is then whether the extra spectral information provided by 16 channels outweighs the significantly greater investment in optics and detectors required compared to an eight-channel system.

As most samples are only labelled with fluorophores emitting in the 450 nm to 650 nm range, we can consider imaging the same 200 nm spectral range with either 16 channels in 12.5 nm wavebands, or eight 25 nm wavebands. A 25 nm bandwidth is similar to that employed in many single-channel imaging systems using a bandpass filter, meaning that raw signal levels would be comparable. Furthermore, Jahr et al. previously showed using their hyperspectral light sheet microscope that a 25 nm bandwidth still provided sufficient information to spectrally unmix EYFP from EGFP [6], two of the most spectrally similar fluorophores in common usage.

This suggests that the only advantage of a 16-channel system would be if more than eight fluorophore species needed to be unmixed (as $N$ channels are required to unmix $N$ species), or if the flexibility to image across a larger spectral range (*i.e.* keeping the same 25 nm waveband width but extending into the far-red and near-infrared) were needed. As such, we selected eight channels, and hence seven dichroic mirrors, as the optimum number. However, we note that, if more than eight, but less than 16, channels are desired, this can be achieved without investment in a full 16-channel system by adopting a non-fully-branched tree layout (*e.g.* nine channels can be produced by just adding one dichroic mirror and an additional detector to an appropriate path). The number of unmixable components can be extended beyond eight, while maintaining eight detection channels, by also incorporating excitation-based unmixing, but this precludes the possibility of simultaneous capture of all necessary data and hence compromises temporal resolution.

### F USAF 1951 target resolution testing

Testing the resolution of the assembled system using a USAF 1951 target showed that at least group 5–6 could be clearly resolved across all channels, corresponding to 57.0 lp/mm (Figure 2e). For camera 1, viewing ~ 465 nm, group 6–3 was resolvable, while 6–4 was not. 6–3 corresponds to 80.6 lp/mm, close to the Nyquist-sampling limit of 80.5 lp/mm.

| Receptor | Abbreviation | Size / aa | Affinity / nM | Fluorophore |
|---|---|---|---|---|
| Transferrin receptor | TfRB | 88 (128) | 30 | Alexa Fluor 488 |
| Insulin-like growth factor 2 receptor | IGF2RB | 124 (164) | 40 | ATTO 514 |
| Bone morphogenetic protein receptor type 2 | BMPR2B | 59 (99) | 4 | Alexa Fluor 647 |
| Integrin $\alpha 5\beta 1$ | I$\alpha 5\beta 1$B | 65 (105) | 2.2 | CF680R |

**Supplementary Table 2: De novo designed protein-binding protein details.**
For size column, number in brackets includes PC-tag, N- and C-terminal linkers and 10×His tag.

| Figure | Modality | Cell type | Pixel pitch / nm | $Z$ spacing / nm | Exposure / ms | Interval / s | $N_Z$ | $N_C$ | $N_T$ | $\lambda_{ex}$ /nm | Labels | LUTs |
|---|---|---|---|---|---|---|---|---|---|---|---|---|
| 3a,c,d | SDCM | U2OS (37 °C) | 98.5 | N/A | 100 | 15 | 1 | 7 | 50 | 405 | NLS-TagBFP | Blue |
| | | | | | | | | | | | Lyn-Cerulean | Duo intense cyan |
| | | | | | | | | | | 470 | MTS-mAzamiGreen | Red |
| | | | | | | | | | | | GTS-Citrine | Green |
| | | | | | | | | | | 555 | LysoTracker Yellow HCK-123 | Magenta |
| | | | | | | | | | | | mCherry-KDEL | Circus yellow |
| | | | | | | | | | | 640 | iRFP670-SKL | Cyan |
| 3e | SDCM | U2OS (37 °C) | 98.5 | N/A | 100 | 15 | 1 | 6 | 100 | 405 | NLS-TagBFP | Blue |
| | | | | | | | | | | | Lyn-Cerulean | Duo intense cyan |
| | | | | | | | | | | 470 | MTS-mAzamiGreen | Red |
| | | | | | | | | | | | GTS-Citrine | Green |
| | | | | | | | | | | 555 | mCherry-KDEL | Circus yellow |
| | | | | | | | | | | 640 | iRFP670-SKL | Magenta |
| 3f | SDCM | U2OS (37 °C) | 98.5 | N/A | 200 | 15 | 1 | 7 | 25 | 405 | NLS-TagBFP | Blue |
| | | | | | | | | | | | Lyn-Cerulean | Duo intense cyan |
| | | | | | | | | | | 470 | MTS-mAzamiGreen | Red |
| | | | | | | | | | | | GTS-Citrine | Green |
| | | | | | | | | | | 555 | SPY555-tubulin | Pop cyan |
| | | | | | | | | | | | mCherry-KDEL | Circus yellow |
| | | | | | | | | | | 640 | iRFP670-SKL | Magenta |
| 3g | SDCM | U2OS (37 °C) | 98.5 | N/A | 300 | 45 | 1 | 7 | 25 | 405 | NLS-TagBFP | Blue |
| | | | | | | | | | | | Lyn-Cerulean | Grey |
| | | | | | | | | | | 470 | MTS-mAzamiGreen | Red |
| | | | | | | | | | | | GTS-Citrine | Green |
| | | | | | | | | | | 555 | SPY555-actin | Magenta |
| | | | | | | | | | | | mCherry-KDEL | Circus yellow |
| | | | | | | | | | | 640 | iRFP670-SKL | Cyan |
| 4a | OPM | U2OS (37 °C) | 111.6 | 500 | 250 | 10 | 201 | 6 | 1 | 405 | NLS-TagBFP | Blue |
| | | | | | | | | | | | Lyn-Cerulean | Duo intense cyan |
| | | | | | | | | | | 488 | MTS-mAzamiGreen | Red |
| | | | | | | | | | | | GTS-Citrine | Green |
| | | | | | | | | | | 561 | mCherry-KDEL | Circus yellow |
| | | | | | | | | | | 638 | iRFP670-SKL | Grey |

**Supplementary Table 3: Experimental details.**
$N_Z$ is the number of z-planes acquired, $N_C$ is the number of different fluorophores used, $N_T$ is the number of timepoints and $\lambda$ is the wavelength of laser excitation used. LUTs refers to the ImageJ lookup-tables used to visualise data.

| Figure | Modality | Cell type | Pixel pitch / nm | $Z$ spacing / nm | Exposure / ms | Interval / s | $N_Z$ | $N_C$ | $N_T$ | $\lambda_{ex}$ /nm | Labels | LUTs |
|---|---|---|---|---|---|---|---|---|---|---|---|---|
| 5 | OPM | U2OS (37 °C) | 111.6 | Projection | 100 | 0.1 | N/A | 4 | 500 | 488 | MTS-mAzamiGreen | Green |
| | | | | | | | | | | | GTS-Citrine | Magenta |
| | | | | | | | | | | 561 | mCherry-KDEL | Circus yellow |
| | | | | | | | | | | 638 | iRFP670-SKL | Cyan |
| 6 | OPM | HeLa (37 °C) | 111.6 | 750 | 19 | 1.3 | 67 | 6 | 35 | 405 | NLS-TagBFP | Blue |
| | | | | | | | | | | 488 | Alexa Fluor 488 TfR binder | Quartetto red |
| | | | | | | | | | | | ATTO 514 IGFR2 binder | Quartetto yellow |
| | | | | | | | | | | 561 | Alexa Fluor 555 EGF | Quartetto green |
| | | | | | | | | | | 638 | Alexa Fluor 647 BMPR2 binder | Quartetto magenta |
| | | | | | | | | | | | CF680R I$\alpha$5$\beta$1 binder | Cyan |
| SF16 | SDCM | U2OS (37 °C) | 98.5 | 200 | 100 | 45 | 51 | 5 | 16 | 405 | Lyn-Cerulean | Duo intense cyan |
| | | | | | | | | | | 470 | MTS-mAzamiGreen | Red |
| | | | | | | | | | | | GTS-Citrine | Green |
| | | | | | | | | | | 555 | mCherry-KDEL | Circus yellow |
| | | | | | | | | | | 640 | iRFP670-SKL | Magenta |
| SF18 | OPM | *D. discoideum* (20 °C) | 111.6 | 750 | 20 | 0.5 | 27 | 2 | 40 | 488 | PIPpkgE-PH-EGFP | Duo intense cyan |
| | | | | | | | | | | 561 | LifeAct-mCherry | Duo intense yellow |
| SF20a | OPM | NIH 3T3 (37 °C) | 111.6 | 750 | 50 | 1 | 81 | 1 | 1 | 488 | Alexa Fluor 488 TfR binder | Black ink wash |
| | | | | | | | | | | | ATTO 514 IGF2R binder | |
| | | | | | | | | | | 638 | Alexa Fluor 647 BMPR2 binder | |
| | | | | | | | | | | | CF680R I$\alpha$5$\beta$1 binder | |
| SF20b | SDCM | NIH 3T3 (37 °C) | 110 | N/A | 120 | 1 | N/A | 2 | 121 | 488 | EGFP-WASH | Green |
| | | | | | | | | | | 638 | Alexa Fluor 647 TfR binder | Magenta |
| SF20c | SDCM | NIH 3T3 (37 °C) | 110 | N/A | 120 | 1 | N/A | 2 | 50 | 488 | EGFP-WASH | Green |
| | | | | | | | | | | 638 | Alexa Fluor 647 BMPR2 binder | Magenta |
| SF20d,e | SDCM | NIH 3T3 (37 °C) | 110 | N/A | 120 | 1 | N/A | 1 | 1 | 488 | EGFP-WASH | Black ink wash |
| | | | | | | | | | | | Alexa Fluor 488 TfR binder | |

[Supplementary Table 3 continued]

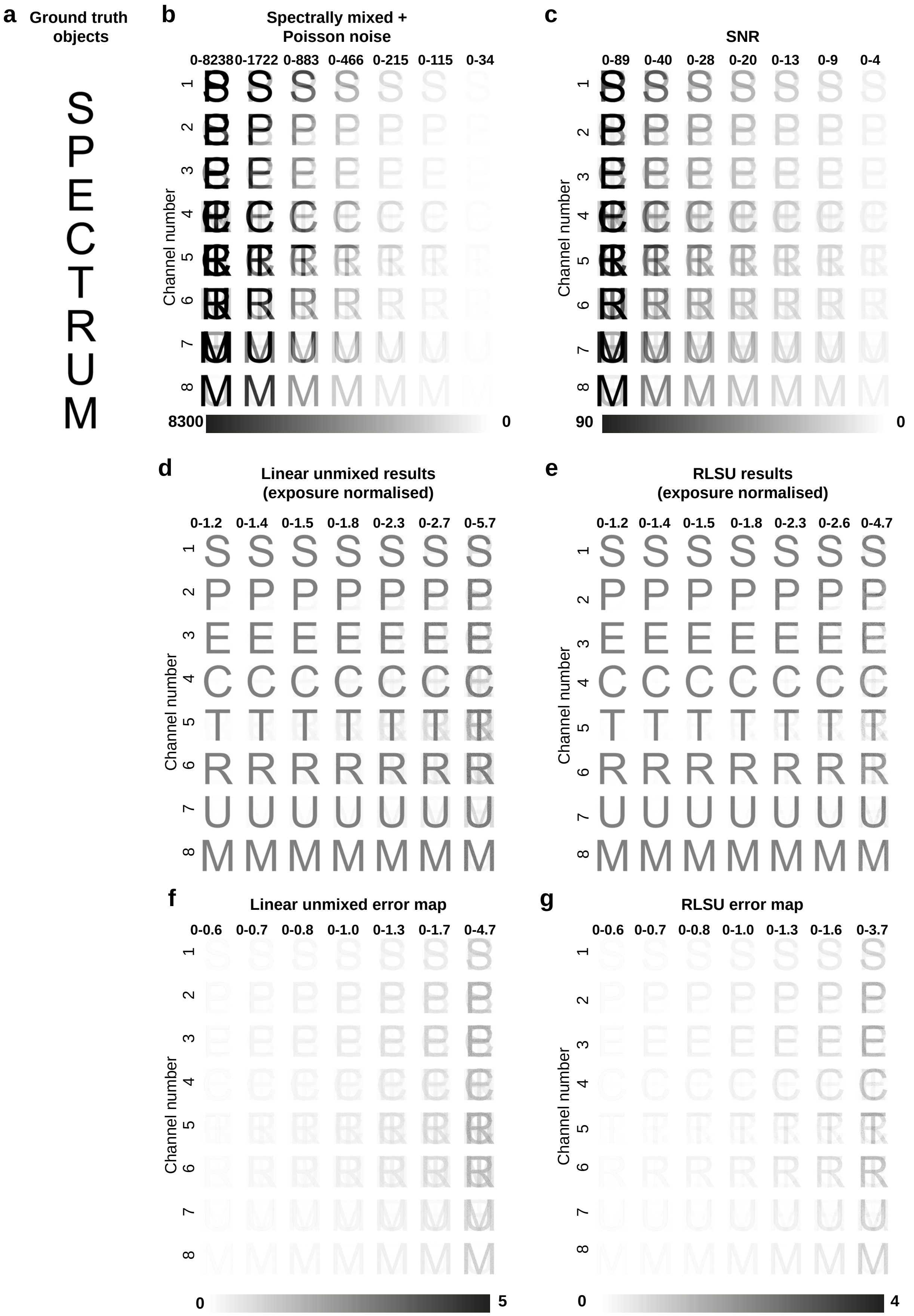


**Supplementary Figure 1: RLSU outperforms linear unmixing for unmixing simulated multispectral datasets with varying SNR.**
**a** Ground truth objects. **b** Mixed data incorporating Poisson (shot) noise, generated from (a) at different signal levels (high signal – low signal, right – left, respectively). This simulates the detection of mixed images with reducing exposure time. **c** SNR maps of data shown in (b). Linearly unmixed (**d**) and RLSU unmixed (**e**) results of data shown in (b). Results are normalised to the exposure time used to generate (b). **f-g** Error maps of data shown in (d-e). RLSU produces fewer errors than linear unmixing when reconstructing objects from mixed noisy images at every SNR tested. Data shown in grayscale inverted LUT with the range of pixel values shown numerically above each column.

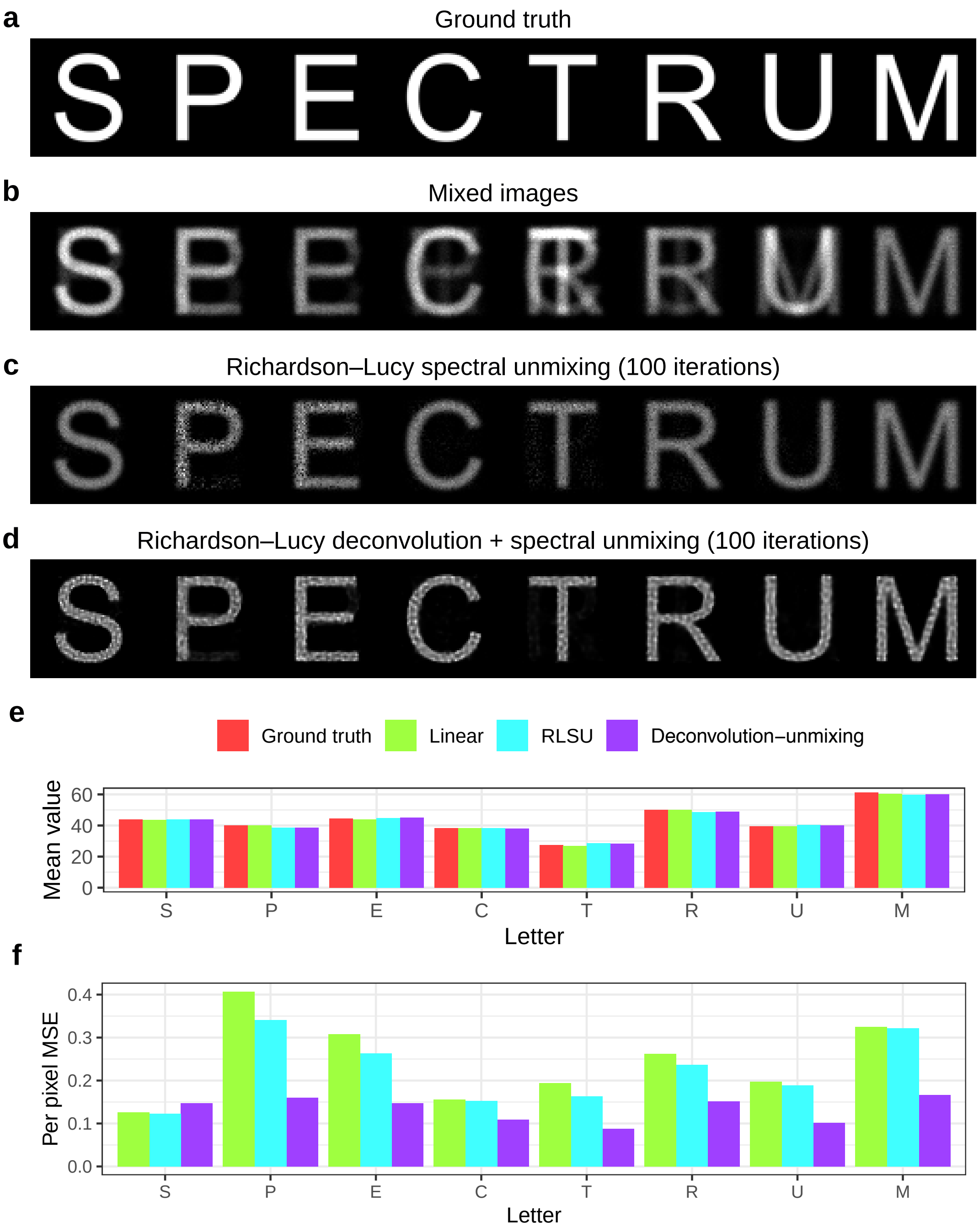


**Supplementary Figure 2: Richardson-Lucy deconvolution and spectral unmixing.**
**a** Ground truth objects, taken as letters from the word SPECTRUM. **b** Blurred, mixed noisy raw data, generated from (a). **c** Unmixed objects produced using RLSU without deconvolution after 100 iterations. **d** Unmixed objects produced using simultaneous Richardson–Lucy deconvolution-unmixing after 100 iterations. **e** Mean pixel values for each object (ground truth) or unmixed reconstructions resulting from linear unmixing, RLSU or simultaneous RL deconvolution-unmixing. **f** Per pixel mean squared error for unmixed reconstructions resulting from linear unmixing, RLSU or simultaneous RL deconvolution-unmixing.

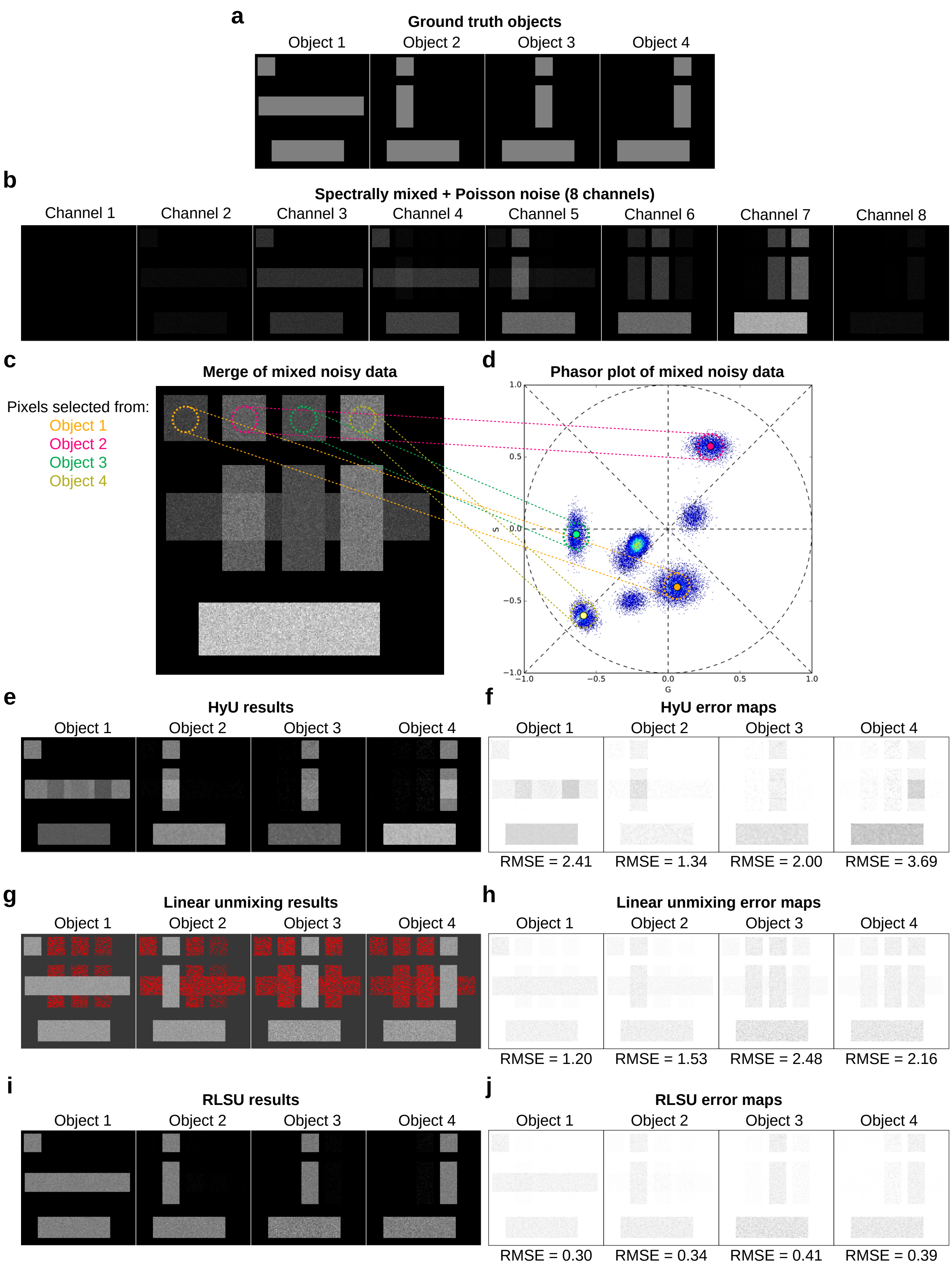


**Supplementary Figure 3: RLSU outperforms phasor-based unmixing (HyU) at unmixing simulated eight-channel multispectral data.** **a** Four ground truth objects. **b** Mixed, Poisson-noisy 8-channel data, generated from (a), designed to be similar to data acquired from our multispectral imaging module. **c** Overlay of data in (b). Coloured dashed circles represent ROIs in the mixed data with pixels deriving solely from object 1 (orange), 2 (magenta), 3 (green) or 4 (yellow). **d** Phasor plot of data in (b). Coloured dashed circles represent pixels associated with single-labelled objects in phasor space. **e** Unmixed object reconstructions generated using HyU. **f** Pixel-wise error map (inverted LUT) of HyU reconstructions, with root mean square error (RMSE) noted below. **g** Unmixed reconstructions using linear unmixing. Red pixels indicate negative pixel values. **h** Error map (inverted LUT) of linear unmixed reconstructions, with RMSEs below. **i** Unmixed object reconstructions generated using RLSU. **j** Error map (inverted LUT) of RLSU reconstructions, with RMSEs below.

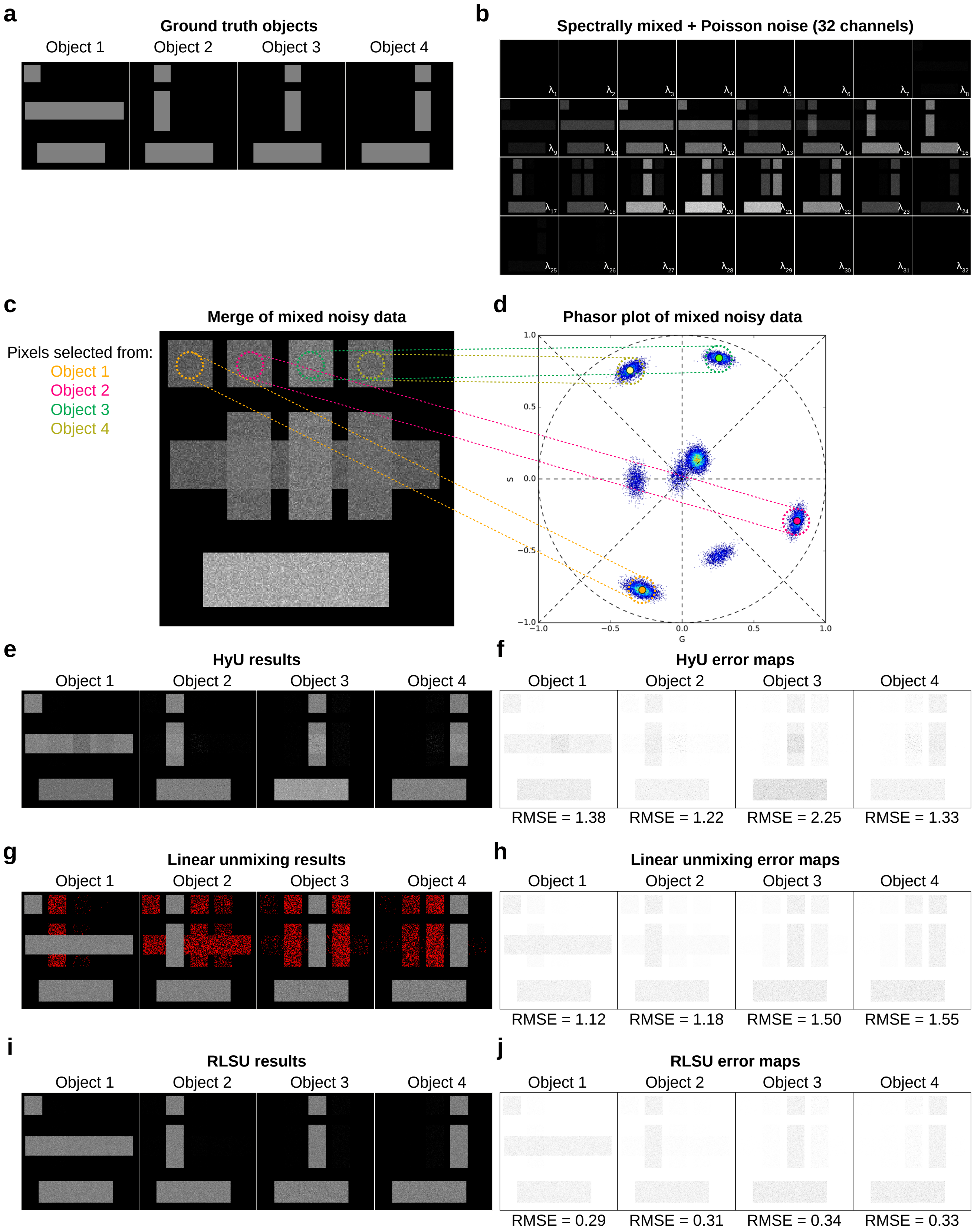


**Supplementary Figure 4: RLSU outperforms phasor-based unmixing (HyU) at unmixing simulated 32-channel multispectral data.** **a** Four ground truth objects. **b** Mixed, Poisson-noisy 32-channel data, generated from (a), designed to be similar to data acquired from Zeiss QUASAR detectors. **c** Overlay of data in (b). Coloured dashed circles represent ROIs in the mixed data with pixels deriving solely from object 1 (orange), 2 (magenta), 3 (green) or 4 (yellow). **d** Phasor plot of data in (b). Coloured dashed circles represent pixels associated with single-labelled objects in phasor space. **e** Unmixed object reconstructions generated using HyU. **f** Pixel-wise error map (inverted LUT) of HyU reconstructions, with root mean square error (RMSE) noted below. **g** Unmixed reconstructions using linear unmixing. Red pixels indicate negative pixel values. **h** Error map (inverted LUT) of linear unmixed reconstructions, with RMSEs below. **i** Unmixed object reconstructions generated using RLSU. **j** Error map (inverted LUT) of RLSU reconstructions, with RMSEs below. Error maps are shown in inverted LUT with darker pixels representing larger errors.

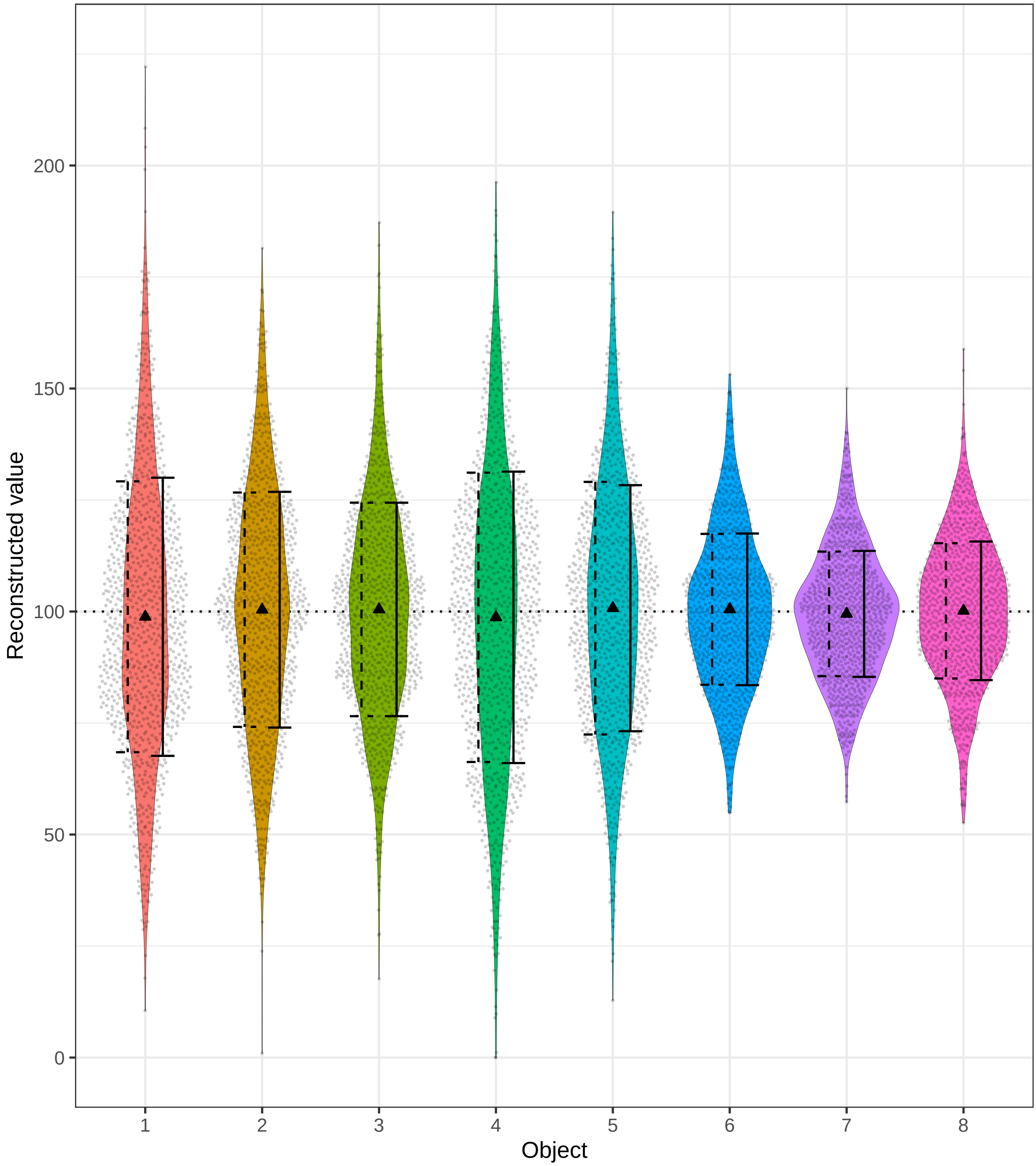


**Supplementary Figure 5: Cramér–Rao lower bound analysis.**
Distribution of RLSU reconstructed values from 1000 independent sets of Poisson-noisy mixed signals for eight ground truth objects of value 100 (see Supplementary Note C for details). Black triangles show the mean of the 1000 values, correctly sitting around 100. Dashed error bars show the square root of the appropriate Cramér–Rao lower bound, with the standard deviation of the 1000 independent reconstructions shown as a solid error bar.

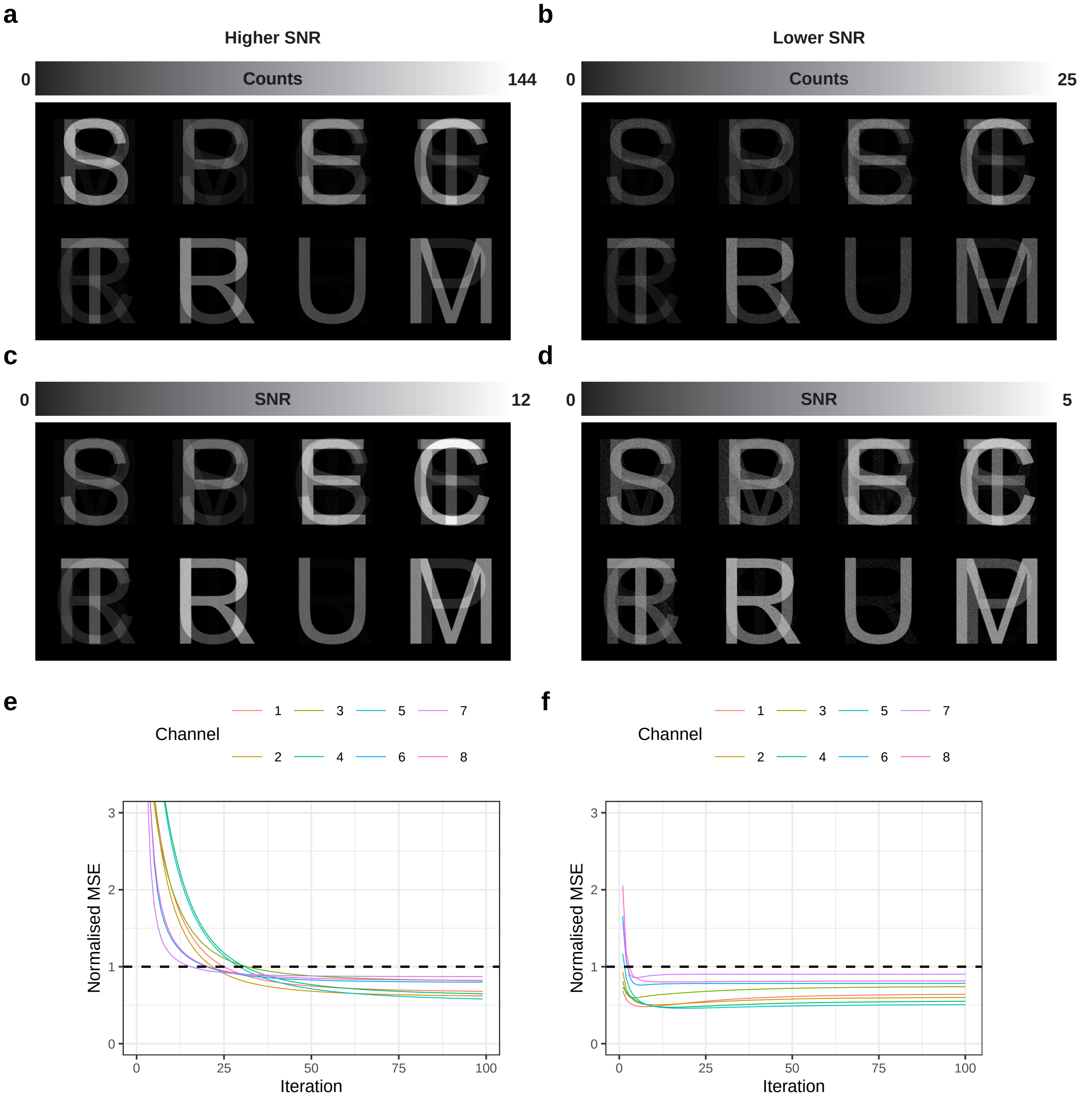


**Supplementary Figure 6: RLSU outperforms linear unmixing for unmixing both low and high SNR simulated multispectral datasets.** **a** Simulated mixed noisy images of higher SNR data. **b** Simulated mixed noisy images of lower SNR data. **c** SNR map of (a). **d** SNR map of (b). **e** Normalised mean squared error of unmixed results at different iterations of RLSU versus linear unmixing for unmixing higher SNR data (a). Normalised mean squared error (RLSU) shown for all eight channels in coloured lines, whilst the linear unmixing result is indicated by dashed horizontal black line. **f** Normalised mean squared error of unmixed results at different iterations of RLSU versus linear unmixing for unmixing lower SNR data (b). Normalised mean squared error (RLSU) shown for all eight channels in coloured lines, whilst the linear unmixing result is indicated by dashed horizontal black line. Note that in both cases within 100 iterations RLSU converges to a solution and outperforms linear unmixing in all eight channels.

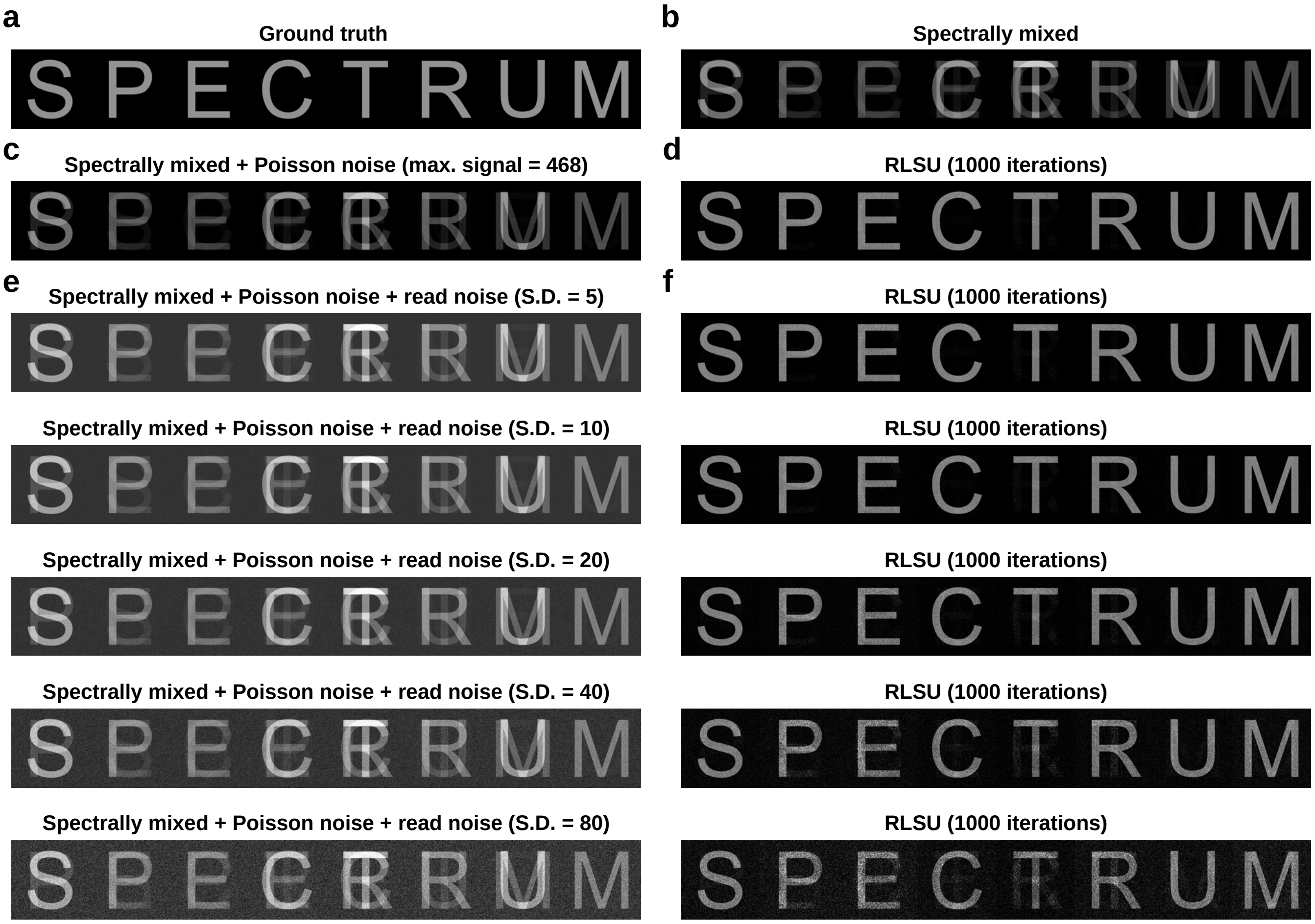


**Supplementary Figure 7: RLSU unmixing of simulated data incorporating Poisson (shot) and read noise.**
**a** Eight simulated ground truth objects corresponding to different fluorophores. **b** Spectrally mixed, noise-free data. **c** Spectrally mixed data with Poisson (shot) noise incorporated, and a maximum realised signal of 468 counts. **d** RLSU reconstruction of ground truth objects from data in (c). **e** Data from (c) with simulated camera background (100 counts) and increasing amounts of Gaussian-distributed read noise. **f** RLSU reconstructions from data in (e). 100 counts were removed prior to unmixing to remove simulated 'camera background'. Results are only noticeably different from the zero-read-noise case at the standard deviation = 20 level (i.e. when the variance of the read noise is comparable to the variance of the Poisson (shot) noise), a much higher level of read noise than found in modern sensors.

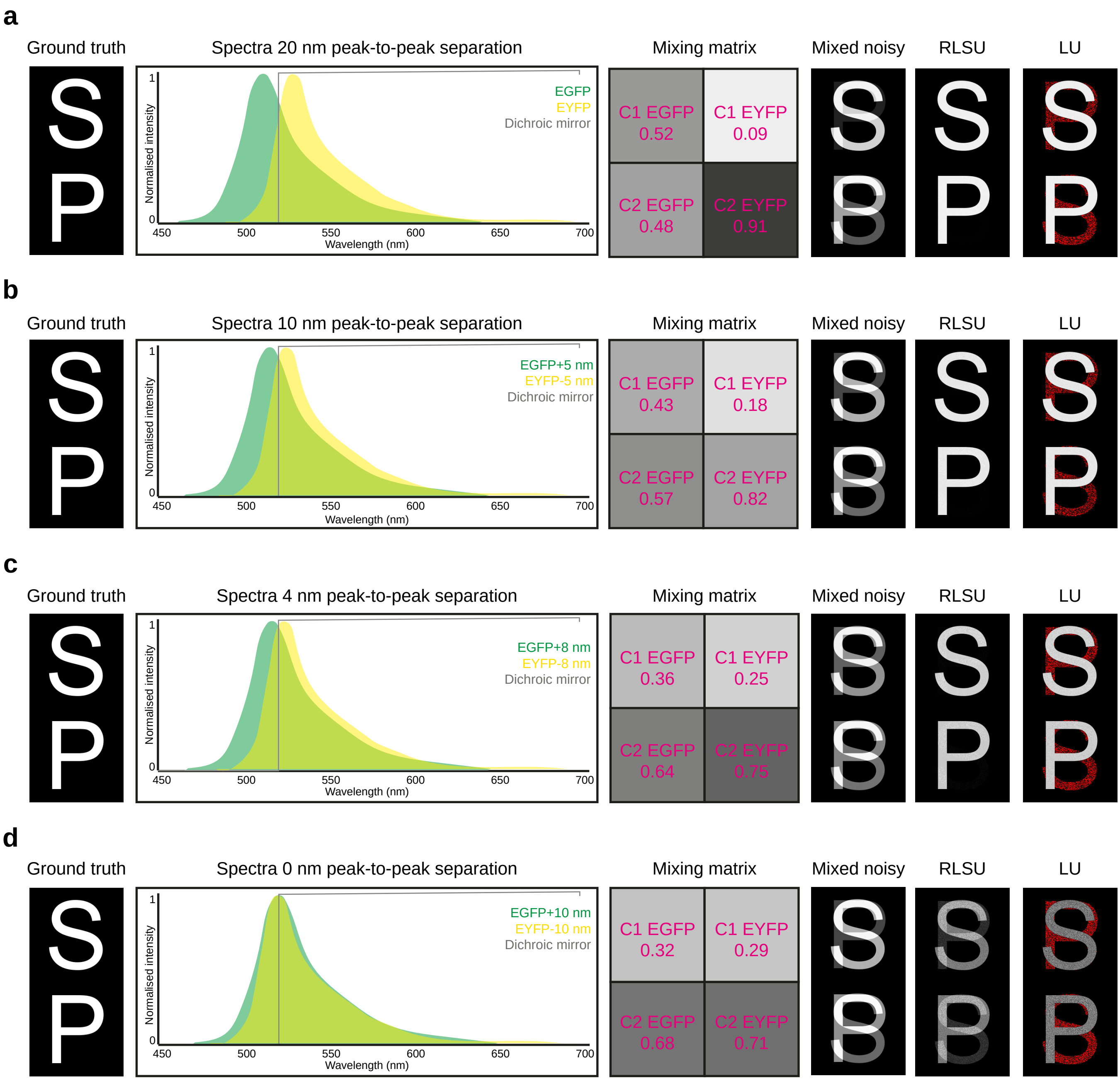


**Supplementary Figure 8: Comparison between RLSU and LU for unmixing simulated 2-channel data of increasing spectral overlap.** **a** Ground truth, spectra, mixing matrix, mixed images, RLSU result and LU result for 20 nm peak-to-peak spectral separation. **b** Ground truth, spectra, mixing matrix, mixed images, RLSU result and LU result for 10 nm peak-to-peak spectral separation. **c** Ground truth, spectra, mixing matrix, mixed images, RLSU result and LU result for 4 nm peak-to-peak spectral separation. **d** Ground truth, spectra, mixing matrix, mixed images, RLSU result and LU result for 0 nm peak-to-peak spectral separation. See Supplementary Note D.

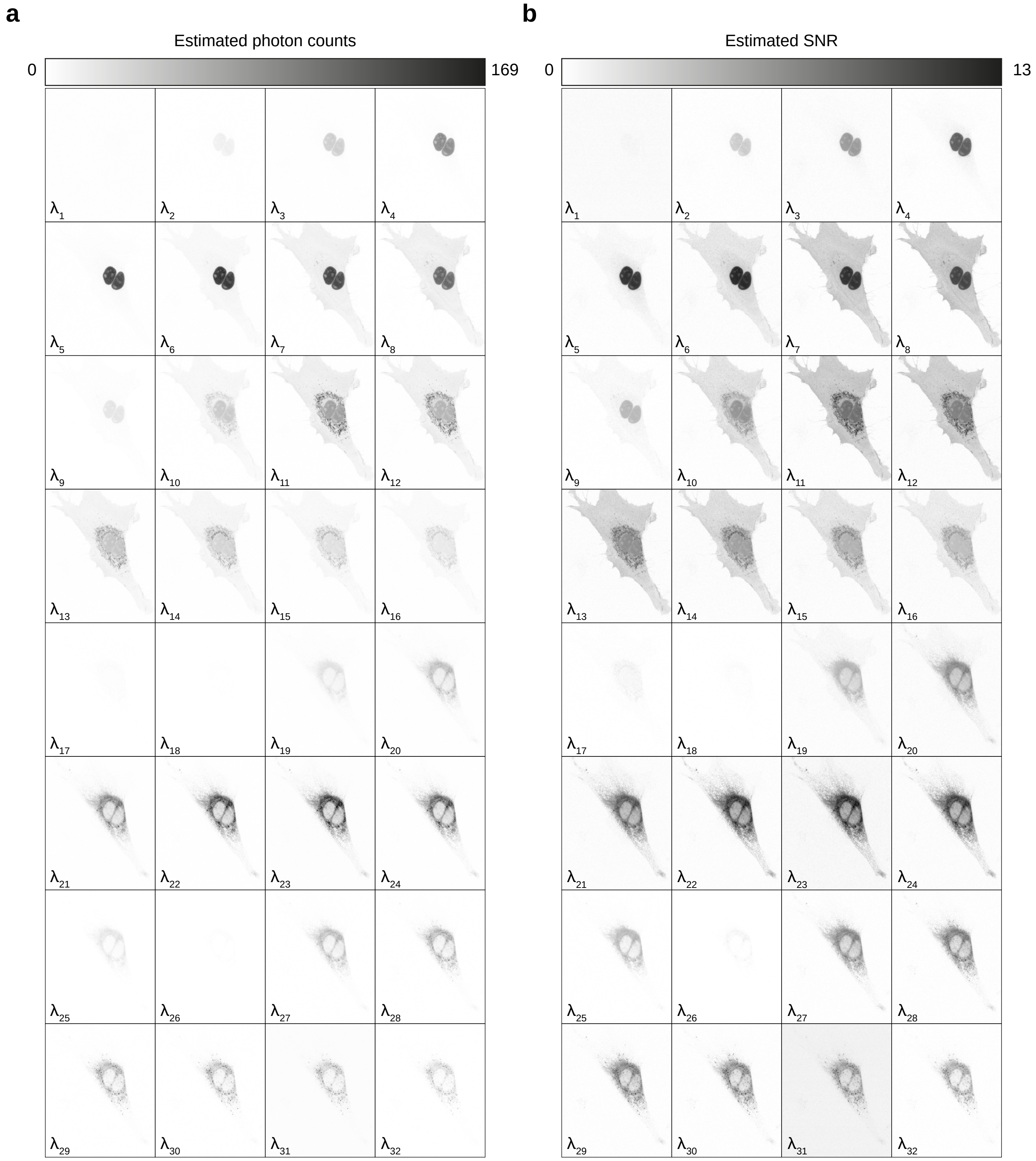


**Supplementary Figure 9: Estimated SNR of Zeiss QUASAR data.**
**a** Raw 32 channel data of a live U2OS cell transfected with ColorfulCell. Estimated photon counts are shown for each channel. **b** Estimated SNR values for data in (a) shown for each channel. An inverted LUT is used with darker pixels representing higher SNR.

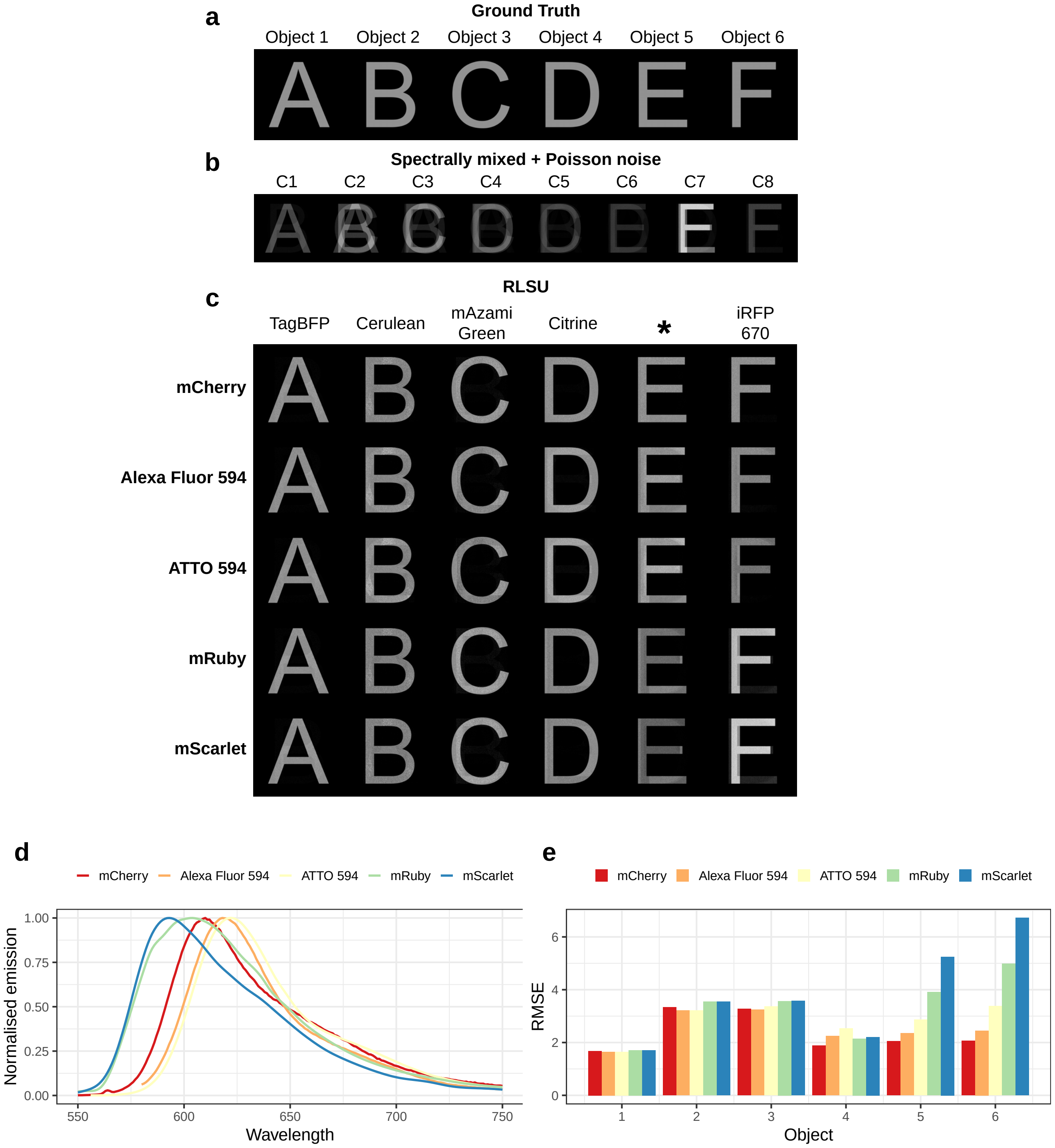


**Supplementary Figure 10: Assessment of RLSU performance when using incorrect mixing matrices.**
**a** Ground truth objects. **b** Spectrally mixed data with Poisson (shot) noise incorporated. Data was mixed by a mixing matrix related to the following fluorophores: TagBFP, Cerluean, mAzami Green, Citrine, mCherry and iRFP 670. **c** Data in (b) was unimixed using RLSU and different mixing matrices. The top row was reconstructed after unmixing with the correct mixing matrix. The other rows were reconstructed after unmixing with incorrect mixing matrices, where mCherry was substituted with Alexa Fluor 596, ATTO 594, mRuby and mScarlet, respectively. This was done to assess how well RLSU could reconstruct ground truth objects when unmixing with erroneous mixing matrices. **d** Emission spectra of mCherry and other fluorophores used to form mixing matrices to produce unmixed data in (b). **e** Root mean squared errors associated with reconstructing ground truth objects after unmixing with the correct mixing matrix, and erroneous mixing matrices where mCherry was substituted for other fluorophores.

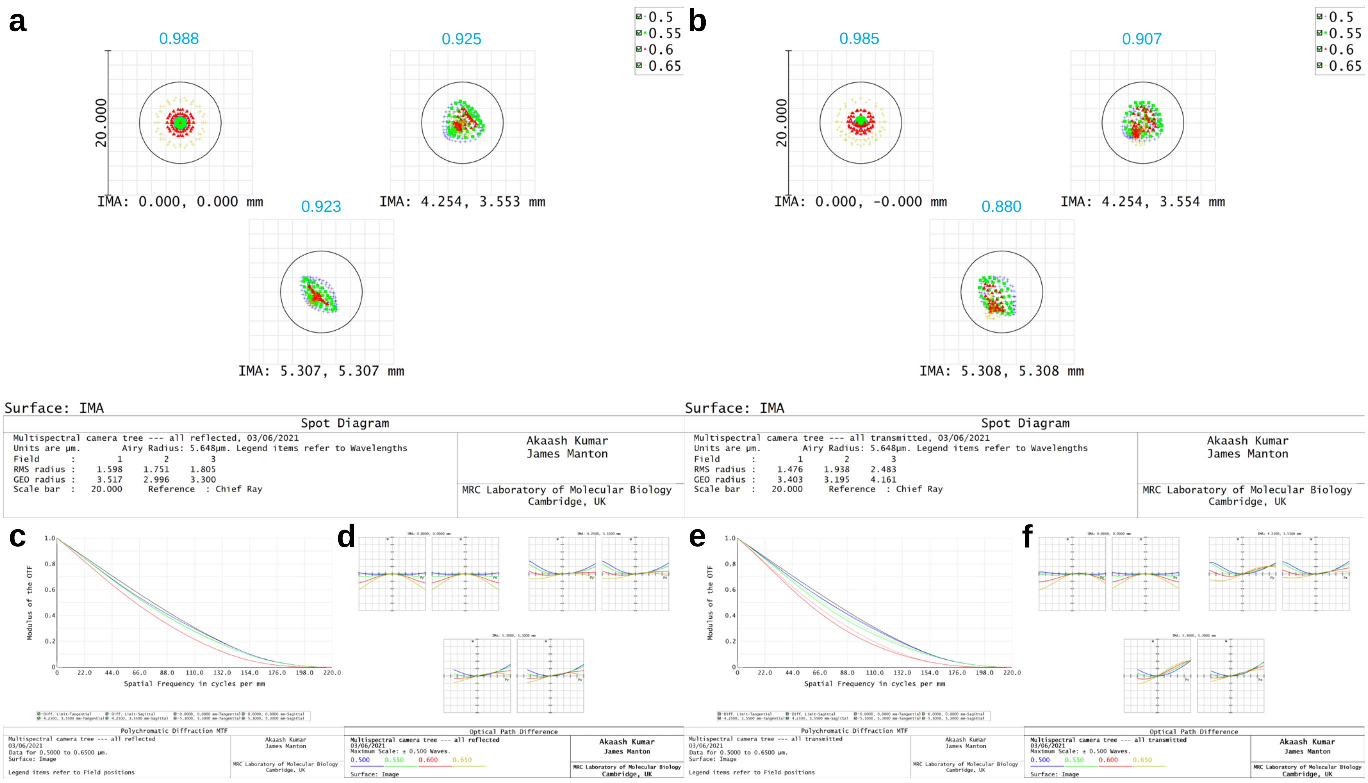


**Supplementary Figure 11: Raytracing analysis of eight channel imaging hardware.**
**a–b** Spot diagrams for central field, edge of camera chip and edge of tube lens image circle for (a) path in which light is reflected off all dichroics and (b) path in which light transmitted through all dichroics. Solid line shows Airy radius, demonstrating diffraction-limited performance across all fields for 500 nm to 650 nm. Cyan numbers indicate Strehl ratios. **c** Modulation transfer functions for reflected path. **d** Optical path difference plots for reflected path. **e** Modulation transfer functions for transmitted path. **f** Optical path difference plots for transmitted path.

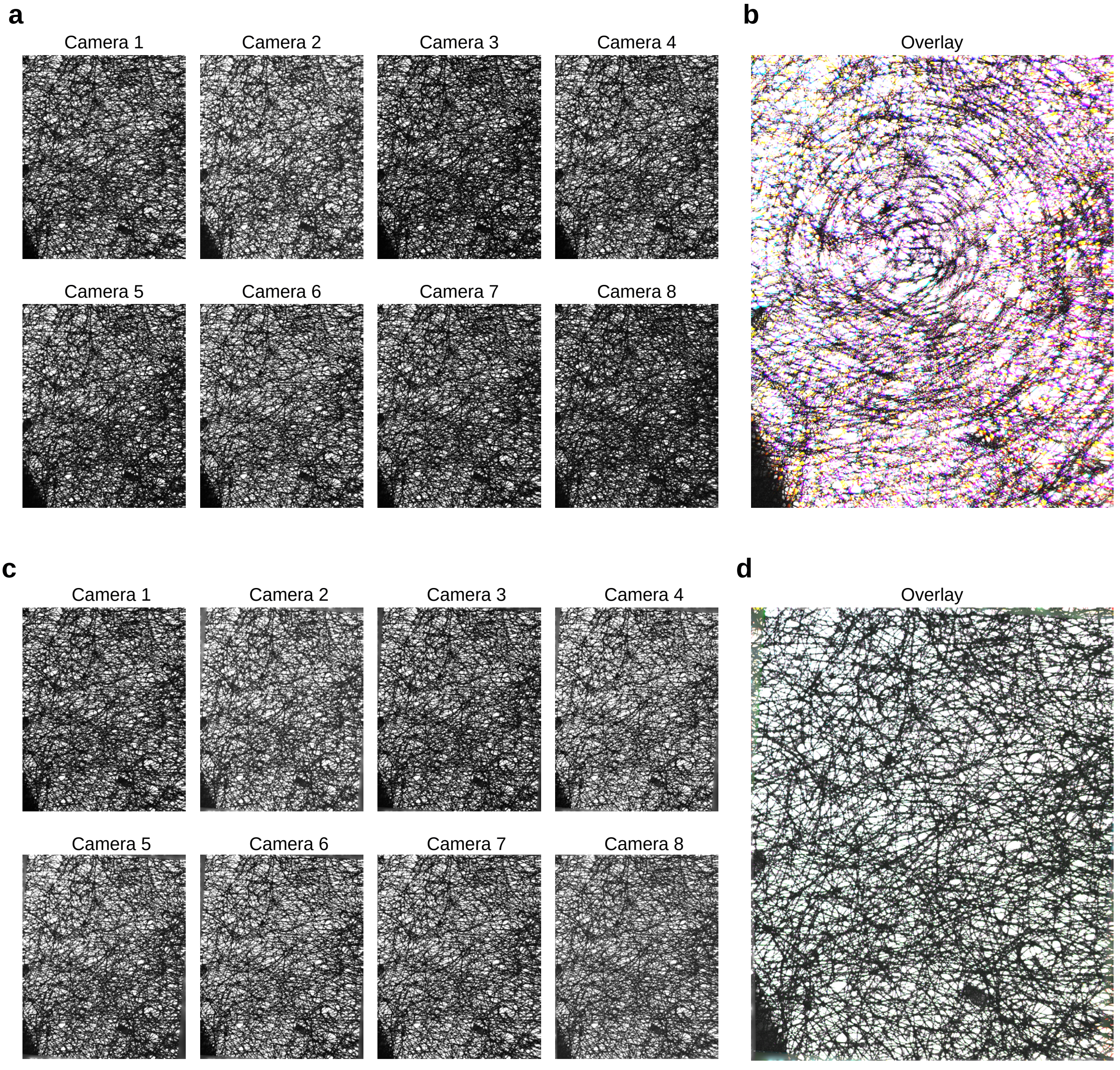


**Supplementary Figure 12: Computational registration using lens tissue images.**
**a** Images aquired by all 8 cameras of lens tissue placed in the microscope image plane. **b** Overlay of the data shown in a. **c** Data from (a), after computational registration. **d** Overlay of the data post computational registration.

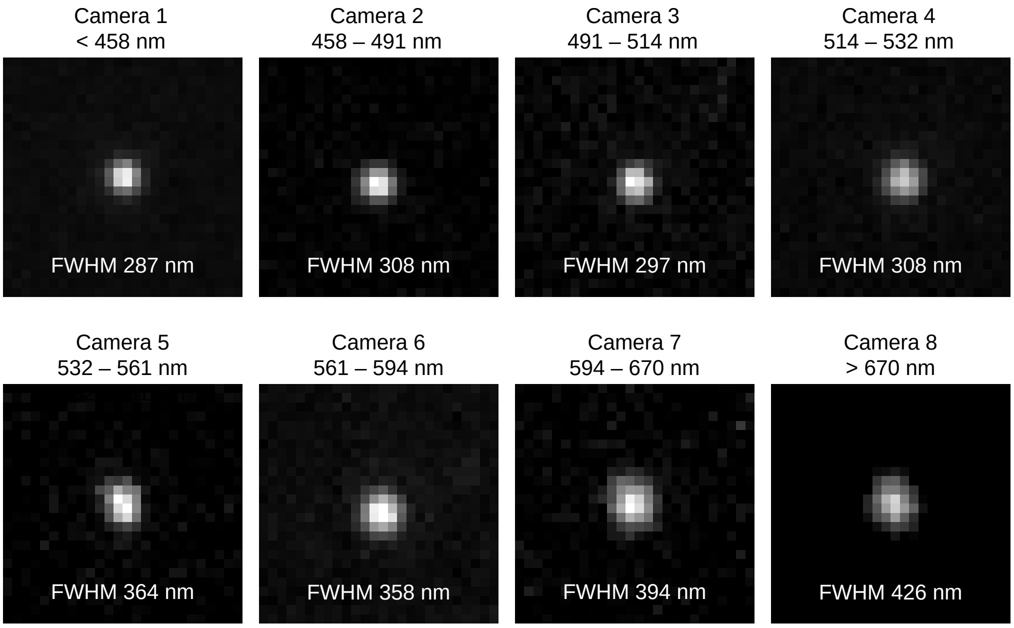


**Supplementary Figure 13: Multispectral spinning disk microscope point spread functions.**
Point spread functions and full-width-half-maximums displayed for the eight cameras on the multispectral spinning disk microscope used in this study. Pixel pitch = 98.5 nm/px.

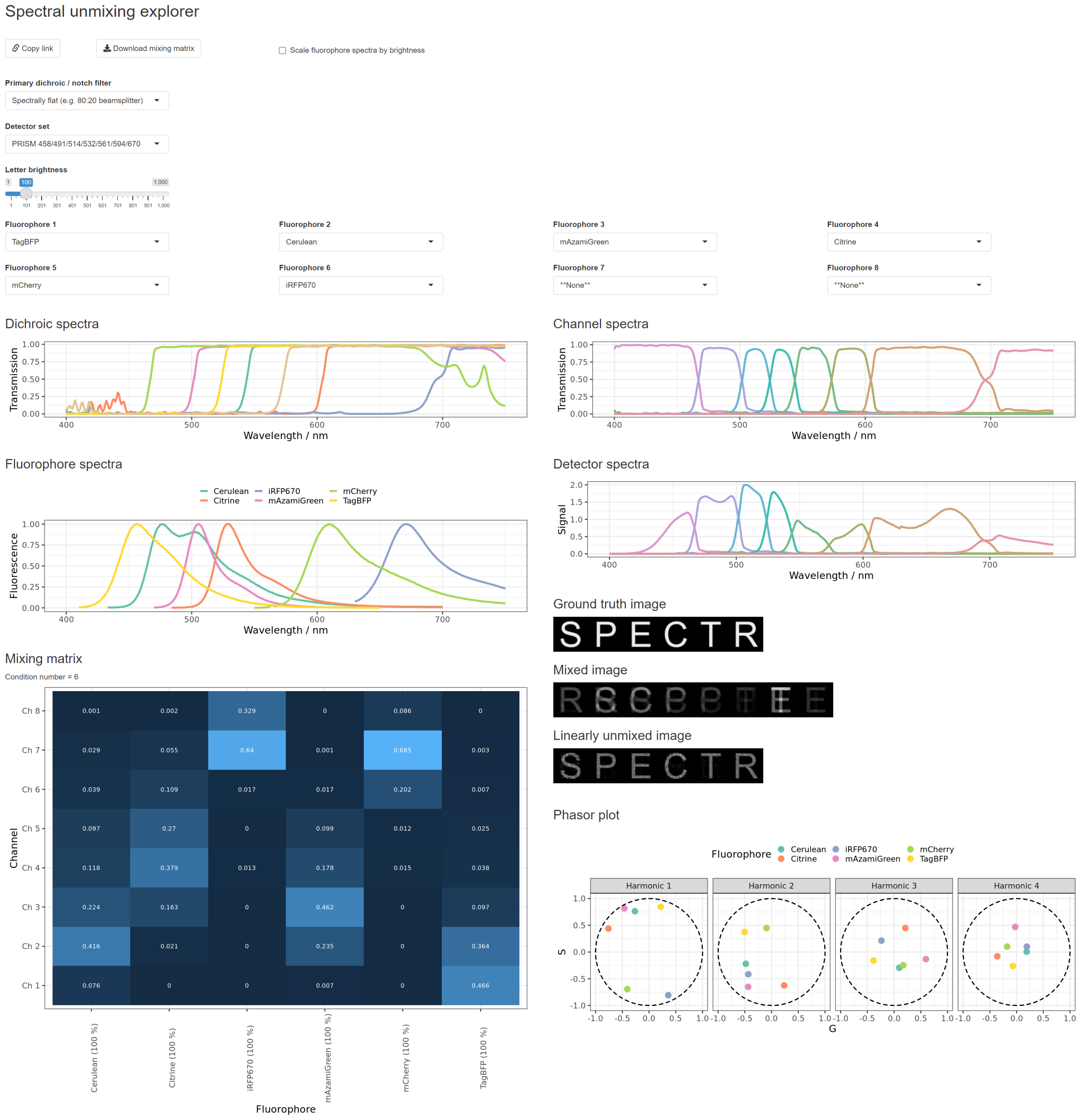


**Supplementary Figure 14: Screenshot of web application developed to facilitate the choice of fluorophores and the computation of unmixing matrices.**

Screenshot of web application hosted at beryl.mrc-lmb.cam.ac.uk/calculators/spectral_unmixing/. Users select fluorophores, using spectral information taken from fpbase.org, and select an appropriate dichroic set. The application then plots the relevant spectra, calculates a mixing matrix (shown as a blue heatmap) and produces simulated raw and linearly unmixed data (to the right of the heatmap). These simulated results, along with the condition number (printed under the mixing matrix heading) and phasor plots (bottom right) help the user check that their fluorophore and dichroic selection is appropriate. In particular, a small condition number, well-dispersed points on the phasor diagrams, and well-unmixed simulated data are desired. If the user wishes to share their selection with another user, a button is provided to generate a link which, when clicked, will preload the application with all the relevant data.

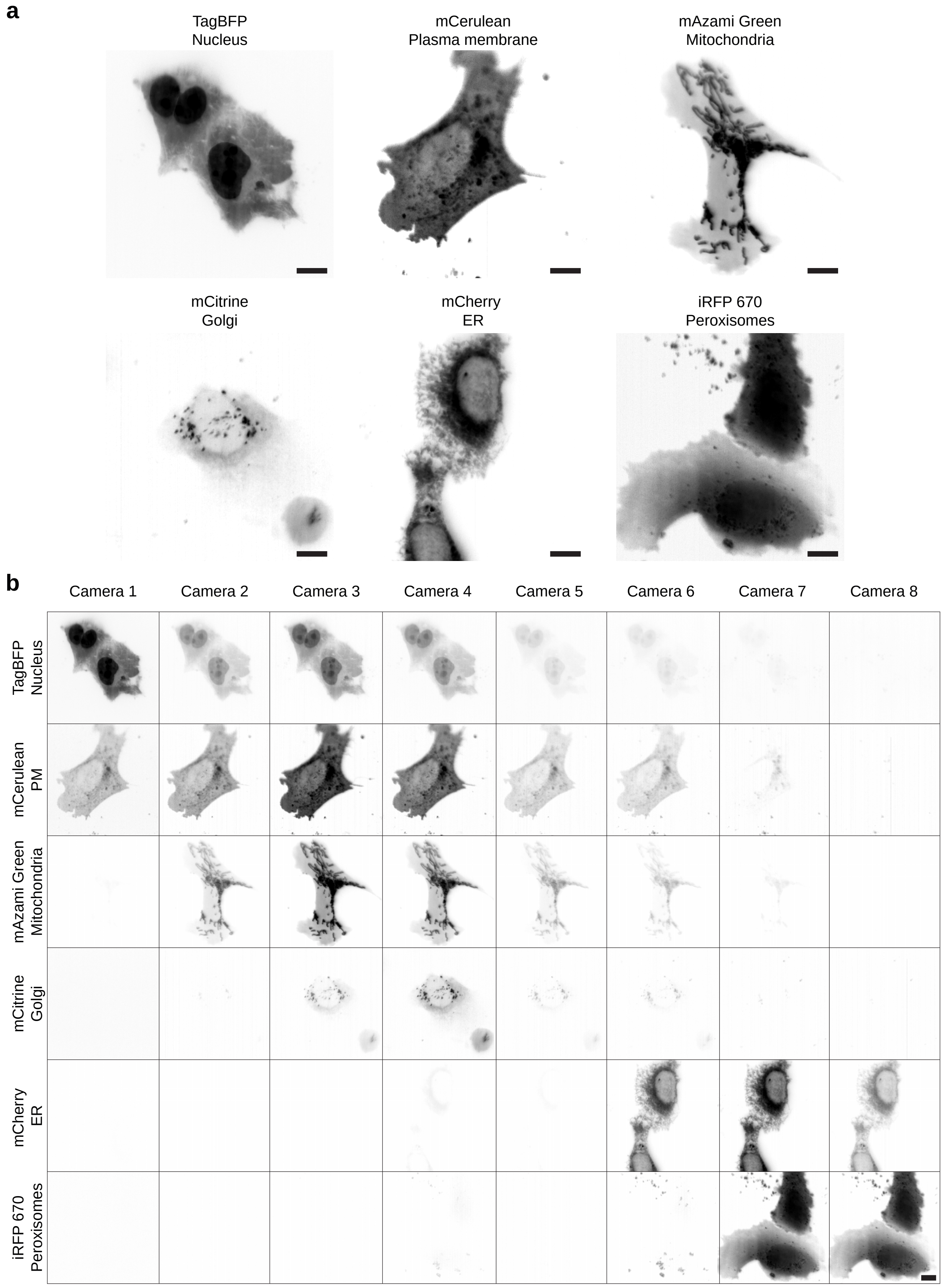


**Supplementary Figure 15: Single-labelled controls of the individual fluorescent markers encoded by ColorfulCell.**
**a** Maximum intensity projections of U2OS cells transfected with individual fluorescent markers present in the ColorfulCell plasmid. This provides definitive examples of what each of the components should look like post-unmixing. Data acquired using the multispectral OPM followed by registering and summing the data from the eight channels (summed intensity projection). **b** Signals across all 8 cameras are shown for each of the cells presented in (a). These signals can be measured for each marker to determine a measured mixing matrix for subsequent RLSU unmixing of ColorfulCell data. Scale bars = 10 µm.

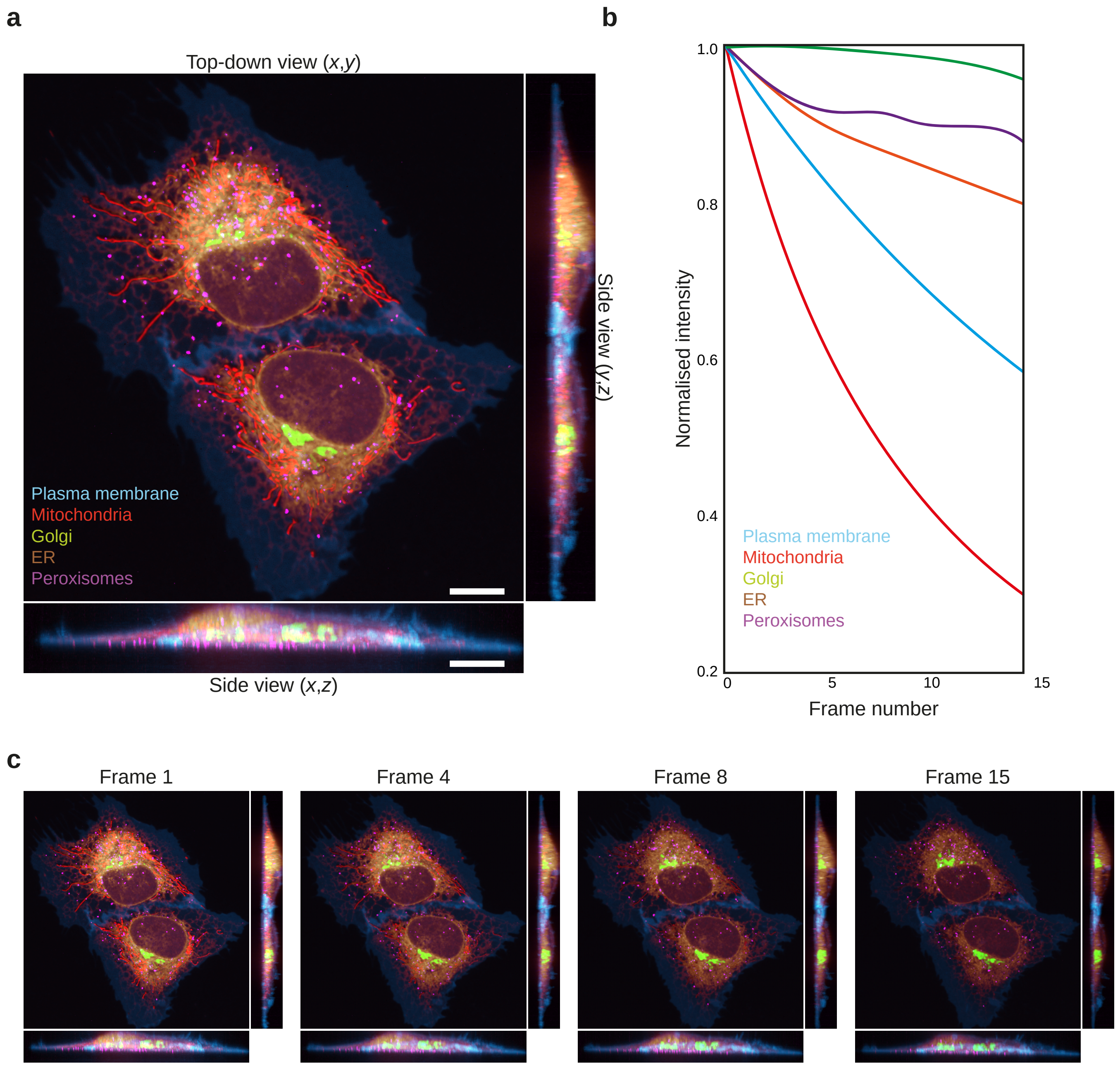


**Supplementary Figure 16: Multispectral spinning disk confocal microscope causes photodamage during fast volumetric acquisitions.** **a** Live U2OS cells transfected with the ColorfulCell plasmid were imaged by multispectral confocal spinning disk followed by RLSU unmixing. Images represent orthogonal maximum intensity projections of confocal volumes in indicated axis (51 planes ; 200 nm increment). **b** Normalised intensity over time for signals in shown in (a). **c** Timelapse volumetric aquisition showing progressive signal bleaching and cellular photodamage (*e.g.* cell shrinking). see also Supplementary Video 6.

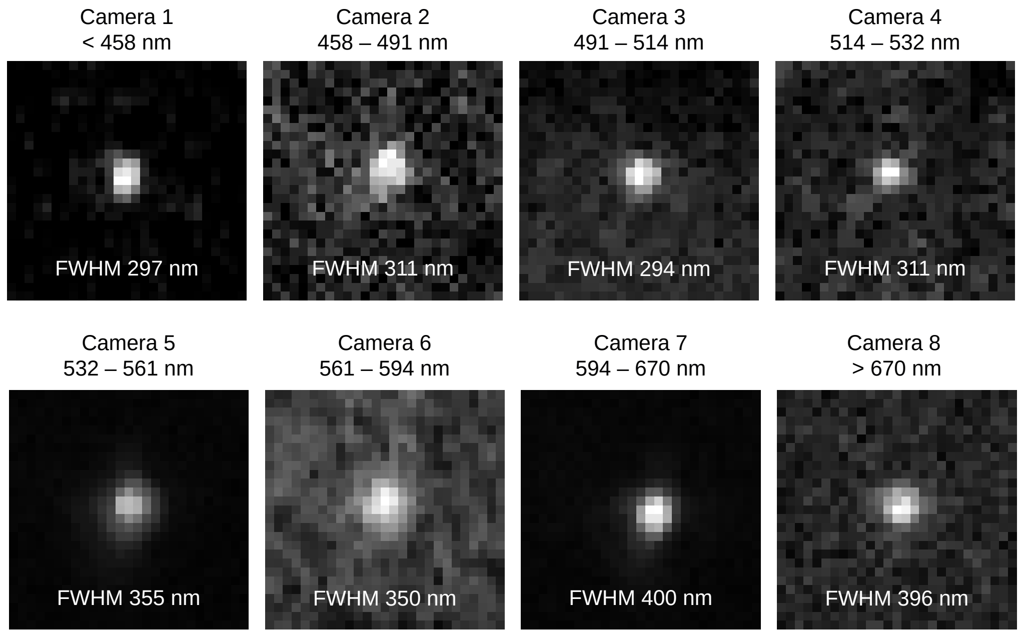


**Supplementary Figure 17: Multispectral oblique plane light sheet microscope point spread functions.**
Point spread functions and full-width-half-maximums displayed for the eight cameras on the multispectral oblique plane light sheet microscope. Pixel pitch = 111.6 nm/px.

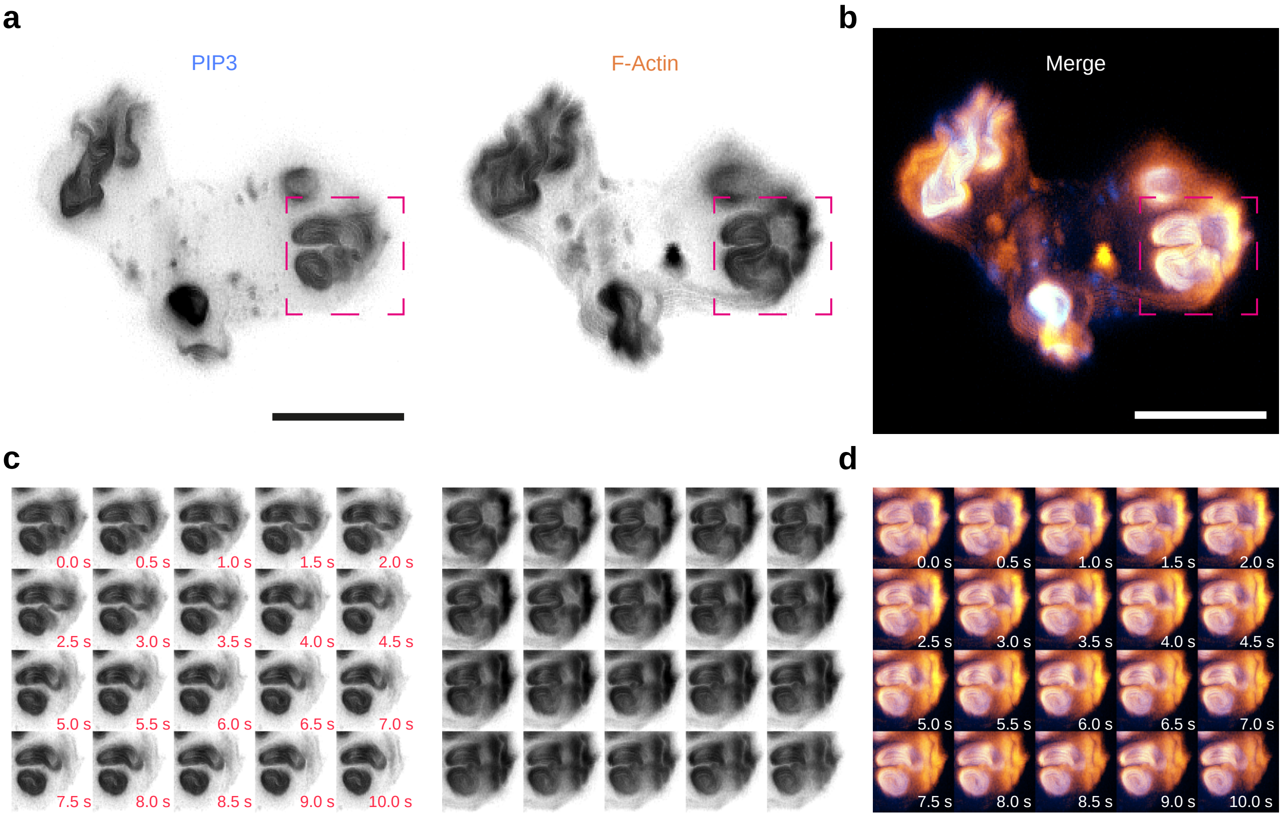


**Supplementary Figure 18: Fast volumetric imaging of live *Dictyostelium discoideum* cells.**
**a** Maximum intensity projections of live *Dictyostelium discoideum* cell expressing EGFP-fused to a PIP3 reporter and LifeAct-mCherry. Data acquired using the multispectral oblique plane light sheet microscope at 2 Hz. See also Supplementary Video 11. **b** Colour merge of data in (a). **c** Frames from timelapse (cyan dashed box in (a)) showing macropynocitic cup closure. Time interval 0.5 s between frames. **d** Colour merge of (c). Scale bars = 10 µm.

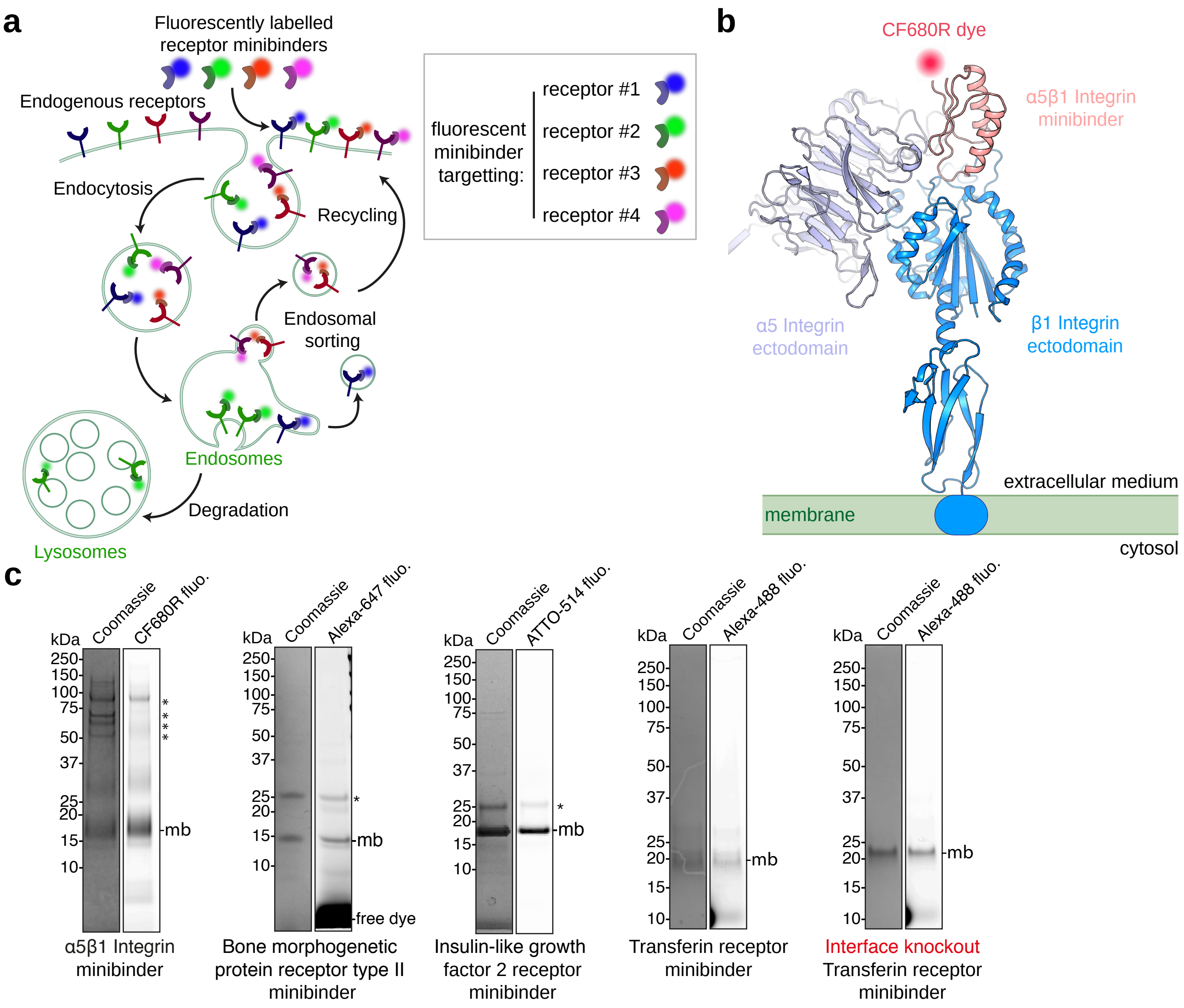


**Supplementary Figure 19: Biochemical characterisation of de novo designed receptor minibinders.**
**a** Principle of the experiment. Cells are incubated with a panel of de novo designed fluorescent minibinders targetting different endogenous receptors on ice to block endocytosis. After 30 minutes, cells are washed extensively and incubated with warm serum-containing medium to trigger endocytosis of the receptors bound to their cognate fluorescent minibinders. The fluorescence signal of the different minibinders can then be used to directly image the sorting of the different receptors into different organelles, for instance between early/sorting endosomes and late endosomes. Different organelles are expected to gradually have different ratios of the different minibinders if sorting occurs. On the contrary, if no sorting occurs, or if the minibinders enter the cell non-specifically by fluid-phase endocytosis, we expect all compartments to consistently have the same ratios of the different minibinders. **b** Example of de novo designed fluorescent minibinder against integrin $\alpha5\beta1$. Note that the minibinder is monovalent, so does not dimerise the receptor, and that the reactive cysteine used to functionalise the minibinder with the organic dye CF680R is positioned away from the surface binding to the receptor. **c** SDS-PAGE gel of indicated receptor minibinder followed showing in-gel fluorescence and Coomassie blue staining. Asterisk indicates contaminants (usually leftover GST from the purification).

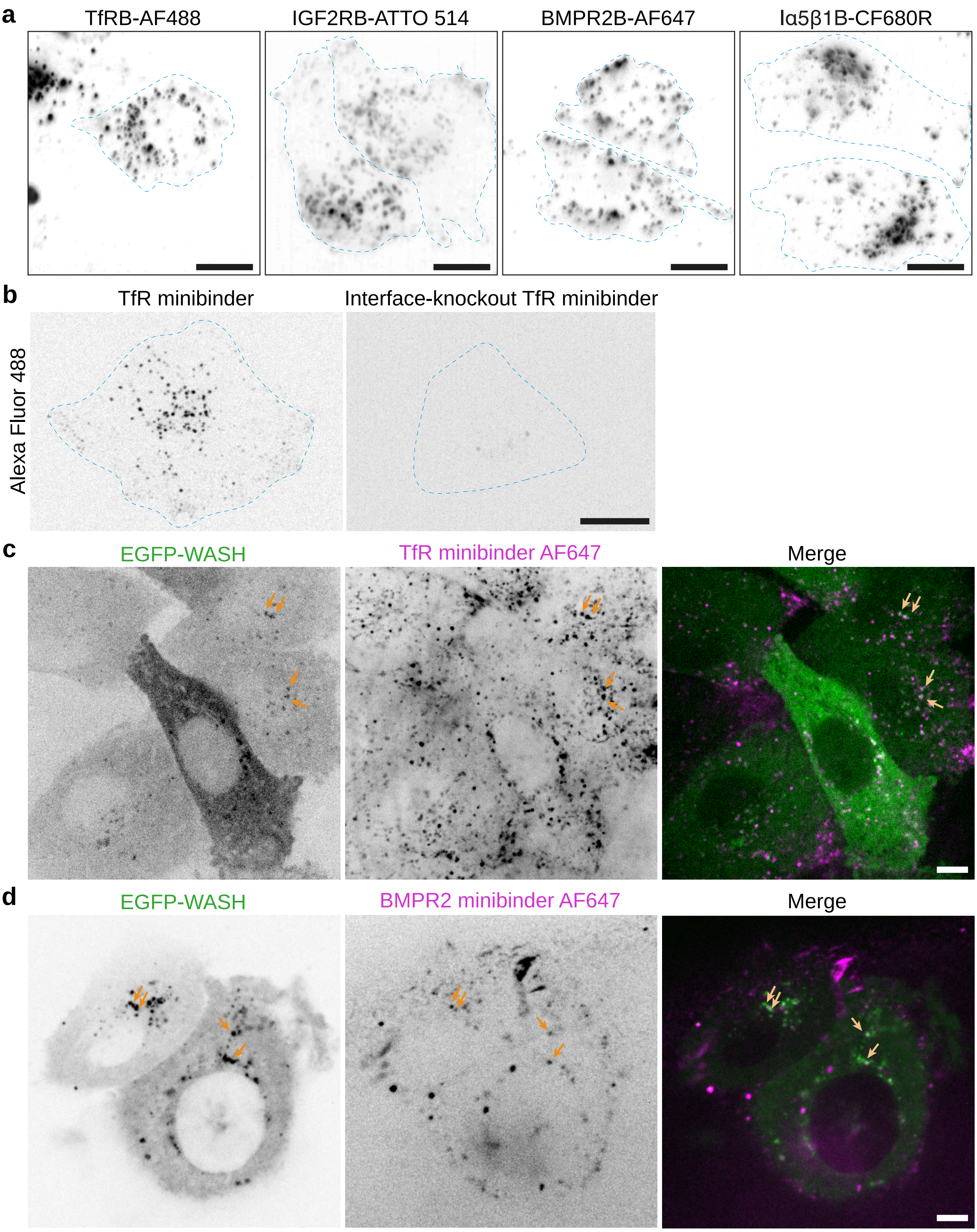


**Supplementary Figure 20: Validation of de novo designed receptor minibinders for imaging of receptor trafficking in live cells.**
**a** HeLa Kyoto cells incubated with indicated fluorescent minibinders (cyan dashed line indicates cell boundaries). Images correspond to maximum intensity projections from multispectral OPM data. This is a control related to Figure 6a showing all four fluorescent minibinders can be uptaken by cells individually. **b** HeLa Kyoto cells incubated with Alexa Fluor 488 labelled, TfR minibinder (left) or a respective interface knockout mutant (right), were imaged live with spinning disk confocal microscopy. Images correspond to single confocal planes (cyan dashed line indicates cell boundaries). Note that no signal is observed for the interface knockout mutant [24], suggesting that the fluorescent signal observed in the wildtype minibinder corresponds to minibinder bound to its target transferrin receptor, and not non-specific fluid phase uptake of the fluorescent minibinder. **c** NIH 3T3 cells stably expressing EGFP-WASH and incubated with TfR minibinder Alexa Fluor 647 were imaged live by spinning disk confocal microscopy (single confocal plane shown) . Note the colocalisation between EGFP and Alexa Fluor 647 (arrow) suggesting that the TfR minibinder is trafficking through WASH-positive early/sorting endosomes. **d** NIH 3T3 cells stably expressing EGFP-WASH and incubated with BMPR2 binder Alexa Fluor 647 were imaged live by spinning disk confocal microscopy (single confocal plane shown) . Note the colocalisation between EGFP and Alexa Fluor 647 (arrow) suggesting that the BMPR2 minibinder is trafficking through WASH-positive early/sorting endosomes. Scale bars = 10 µm.

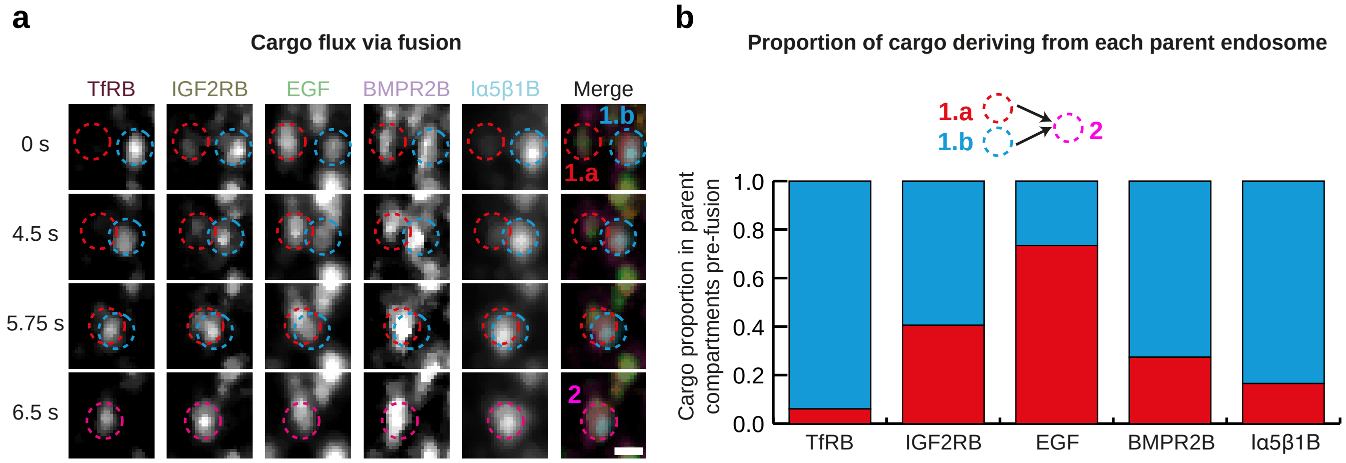


**Supplementary Figure 21: Multispectral light sheet imaging of endosomal fusion using de novo designed receptor minibinders.**
**a** Frames from Supplementary Video 14 showing cargo flux during endosomal fusion. A maximum intensity projection is shown. Individual channels and colour merge are shown. Red and cyan dashed circles indicate two endosomes, 1.a and 1.b respectively, which fuse to form endosome 2, indicated by a magenta dashed circle. Scale bar = 1 μm. **b** Quantification of the event shown in (a) to show the proportion of cargo in endosome 2 deriving from endosomes 1.a and 1.b.

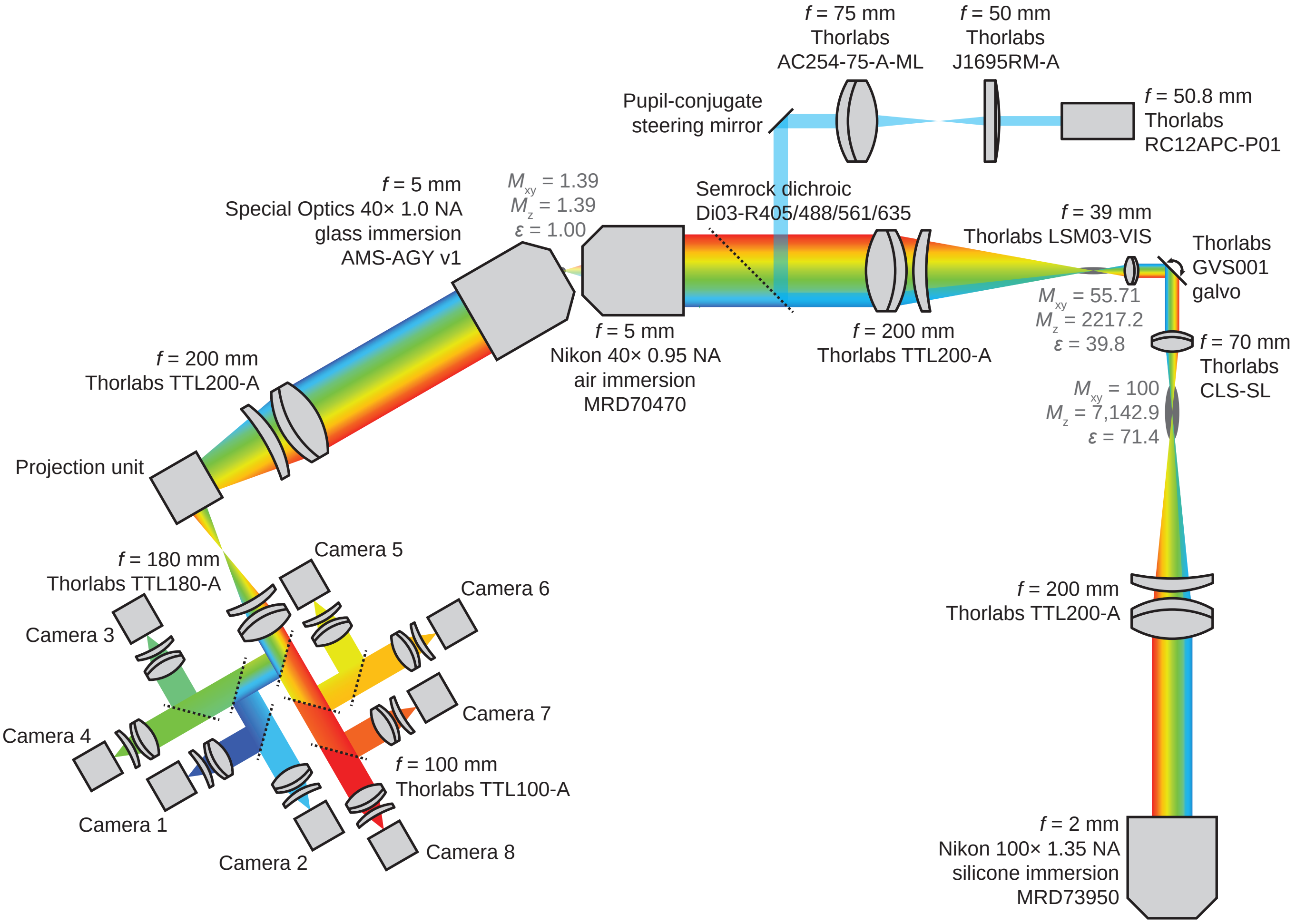


**Supplementary Figure 22: Optical diagram of the multispectral OPM.**
Fluorescence enters the 100× 1.35 NA objective (bottom right) and is directed through a tube lens and scan lens onto a galvanometric mirror, which scans the image plane (and light sheet) through the sample. This light then passes through another scan lens, tube lens and 40× 0.95 NA objective lens into the remote refocussing volume. Here, both the lateral ($M_{xy}$) and axial magnifications ($M_z$) are set to be equal ($\epsilon = 1.00$), and approximately equal to the refractive index of the sample, 1.39. This ensures that an aberration-free 3D volume is formed, a tilted plane of which is then imaged by the solid-immersion 40× 1.0 NA lens and tube lens onto the input plane of the multispectral camera system. A projection unit, consisting of another galvanometric mirror synchronised to the one moving the light sheet and image plane, is inserted just before the input image plane of the multispectral camera system to facilitate projection imaging. For illumination, laser light is directed from a reflective collimator through a cylindrical lens and achromatic doublet onto a dichroic mirror which couples the light into the main optical path, propagating in the opposite direction to the fluorescence light. A pupil-conjugate steering mirror facilitates alignment of the sheet of illumination to the tilted focal plane of the instrument.

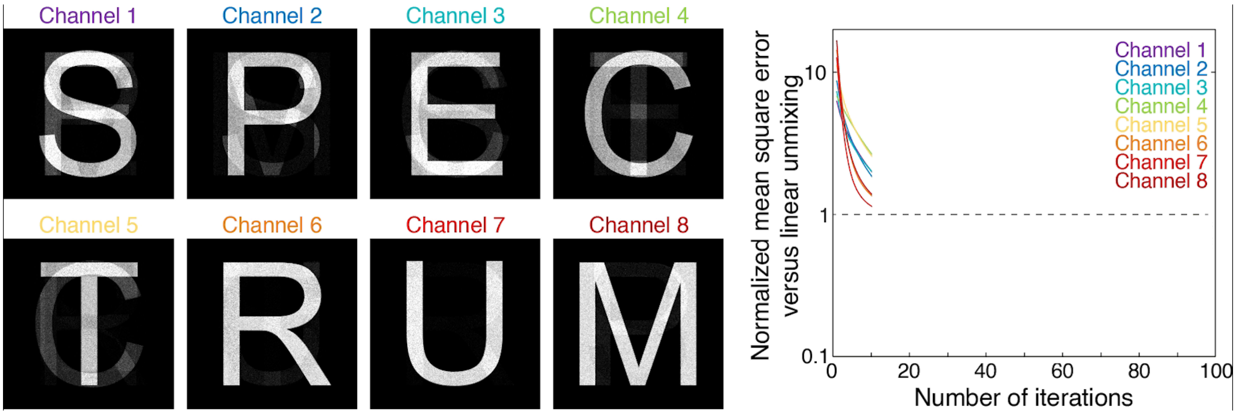


**Supplementary Video 1: RLSU outperforms linear unmixing with simulated multispectral datasets.**
Relates to Supplementary Figure 6. Eight simulated ground truth objects were spectrally mixed and Poisson shot noise was incorporated to simulate eight-channel multispectral fluorescence microscopy data. These images were unmixed using RLSU. Left panel shows how the data progressively gets unmixed as the number of iteration increases. Right panel shows the normalised mean squared error of unmixing for RLSU iterations versus linear unmixing for the data presented on the left panel Unmixing mean squared error shown for all 8 channels in RLSU in lines colour-coded by channel, whilst the linear unmixing result is indicated by dashed horizontal black line.

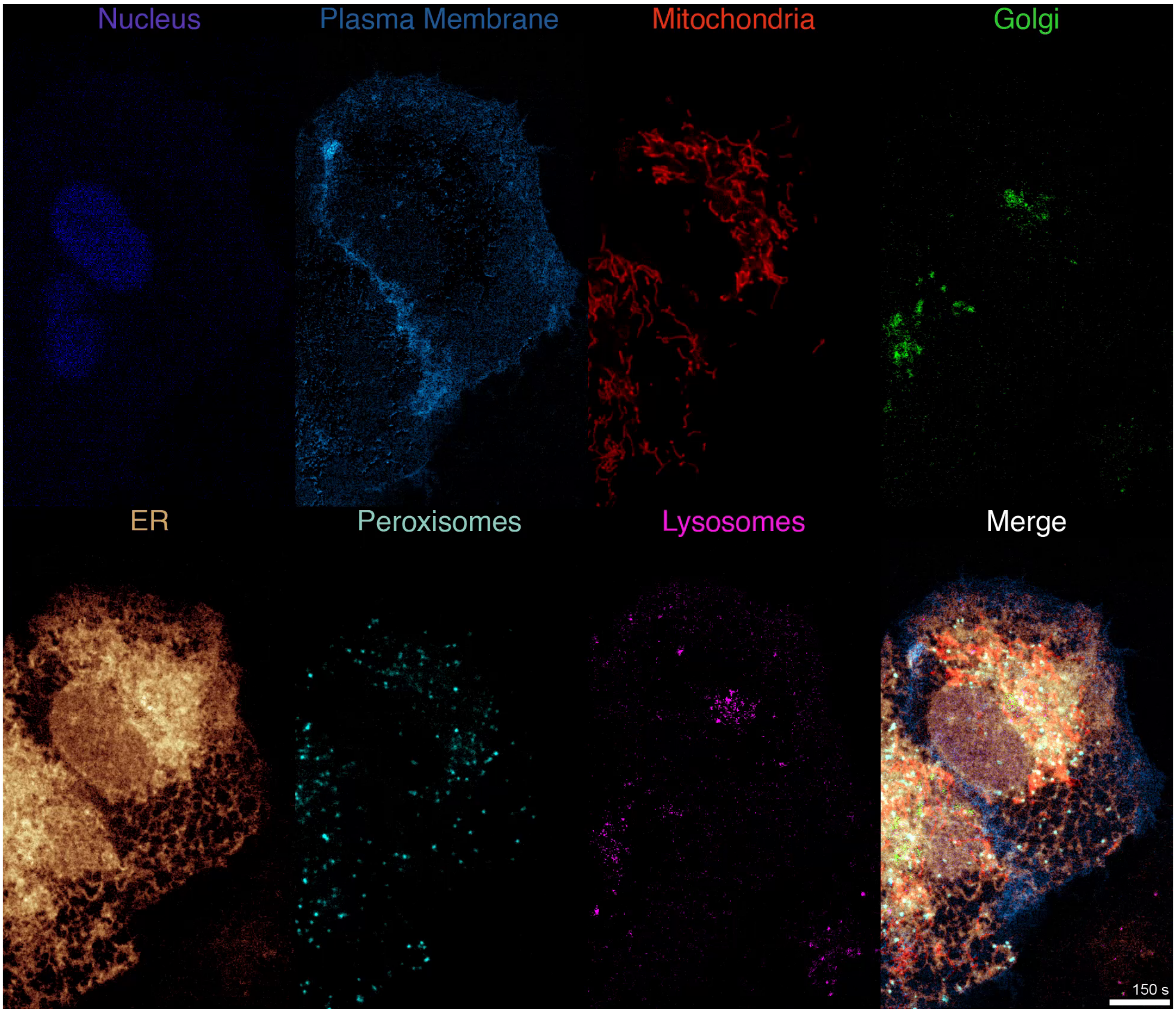


**Supplementary Video 2: Multispectral spinning disk confocal imaging of organelle trafficking.**
Relates to Figure 3d. U2OS cells transfected with the ColorfulCell plasmid were incubated with LysoTracker Yellow before imaging on a confocal spinning disk instrument equipped with the multispectral camera unit followed by RLSU unmixing. Scale bar = 10 µm.

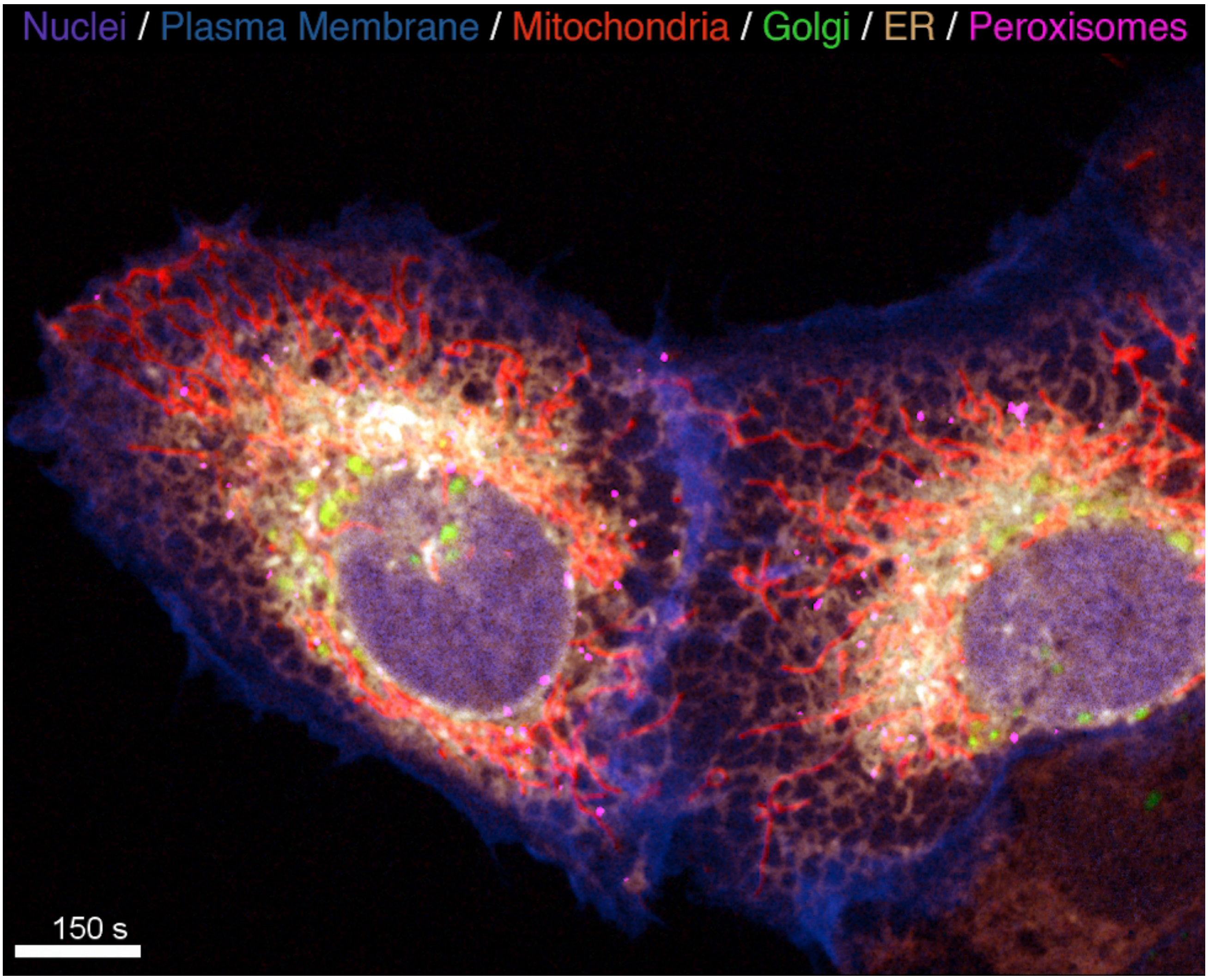


**Supplementary Video 3: Multispectral spinning disk confocal imaging.**
Relates to Figure 3e. U2OS cells transfected with the ColorfulCell plasmid were imaged on a confocal spinning disk instrument equipped with the multispectral camera unit followed by RLSU unmixing. Scale bar = 10 µm.

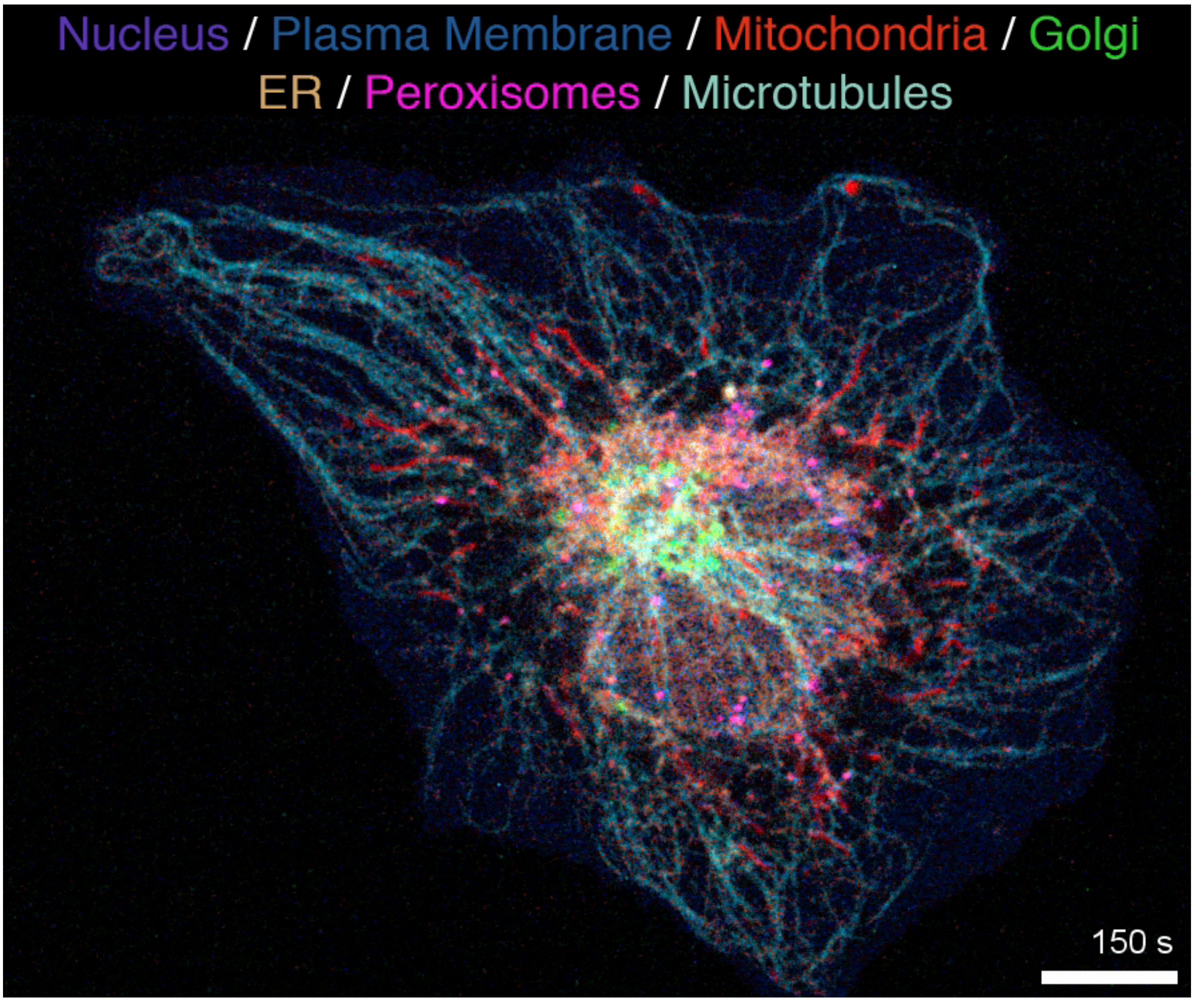


**Supplementary Video 4: Multispectral spinning disk confocal imaging of microtubules and various organelles.**
Relates to Figure 3f. U2OS cells transfected with the ColorfulCell plasmid and incubated with SPY555-tubulin were imaged on a confocal spinning disk instrument equipped with the multispectral camera unit followed by RLSU unmixing. Scale bar = 10 µm.

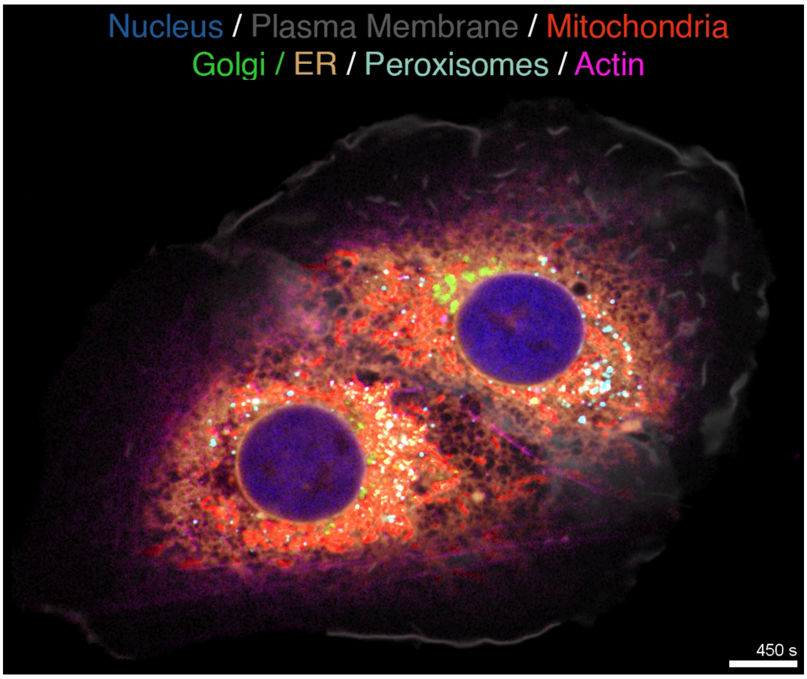


**Supplementary Video 5: Multispectral spinning disk confocal imaging of actin and various organelles.**
Relates to Figure 3g. U2OS cells transfected with the ColorfulCell plasmid and incubated with SPY555-actin were imaged on a confocal spinning disk instrument equipped with the multispectral camera unit followed by RLSU unmixing. Scale bar = 10 µm.

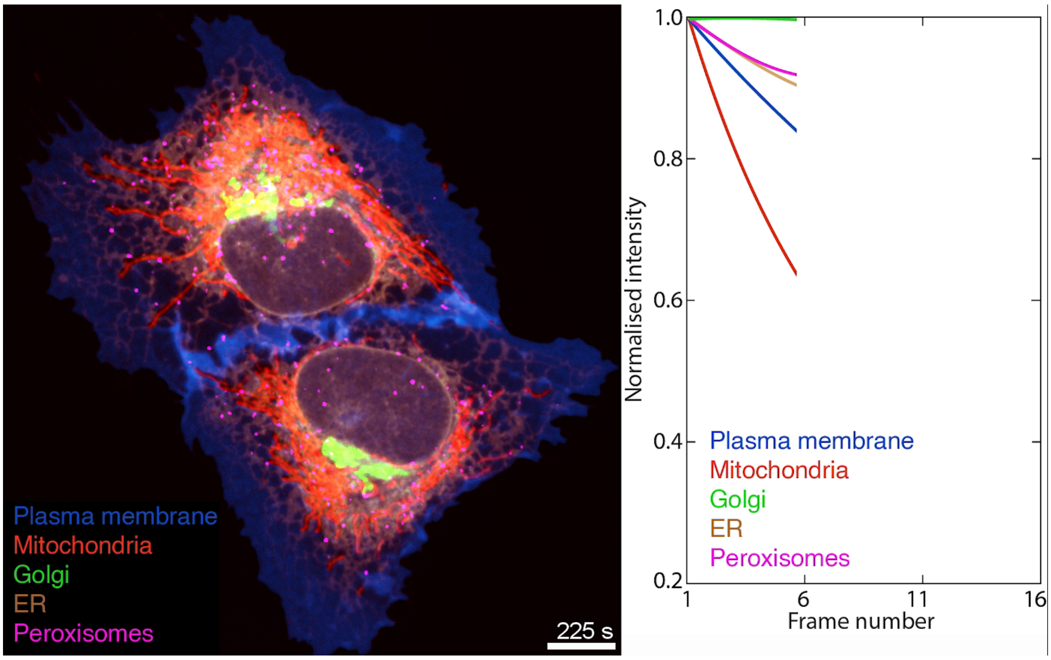


**Supplementary Video 6: Volumetric multispectral spinning disk confocal microscopy exhibits significant photobleaching.**
Relates to Supplementary Figure 16. Left panel: U2OS cells transfected with the ColorfulCell plasmid were imaged on a confocal spinning disk instrument equipped with the multispectral camera unit followed by RLSU unmixing. Images represent orthogonal maximum intensity projections of confocal volumes (51 planes ; 200 nm increment). Right panel: normalised intensity over time of left panel, showing significant photobleaching over time. Scale bar = 10 µm.

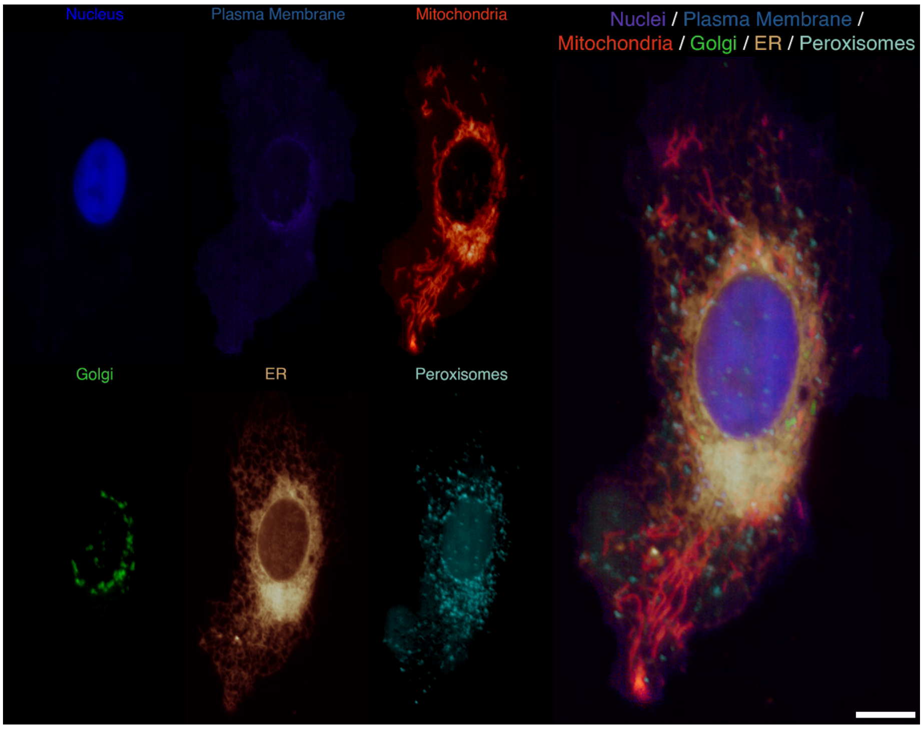


**Supplementary Video 7: Multispectral volumetric light sheet imaging.**
Relates to Figure 4. U2OS cells transfected with the ColorfulCell plasmid were imaged on an oblique plane light sheet instrument equipped with the multispectral camera unit followed by RLSU unmixing. movie show full cell volume (201 planes). Scale bar = 10 µm.

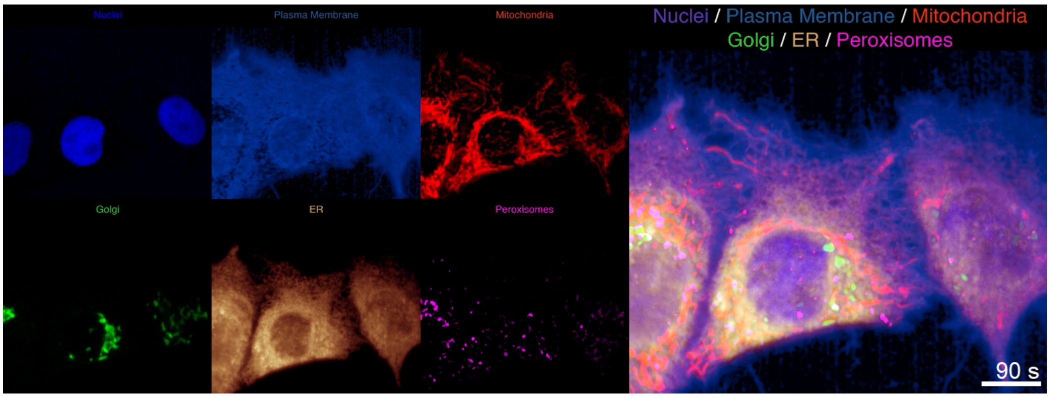


**Supplementary Video 8: Multispectral volumetric light sheet imaging timelapse.**
Relates to Figure 4. HeLa cells transfected with the ColorfulCell plasmid were imaged on an oblique plane light sheet instrument equipped with the multispectral camera unit followed by RLSU unmixing. Movie shows maximum intensity projections of full cell volumes (81 planes) over time. Scale bar = 10 µm.

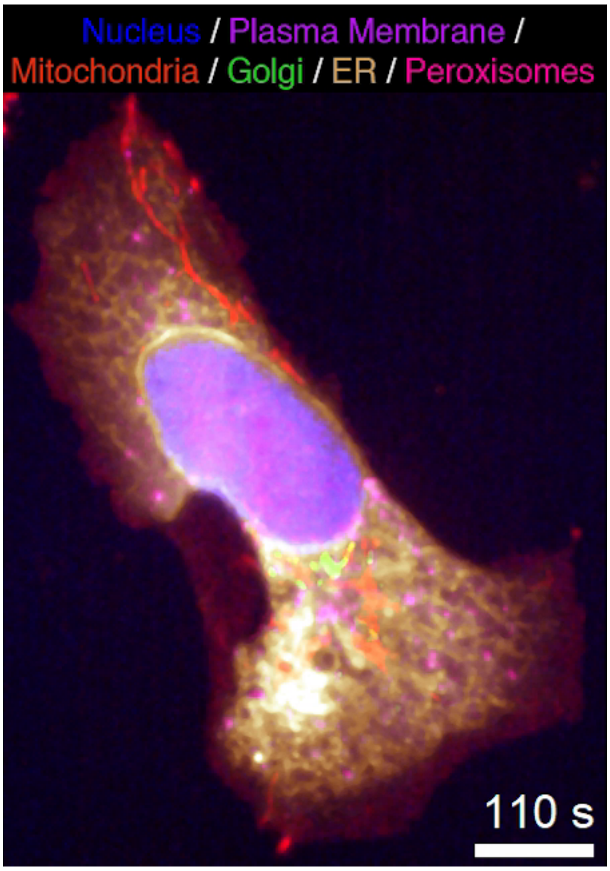


**Supplementary Video 9: Long term, volumetric imaging by multispectral oblique plane light sheet microscopy.**
Relates to Figure 4. U2OS cells transfected with the ColorfulCell plasmid were imaged on a oblique plane light sheet instrument equipped with the multispectral camera unit followed by RLSU unmixing. Movie shows maximum intensity projections of full cell volumes (101 planes) over time. Note the negligible photobleaching observed over the 200 timepoint timelapse (1 volume every 10 s) due to the gentle excitation strategy of the oblique plane light sheet microscope. Scale bar = 10 µm.

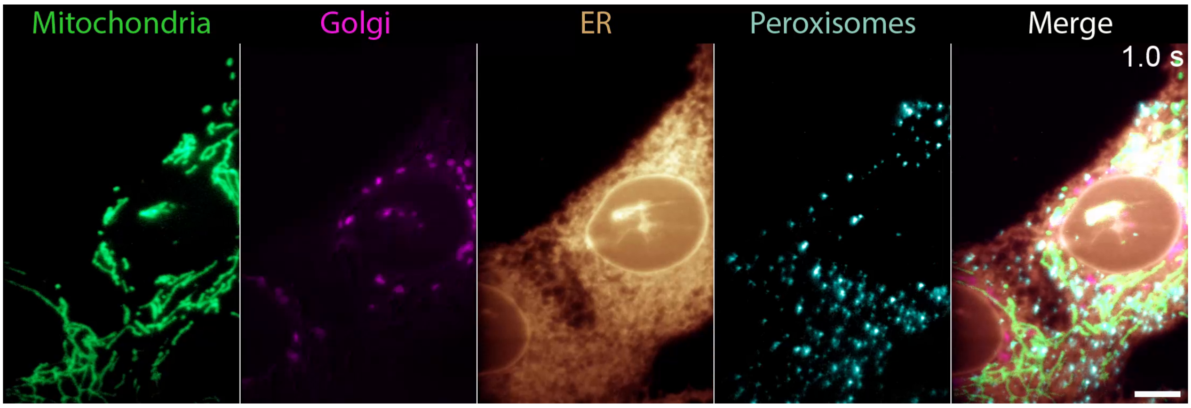


**Supplementary Video 10: Fast multispectral live-cell imaging by projection light sheet microscopy.**
Relates to Figure 5. U2OS cells transfected with the ColorfulCell plasmid were imaged by projection imaging on an oblique plane light sheet instrument equipped with the multispectral camera unit followed by RLSU unmixing. Only four markers were imaged (mAzamiGreen-mitochondria, Citrine-Golgi, mCherry-endoplasmic reticulum and iRFP670-peroxisomes) as 405 nm laser was not turned on in this specific experiment. Note how the fast framerate reached by the instrument (10 projections of the entire cells per second) allows one to simultaneously observe fast dyanmics (ER tubules) and slow dynamics (peroxisome Brownian motion). Scale bar = 10 µm.

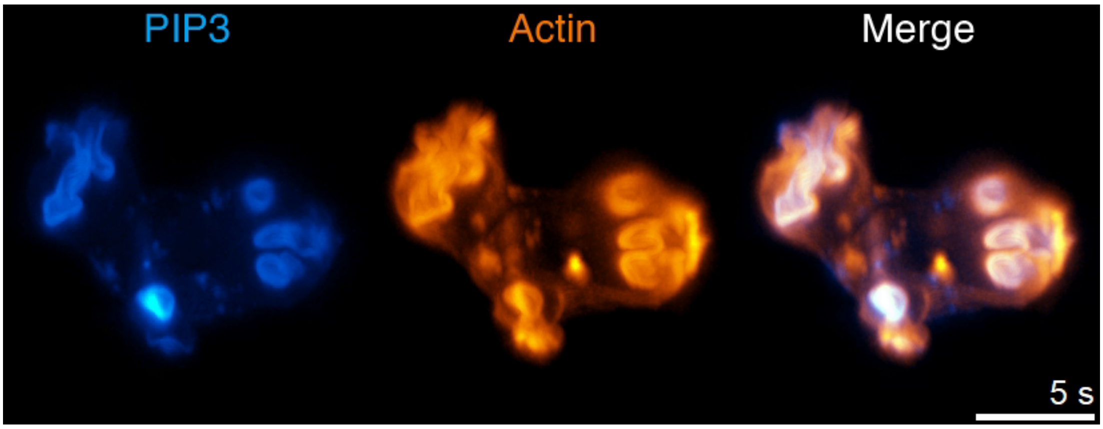


**Supplementary Video 11: Fast volumetric imaging of live *Dictyostelium discoideum* cells.**
Relates to Supplementary Figure 18. Live *Dictyostelium discoideum* cell expressing EGFP-fused to a PIP3 reporter and LifeAct-mCherry were imaged using the multispectral OPM at 2 volumes per second. Images correspond to maximum intensity projections. Note that this volumetric aquisition speed is enough to resolve macropinocytic cup closure. Scale bar = 10 µm.

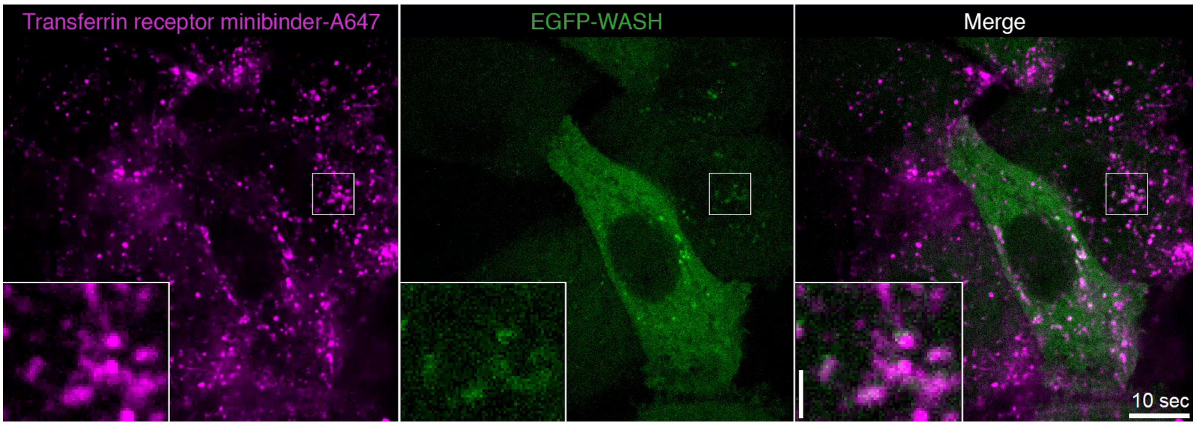


**Supplementary Video 12: Validation that de novo designed binders reach early-sorting endosomes in live cells.**
Relates to Supplementary Figure 20. NIH 3T3 cells stably expressing EGFP-WASH and incubated with Alexa Fluor 647-labelled TfR minibinder were imaged live by spinning disk confocal microscopy. Images correspond to single confocal planes. Scale bar = 10 µm.

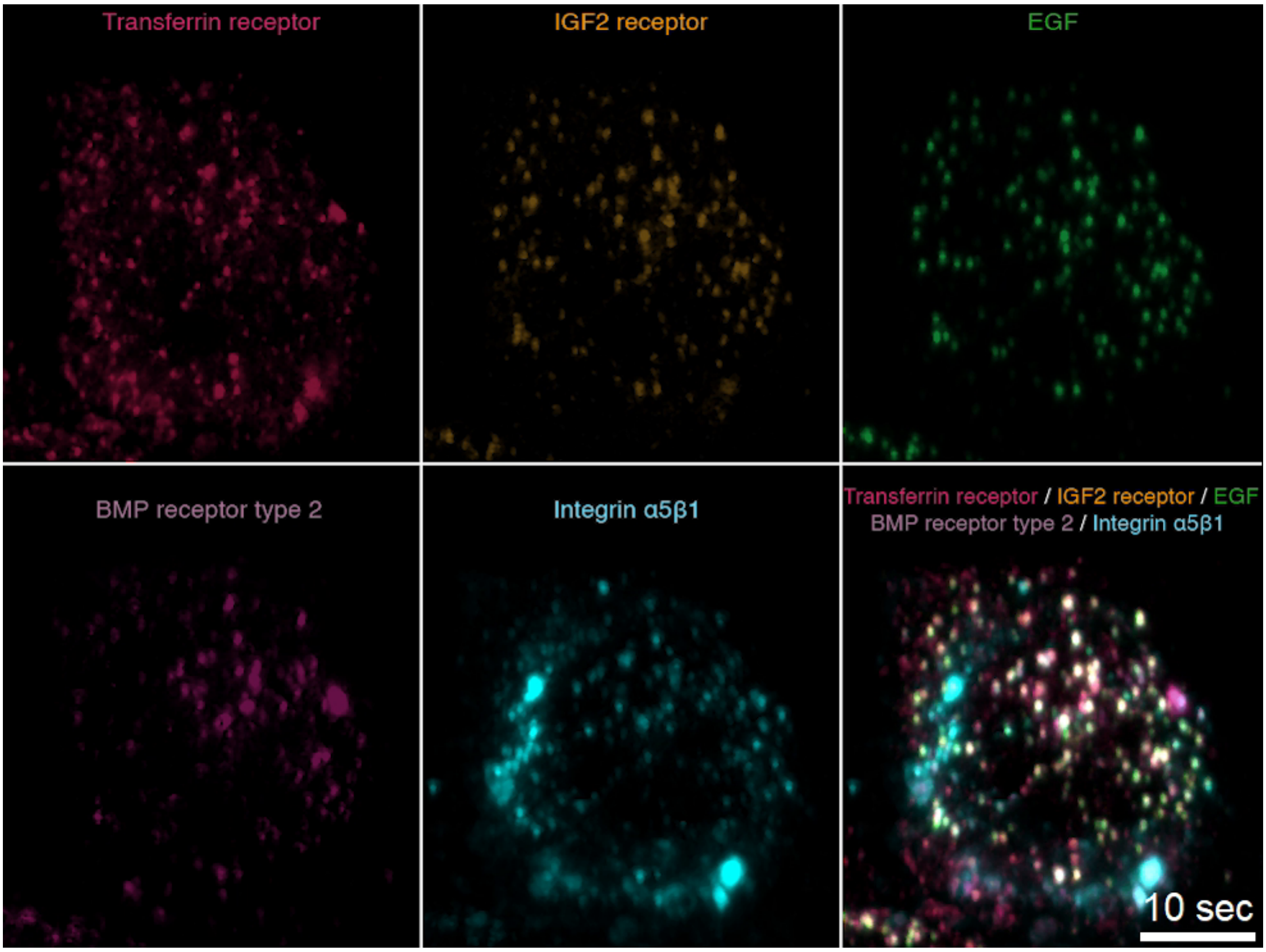


**Supplementary Video 13: Multispectral live-cell light sheet imaging of intracellular trafficking using de-novo designed receptor minibinders.**
Relates to Figure 6a. TagBFP-NLS-expressing HeLa Kyoto cell loaded with four fluorescently-labelled, computationally designed minibinders against cell surface receptors (TfR, IGF2R, BMPR2 & I$\alpha5\beta1$) and one fluorescently-labelled receptor ligand (EGF) were imaged live by multispectral oblique plane light sheet microscopy. Images correspond to maximum intensity projections of the entire cell volume (67 planes), and volumetric imaging is performed at 1 Hz. For convenience, the signal of TagBFP-NLS is not shown. Note that the ratio of the different minibinders and EGF is not the same in all endosomes, suggesting the different receptors are being sorted away from one another. Scale bar = 10 µm.

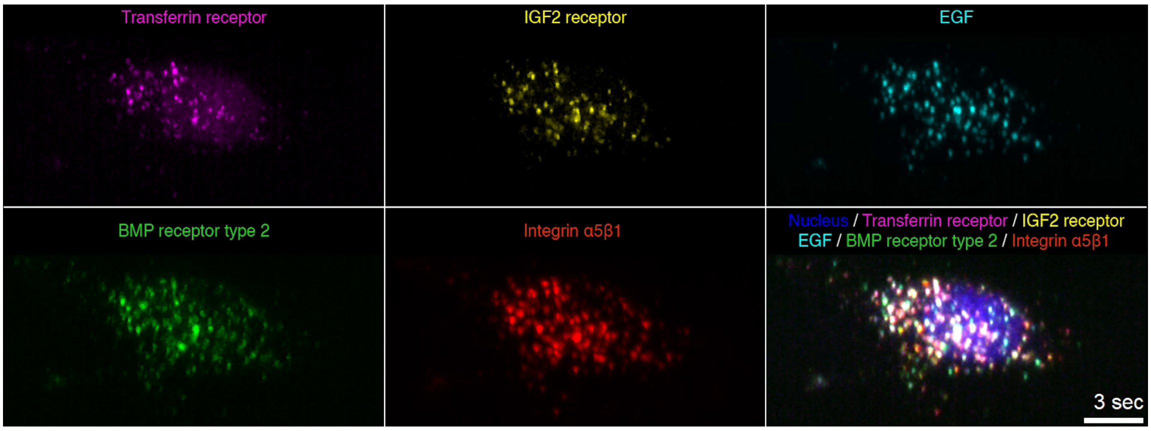


**Supplementary Video 14: Fast multispectral live-cell light sheet imaging of intracellular trafficking using de-novo designed receptor minibinders.**
Relates to Figure 6. TagBFP-NLS-expressing HeLa Kyoto cell loaded with four fluorescently-labelled, computationally designed minibinders against cell surface receptors (TfR, IGF2R, BMPR2 & I$\alpha$5$\beta$1) and one fluorescently-labelled receptor ligand (EGF) were imaged live by multispectral oblique plane light sheet microscopy. Images correspond to maximum intensity projections of the entire cell volume (47 planes), and volumetric imaging is performed at 3 Hz. Note that the fast aquisition speed allows tracking of all endosomes in the cell in 3D without motion blur even for fast-moving organelles. Also note the absence of colour misregistration caused by delays in acquiring colour channels in typical sequential aquisition schemes. Scale bar = 10 µm.
